# Diauxic lags explain unexpected coexistence in multi-resource environments

**DOI:** 10.1101/2021.08.09.455603

**Authors:** William Bloxham, Hyunseok Lee, Jeff Gore

**Affiliations:** Physics of Living Systems, Department of Physics, Massachusetts Institute of Technology, Cambridge MA, USA

## Abstract

How the coexistence of species is affected by the presence of multiple resources is a major question in microbial ecology. We experimentally demonstrate that differences in diauxic lags, which occur as species deplete their own environments and adapt their metabolisms, allow slow-growing microbes to stably coexist with faster-growing species in multi-resource environments despite being excluded in single-resource environments. In our focal example, an *Acinetobacter* species (Aci2) competitively excludes *Pseudomonas aurantiaca* (Pa) on alanine and on glutamate. However, they coexist on the combination of both resources. Experiments reveal that Aci2 grows faster but Pa has shorter diauxic lags. We establish a tradeoff between Aci2’s fast growth and Pa’s short lags as their mechanism for coexistence. We model this tradeoff to accurately predict how environmental changes affect community composition. We extend our work by surveying a large set of competitions and observe coexistence nearly three times as frequently when the slow-grower is the fast-switcher. Our work illustrates a potentially common mechanism for the emergence of multi-resource coexistence despite single-resource competitive exclusions.

## Introduction

Explaining the rich biodiversity observed in nature and the mechanisms by which species coexist are central questions in microbial ecology^1–4^. Microbes exist in complex environments and often have multiple resources to choose between. It is therefore important to study coexistence from the perspective of how microbes interact with their environments, including which resources they consume and how they consume them^5–11^. Previous research has suggested communities grown on a single carbon resource can be used to predict multi-resource communities^9,12–15^. There are, however, frequent exceptions, such as a species being excluded in single-resource communities but coexisting in multi-resource communities^9,16^. These exceptions suggest dynamics specific to multi-resource environments must be considered to fully understand community assembly.

Diauxie is a commonly observed phenomenon for microbes growing in a multi-resource environment^16–20^. Diauxie was first observed in yeast in 1900^21^ and then studied in greater detail in *E. coli* by Monod beginning in 1940^22,23^. When grown in a media containing two sugars, *E. coli* first displays exponential growth, then exhibits a period of little or no growth, and then resumes growing. In these cases, *E. coli* is consuming one sugar at a time^23^. For example, when presented with glucose and xylose, *E. coli* consumes only glucose until glucose runs out and then switches to xylose. In between growth on glucose and growth on xylose are approximately two hours in which *E. coli* displays no growth^19^. This period is known as a diauxic lag and occurs when a microbe needs to reconfigure its metabolism before continuing to grow^24–26^. Depending on the microbe and the combination of resources diauxic lags can last anywhere from a few minutes to several hours^19,22,23,27,28^.

Diauxie is often contrasted to co-utilization, a metabolic strategy in which a microbe consumes multiple resources at the same time. However, co-utilizing microbes experience the same resource depletions as specializing species and also need to readjust their metabolisms^24,29^. Lag phases associated with this readjustment occur, although they may be shorter^25,30,31^, and produce the growth-lag-growth pattern characteristic of diauxie^20,24,32^. Due to these similarities, we refer to a lag phase associated with a metabolic readjustment as a diauxic lag regardless of whether the microbe was previously co-utilizing.

Diauxic lags can have a significant impact on interspecies competition. Two microbes often have very different lags after the same resource depletion^25,33,34^. Some experiments have even shown correlations between growth rate and lag time such that slow-growers are more likely to be fast-switchers^28,31,32,35^. In co-culture, this tradeoff can create a back-and-forth in which a fast-grower outpaces a slow-grower before a resource depletion, but the slow-grower, by being the fast-switcher, catches up to and even overtakes the fast-grower after the resource depletion^25,32,33^. Alternating resource supplies can also drive species to evolve shorter diauxic lags, often producing distinct slow-switcher and fast-switcher phenotypes^33,34,36^. These results suggest diauxic lags could be a common source of stable coexistence.

We provide an experimental demonstration of a tradeoff between growth rate and diauxic lag time producing stable coexistence between two species. We investigate the case of an *Acinetobacter* species (Aci2) and *Pseudomonas aurantiaca* (Pa) growing on alanine and glutamate. Despite being the slow-grower and being competitively excluded by Aci2 in both single-resource environments, Pa stably coexists with Aci2 in the two-resource environment. We discover that, while Aci2 is the fast-grower, Pa is the fast-switcher. Through modeling and additional experiments we confirm the tradeoff between growth rate and diauxic lag time as the mechanism for coexistence. We extend our work by surveying additional species and resources and see that coexistence is nearly three times as likely to occur when the slow-grower is the fast-switcher. Our research establishes a mechanism for the otherwise unexpected coexistence of two species based on tradeoffs between growth rate and diauxic lag time and highlights the importance of diauxic lags on interspecies competition in multi-resource environments.

## Results

### Pa unexpectedly coexists with Aci2 in a multi-resource environment

To understand community assembly in multi-resource environments, we cocultured an *Acinetobacter* species (Aci2) and *Pseudomonas aurantiaca* (Pa) in three environments under batch culture with daily dilution. When Aci2 and Pa were cocultured in an environment containing alanine as the only carbon source, Aci2 competitively excluded Pa (Fig. 1*A*). When cocultured with glutamate as the only carbon source, Aci2 again competitively excluded Pa (Fig. 1*B*). If two-resource communities were a simple sum of single-resource communities^9,14,15^, one would expect Aci2 to competitively exclude Pa in an environment containing alanine and glutamate. Contrary to this expectation, when Aci2 and Pa were cocultured on an equal mix of alanine and glutamate, they stably coexisted (Fig. 1*C*). Across eight replicates Pa had a population fraction between 0.34 and 0.42 that was independent of the starting fraction (Fig. 1*C*). This was an intriguing result with the potential to illuminate mechanisms for coexistence unique to multi-resource environments.

**Fig. 1.**
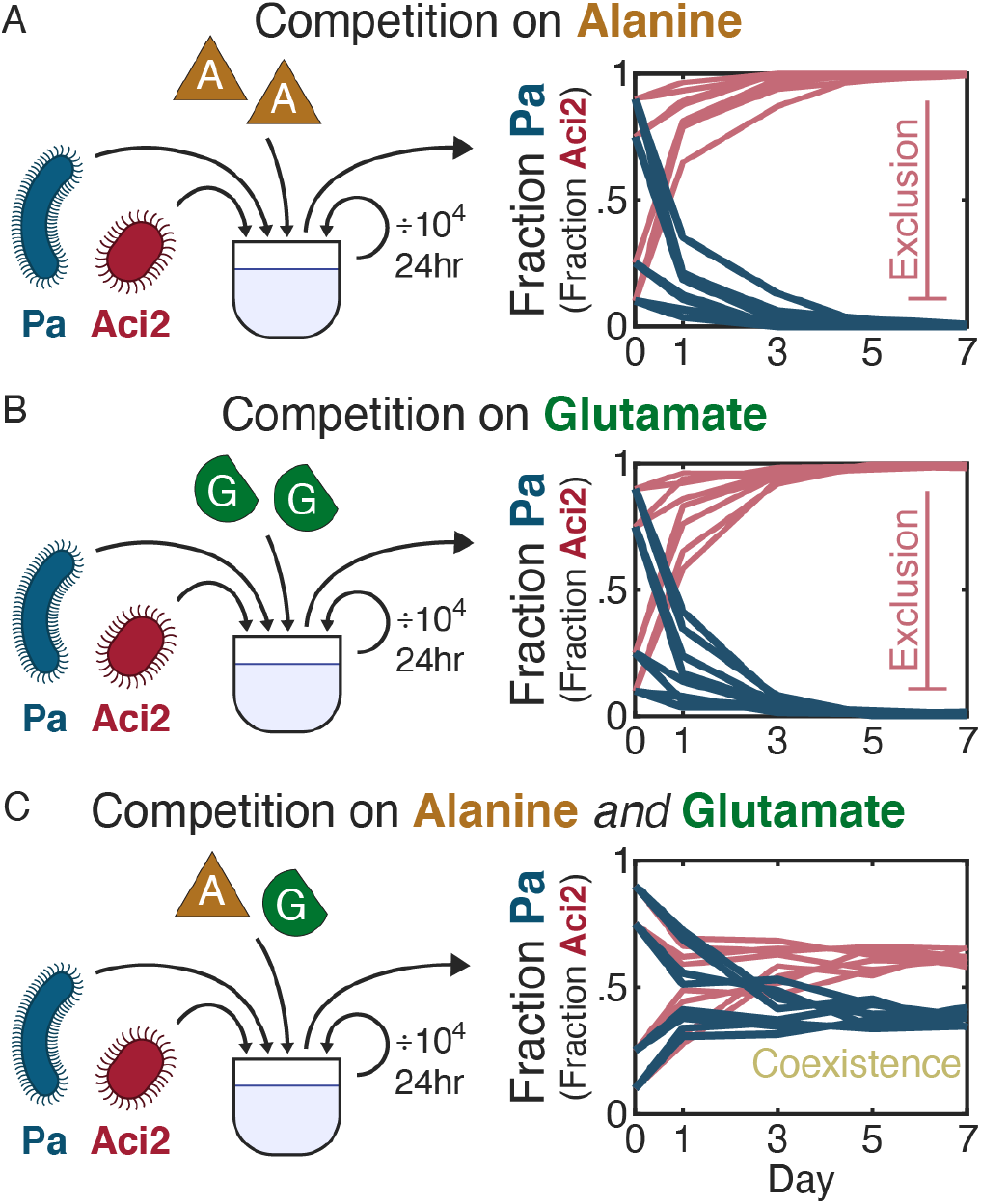
Pa stably coexists with Aci2 when co-cultured on alanine and glutamate despite being competitively excluded when co-cultured on either resource alone. (*A*) Pa (*Pseudomonas aurantiaca*) and Aci2 (*Acinetobacter* sp.) were co-cultured with a 10^4^ dilution every 24 hours and 0.1%w/v alanine supplied as the only carbon source. Two replicates were performed for each of four initial Pa fractions: 0.1, 0.25, 0.75, and 0.9. Population sizes were measured via colony counting after the first, third, fifth, and seventh days. With only alanine supplied, Aci2 competitively excluded Pa. (*B*) An identical experiment was performed with 0.1%w/v glutamate supplied as the only carbon source. Aci2 again competitively excluded Pa. (*C*) A third experiment was performed with both 0.05%w/v alanine and 0.05%w/v glutamate supplied. Pa coexisted with Aci2 and reached a population fraction between 0.34 and 0.42 independent of initial starting fraction. Two-resource coexistence was unexpected given the single-resource competitive exclusions.

We began our investigation into the coexistence of Aci2 and Pa by ruling out one of the simplest explanations. Species generally grow faster as the number of available resources increases^37,38^. We hypothesized that Aci2 was the fast-grower in both single-resource environments while Pa was the fast-grower in the two-resource environment. We measured growth rates and saw that Aci2 was the fast-grower in both single-resource environments (Supp. Fig. 1), which explained why Aci2 excluded Pa in those cocultures. However, we also saw that Aci2 was the fast-grower in the two-resource environment, with a growth rate 31% +/− 2% faster than Pa’s (0.88 +/− 0.01 hr^−1^ vs 0.67 +/− 0.01 hr^−1^) (Fig. 2*A*). That Aci2 was the consistent fast-grower made it even more surprising that Pa had coexisted with Aci2 in the alanine-glutamate environment.

**Fig. 2.**
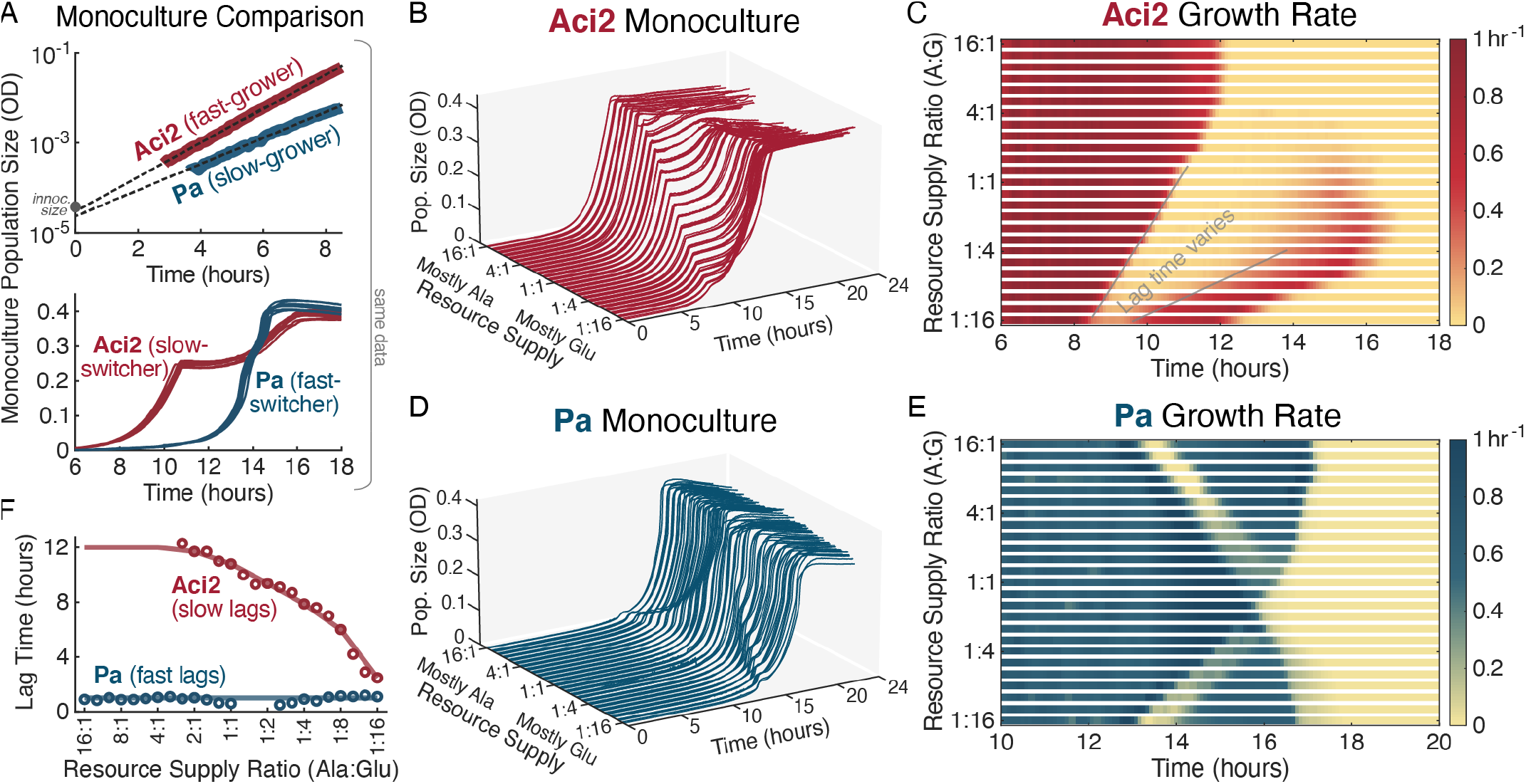
Monoculture growth dynamics reveal Aci2 is a fast-grower but slow-switcher whereas Pa is a slow-grower but fast-switcher. (A) Aci2 and Pa were both grown in monoculture in the two-resource environment. The same data is shown in both plots. The top plot shows the average of eight replicates on a log scale, while the bottom plot shows each of the replicates on a linear scale. Overlain on the top plot are growth rate fits of *g*_Aci2_ = 0.88hr^−1^ and *g*_Pa_ = 0.67hr^−1^. (*B*) Aci2 was grown with alanine and glutamate supplied at 26 different ratios, ranging from 1:16 to 16:1 with four replicates at most supply ratios. The total resource supply was kept constant at 0.1%w/v. The population size at which Aci2’s diauxic shift occurred is linearly correlated to the alanine supply concentration (see Supp. Fig. 2 for detail). (*C*) Instantaneous growth rates were extracted from the Aci2 monoculture data and averaged across replicates. (*D*–*E*) The same monoculture experiments were performed for Pa. The appearance of Pa having a long initial lag in *D* is primarily due to the growth that it needs to accomplish before its population size becomes significant on a linear scale. (*F*) Diauxic lag times were fit from the monoculture data (see Supp. Fig. 3) and are plotted as circles. Lag times could not be fit in the with too little growth on the remaining resource. The diauxic lag times used in the modeling at shown as lines through the data: for Aci2 this is a linear interpolation through the whole-number resource supply ratios with a value of 12 hours used for ratios greater than 4:1; for Pa this is a constant value of 1 hour.

### Aci2 is the fast-grower, but Pa is the fast-switcher

The growth rate experiments did, however, contain a hint towards another explanation. Aci2 had a several-hour diauxic lag around half-saturation whereas Pa had a short diauxic lag lasting no more than an hour (Fig. 2*A*). Aci2 was the fast-grower but slow-switcher, whereas Pa was the slow-grower but fast-switcher. We hypothesized that a tradeoff between these properties could be the source of coexistence.

To test whether Pa’s short lag could explain the coexistence of Aci2 and Pa, we first needed a more detailed understanding of each species. To determine which resource each species consumed first, we varied the resource supply ratio in our monoculture experiments. The population density at which Aci2 displayed its diauxic shift was proportional to the alanine supply concentration (Fig. 2*B* and Supp. Fig. 3), such that the amount Aci2 could grow before its diauxic shift directly matched the amount of alanine available, indicating that Aci2 first consumes almost entirely alanine.

In contrast to Aci2’s initial single-resource consumption, Pa’s dynamics showed clear signs of co-utilization. At a low supply fraction of glutamate, Pa had a diauxic lag that occurred at larger population sizes with increasing glutamate supply. But at a low alanine supply, Pa’s lag occurred at larger population sizes with increasing alanine supply (Fig. 2*D* and 2*E* and Supp. Fig. 3). As a supply ratio of 2:3 alanine:glutamate was approached, the population density at which Pa’s diauxic shift occurred converged towards the saturation density, such that near a 2:3 supply ratio Pa simply saturated with no diauxic shift. These observations indicated that Pa consumed alanine and glutamate in a fixed ratio of approximately 2:3 when growing in the two-resource environment.

These monoculture experiments also yielded additional information about the diauxic lag times of each species. Notably, Aci2’s diauxic lag time varied considerably with resource supply ratio. When mostly glutamate was supplied Aci2’s lag time was the shortest (~2 hours), and as the supply of alanine increased Aci2’s lag time became longer (up to ~12 hours) (Fig. 2*C* and 2*F* and Supp. Fig. 3). Examples of variable diauxic lag times have been studied and seen to be the result of sensing relative resource supplies^26^ and adaptive bet-hedging^30,39^. Pa’s lag time appeared constant at around 1 hour and was therefore always shorter than Aci2’s (Fig. 2*E* and 2*F* and Supp. Fig. 3). That a slow-growing and previously co-utilizing species would have a shorter lag makes intuitive sense and aligns with previous results^25,26^. That Pa would still briefly stop growing could be due to not expressing enzymes to convert between glutamate and alpha-ketoglutarate prior to resource depletion, thus preventing efficient energy production from glutamate nor synthesis of glutamate-derived amino acids from alanine until Pa’s lag has finished. Combined with a confirmation that neither species’ initial growth rate varied with resource supply ratio (Supp. Fig. 4), these observations established Aci2 as the fast-grower but slow-switcher and Pa the slow-grower but fast-switcher at all resource supply ratios.

### Simple growth-lag model reproduces monoculture dynamics

With an understanding of each species’ growth dynamics, we developed a growth-lag model in which the only tradeoff would be between growth rate and diauxic lag time. We used a single exponential growth rate for each species that applied to all environments to eliminate any tradeoff between two-resource and single-resource growth rates (Fig. 3*A*). The model used the experimentally determined two-resource growth rates: 0.88 hr^−1^ for Aci2 and 0.67 hr ^−1^ for Pa (Fig. 2*A*). Lag times were interpolated from experimentally determined values (Fig. 2*F*), including the dependence of Aci2’s lag time on resource supply ratios.

**Fig. 3.**
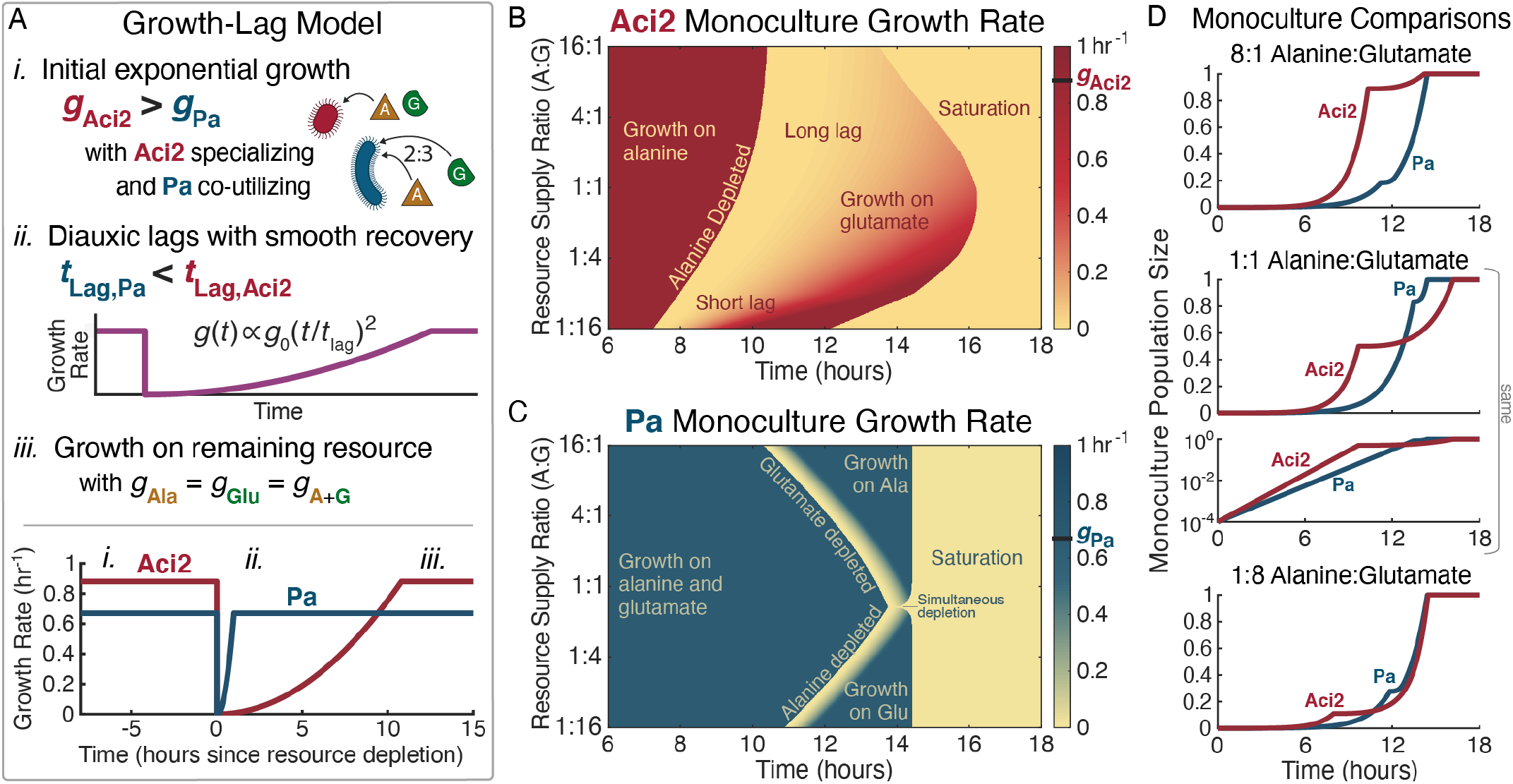
Growth-lag model reproduces monoculture growth dynamics. (*A*) A simple model of growth and lag phases. (*i*.) Species initially grow exponentially with Aci2 consuming only alanine and Pa consuming alanine and glutamate in a fixed 2:3 ratio. (*ii*.) When a resource is depleted, both species experience a diauxic lag with a smooth recovery. Aci2’s lag time is long and varies with resource supply (using values in Fig. 2*F*). Pa’s lag is short and constant at 1 hour. Growth rates during the lag phase are proportional to the square of the time since the resource depletion. (This quadratic model was a better fit when extracting growth rates than other recovery shapes considered. See Supp. Fig. 4 for comparison.) The plot at the bottom of *A* compares the growth rate recoveries of Aci2 and Pa using Aci2’s lag time from the 1:1 alanine:glutamate condition. (*iii*.) Once recovered from its diauxic lag, each species grows on the remaining resource at the same rate as its initial growth. (*B*) Model predictions for Aci2’s monoculture growth for different resource supply ratios when started at a 104 dilution from carrying capacity. Each row of pixels is the calculation for a separate condition. (*C*) The equivalent predictions for Pa’s growth. (*D*) Comparison of the monoculture growth calculations at 8:1, 1:1, and 1:8 resource supply ratios. The appearance of initial lag is due to the growth that needs to happen before population sizes become significant on a linear scale, as can be seen in the log-scale plot for the 1:1 condition.

The growth-lag model reproduced the key features of Aci2’s monoculture dynamics (Fig. 3*B* in comparison to Fig. 2*C*). Aci2 initially consumed only alanine, so in both model and experiment Aci2’s diauxic shift occurred later the more alanine was supplied. Saturation, meanwhile, occurred latest at roughly equal resource supply ratios. Quick saturation at low alanine supply fractions was the result of Aci2’s shorter diauxic lag. Quick saturation at high alanine supply fractions was achieved despite Aci2’s long lag because Aci2 did not need to significantly recover its growth rate to complete the small amount of remaining growth. The reproduction of these features indicates that our simple model was accurately capturing Aci2’s growth and resource consumption behavior.

The growth-lag model for Pa contained the same elements as for Aci2, but defining Pa as initially co-utilizing alanine and glutamate and giving Pa a shorter, constant lag produced distinct dynamics. These dynamics were again a close match between model and experiment (Fig. 3*C* in comparison to Fig. 2*E*). The onset of Pa’s diauxic shift occurred as a result of alanine running out at small alanine supply and as a result of glutamate running out at small glutamate supply. Pa’s short and constant diauxic lag meant it saturated at the same time for most resource supply ratios. When the supply ratio matched Pa’s consumption ratio, alanine and glutamate ran out at the same time so Pa had no diauxic shift and saturated slightly earlier. These features of Pa’s growth dynamics all agreed between model and experiment. The growth-lag model therefore reproduced the distinct growth and resource-consumption dynamics of both Aci2 and Pa.

### Tradeoff between growth rate and lag time is sufficient for coexistence

With a growth-lag model developed and shown to accurately reproduce the monoculture dynamics, we were ready to test whether the tradeoff between growth rate and lag time was sufficient for the experimentally observed coexistence. In competition, the two species maintained the same resource consumption dynamics as in monoculture but now competed for the same resource pool. Each time the species depleted both resources, their populations were divided by the dilution factor and resource concentrations were returned to the supply concentrations (Fig. 4*A*). We used this model to predict the species’ population fractions over seven dilution cycles (as was done experimentally in Fig. 1*C*).

**Fig. 4.**
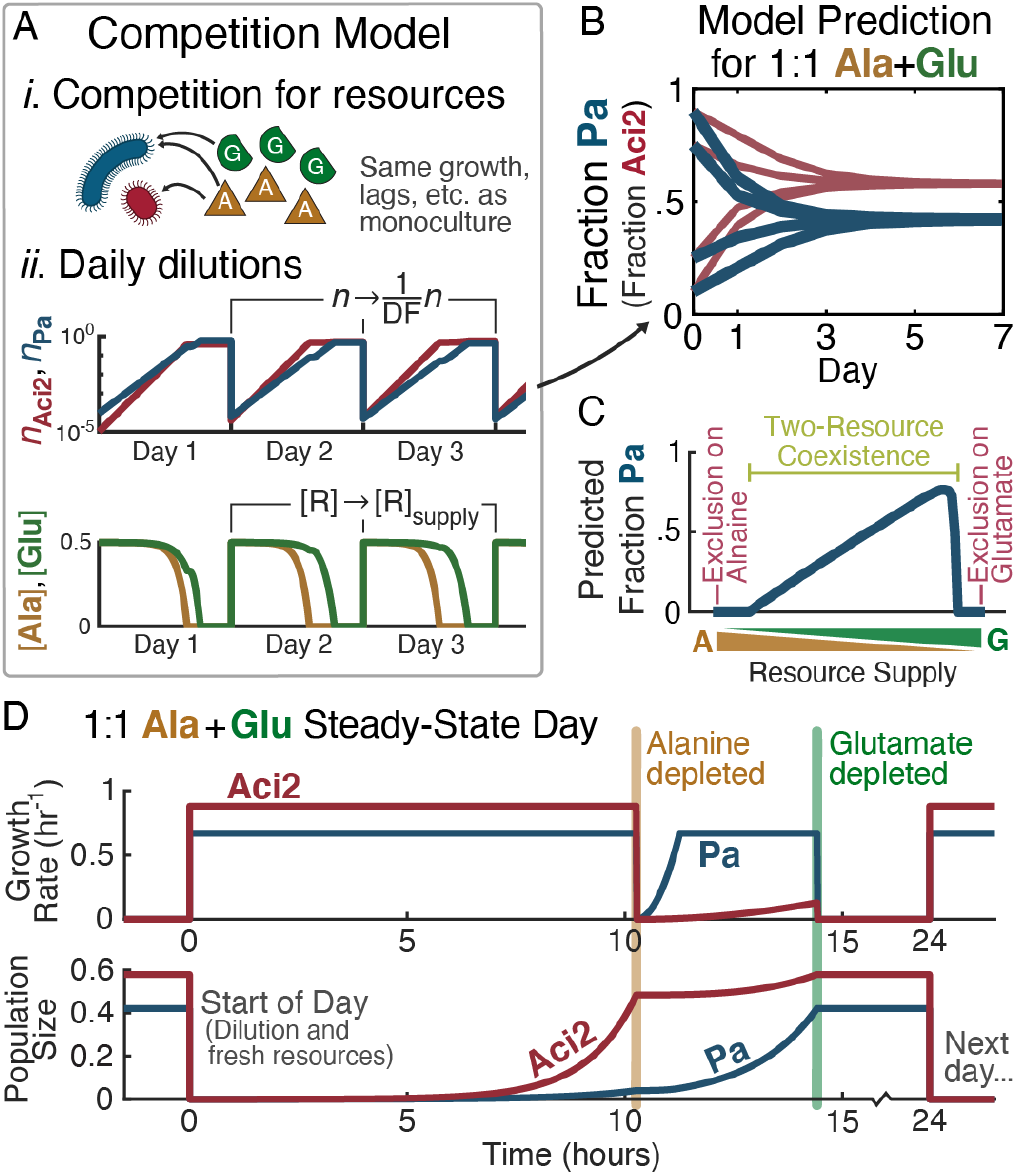
Growth-lag model uses the tradeoff between Aci2’s fast growth and Pa’s short lag to explain two-resource coexistence. (*A*) The competition model is an extension of the monoculture model. (*i*.) Species have the same growth, lags, and resource consumption, but now compete for the same resource pool. (*ii*.) After saturation, population sizes are divided by the dilution factor (104 in this figure) and resource concentrations are reset to the supply concentrations. Shown in *A* are the first three days of the 1:1 alanine:glutamate competition started with a Pa fraction of 0.9. (*B*) With parameters matching those used experimentally in Fig. 1*C*, the growth-lag model correctly predicts coexistence of Aci2 and Pa on alanine and glutamate. (*C*) The model predicts the competitive outcome will depend on the resource supply fractions with exclusion eventually occurring as the supply approaches entirely alanine or entirely glutamate. (*D*) Growth rates and population sizes over the course of one day after the 1:1 alanine:glutamate co-culture has reached steady state. Pa starts the day with a population fraction of 0.42, declines to a population fraction of 0.07 at alanine depletion, and recovers to a population fraction of 0.42 by the time glutamate is depleted.

Our model successfully reproduced stable coexistence of Aci2 and Pa on alanine and glutamate with the same steady state being reached regardless of initial starting fractions (Fig. 4*B*). At steady state, population dynamics were driven by diauxic resource consumption, with two distinct growth periods causing population fractions to vary considerably over the course of a day. Aci2 initially grew faster, causing Pa’s population fraction to decrease from 0.42 to just 0.07 over the first ~10 hours until alanine was depleted. After alanine depletion, Pa’s growth rate quickly recovered while Aci2 suffered a long lag, allowing Pa to catch back up to a population fraction of 0.42 during the ~4 hours until glutamate was also depleted (Fig 4*D*). Thus, our model balanced two distinct periods of growth to explain the experimental observation of unexpected coexistence.

### Accurate prediction of response to environmental changes validates growth-lag model

Our model predicted community composition would vary considerably with changes to resource supply ratios (Fig. 4*C*), so we decided to test whether the tradeoff between growth rate and lag time could be used to accurately predict the response to environmental changes. We chose a 54-condition parameter grid of nine resource supply ratios and six dilution factors, calculated the model’s steady-state predictions (Fig. 5*B*), and performed the coculture competitions, counting over 10^5^ colonies in total to determine precise species fractions (Fig. 5*A* and 5*C* and Supp. Fig. 5).

**Fig. 5.**
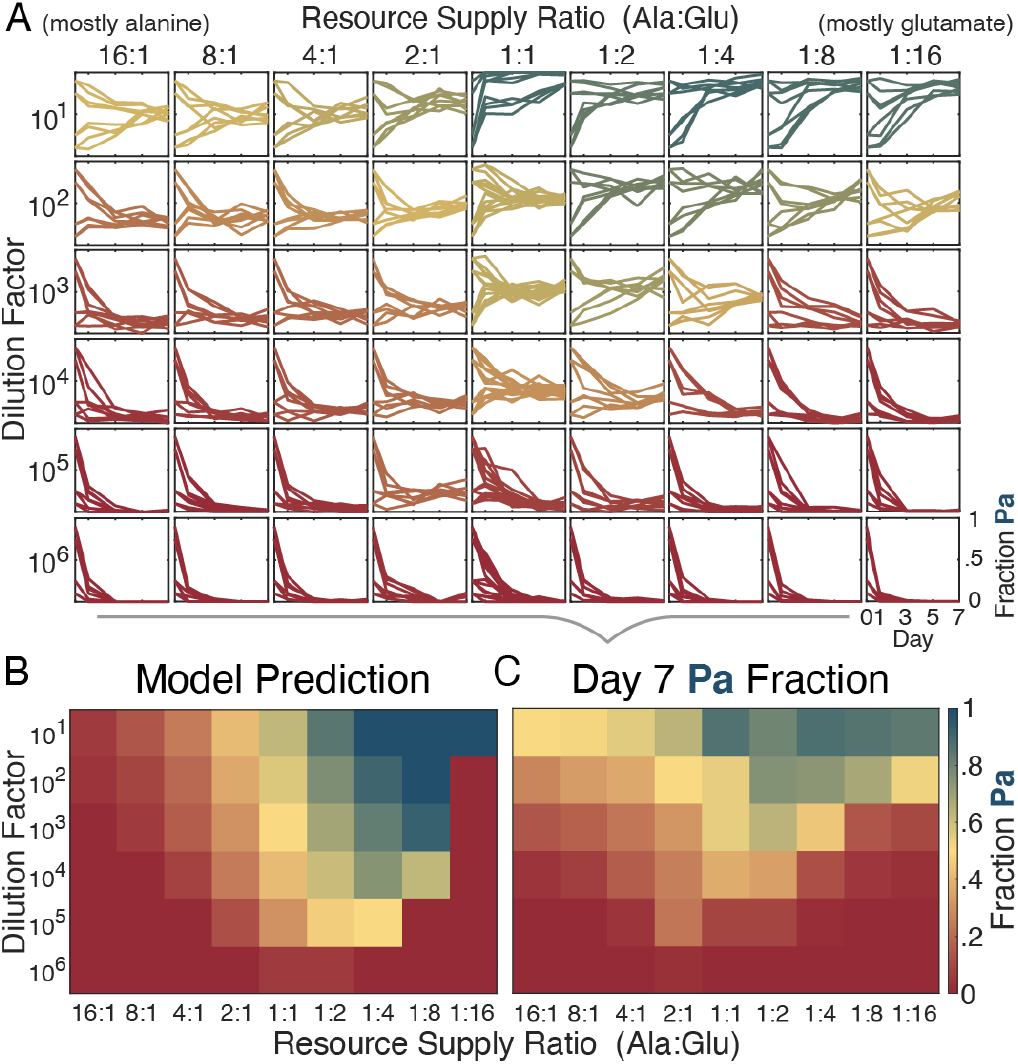
Diauxic-lag model successfully predicts how community composition responds to changes in the dilution factor and resource supply. (*A*) In order to validate the modeling and test its predictive power, we repeated the competition experiments between Aci2 and Pa across a 54-condition grid of resource supply ratios and dilution factors. 8 replicates were run for each experimental condition. (*B*) We used the model to predict steady-state population fractions for each condition. (*C*) We calculated mean final population fractions from the experimental data and compared these fractions to the model predictions.

The model and experiment agreed on several notable features (Fig. 5*C* in comparison to Fig. 5*B*). First, both featured a broad region of coexistence that covered more than half the conditions tested (32/54 in model, 42/54 in experiment). Second, Aci2 competitively excluded Pa at large dilution factors regardless of resource supply ratio due to species spending more time in the initial growth phase, during which Aci2 had an advantage. Aci2 also excluded or nearly excluded Pa at intermediate dilution factors and large supply fractions of either alanine or glutamate. This trend centered the region of coexistence on roughly equal resource supply ratios (Fig. 5*B* and 5*C*).

Third, the model accurately predicted that Pa would achieve its largest population fractions at low dilution factors and large glutamate supply fractions. This exclusion was the result of relatively little growth occurring before alanine ran out and more growth occurring during Aci2’s diauxic lag when Pa had a strong but temporary advantage. The model predicted four conditions at which Pa would even competitively exclude Aci2 (Fig. 5*B*). This competitive exclusion was not observed experimentally, but there were four conditions at which Pa obtained population fractions greater than 0.83 (Fig. 5*C*). That the slow-grower could become the majority of the population at specific conditions was a surprising and nontrivial result of both the model and experiment.

The model did not, however, capture all the experimental observations. In particular, at high alanine supply fractions and low to intermediate dilution factors, the model predicted exclusion or near-exclusion of Pa whereas the experimental observation was coexistence. This discrepancy was also present to a lesser degree at high glutamate supply fractions and notably at low dilution factors in single-resource competitions (Supp. Fig. 5). A likely explanation for unpredicted coexistence would be cross-feeding, which is expected to become more relevant at low dilution factors because species spend more of their time in a densely populated culture with the excretions from many individuals. In the Supplemental Information we explore the addition of cross-feeding steps after growth on alanine and glutamate and show that this addition could explain the single-resource coexistence and provides a better fit to the experimental data at low dilution factors (Supp. Fig. 9).

Overall, across all conditions tested the root-mean-square error between the model prediction and the experimental observation was a species fraction of 0.243 (Supp. Fig. 7). This error could have been reduced by adding additional model components or fitting parameters to competitive outcomes unconstrained by the monoculture characterizations, but we were encouraged that a model based entirely on monoculture measurements could correctly predict patterns in competitive outcomes across a wide range of experimental conditions and took this result as confirmation that the competitive dynamics between Aci2 and Pa are primarily driven by the tradeoff between Aci2’s fast growth and Pa’s short diauxic lags.

### Coexistence is more likely when slow-growers are fast-switchers

Having established the fitness tradeoff between Aci2’s fast growth and Pa’s fast diauxic shift as the mechanism by which they coexisted, we proceeded to explore whether this fitness tradeoff could be a common mechanism for coexistence between species. We added *Pseudomonas putida* (Pp), *Klebsiella aerogenes* (Ka), and an *Arthrobacter* species (Arth) to our set of species and fructose, glucose, citrate, and aspartate to our set of resources. We performed coculture competitions for each species pair in each two-resource environments (Supp. Fig. 42). We then determined initial growth rates and diauxic lag times for each species in each two-resource environment (Supp. Fig. 43) and calculated differences in growth rates and lag times for each species pair in each two-resource environment. We excluded some cases in which we could not fit growth rates and lag times, such as when a species did not grow on both resources. There were 51 competitions in which the slow-grower was the fast-switcher and 26 competitions in which it was not. The slow-grower was significantly more likely to be the fast-switcher (p = 0.003, binomial test), highlighting that such a trade-off is likely not unusual within microbial communities.

When we compared these categorizations to the competitive outcomes, we saw that coexistence occurred 2.6 times more frequently when the slow-grower had been the fast-switcher (p=0.04, Fisher’s exact test), occurring in 39% +/− 7% of cases compared to just 15% +/− 7% of cases (Fig. 6*B*). These results do not prove that the tradeoff between growth rate and diauxic lag time caused coexistence in each of these cases, but they do argue that such tradeoffs may be a significant factor supporting the coexistence of species in multi-resource environments.

**Fig. 6.**
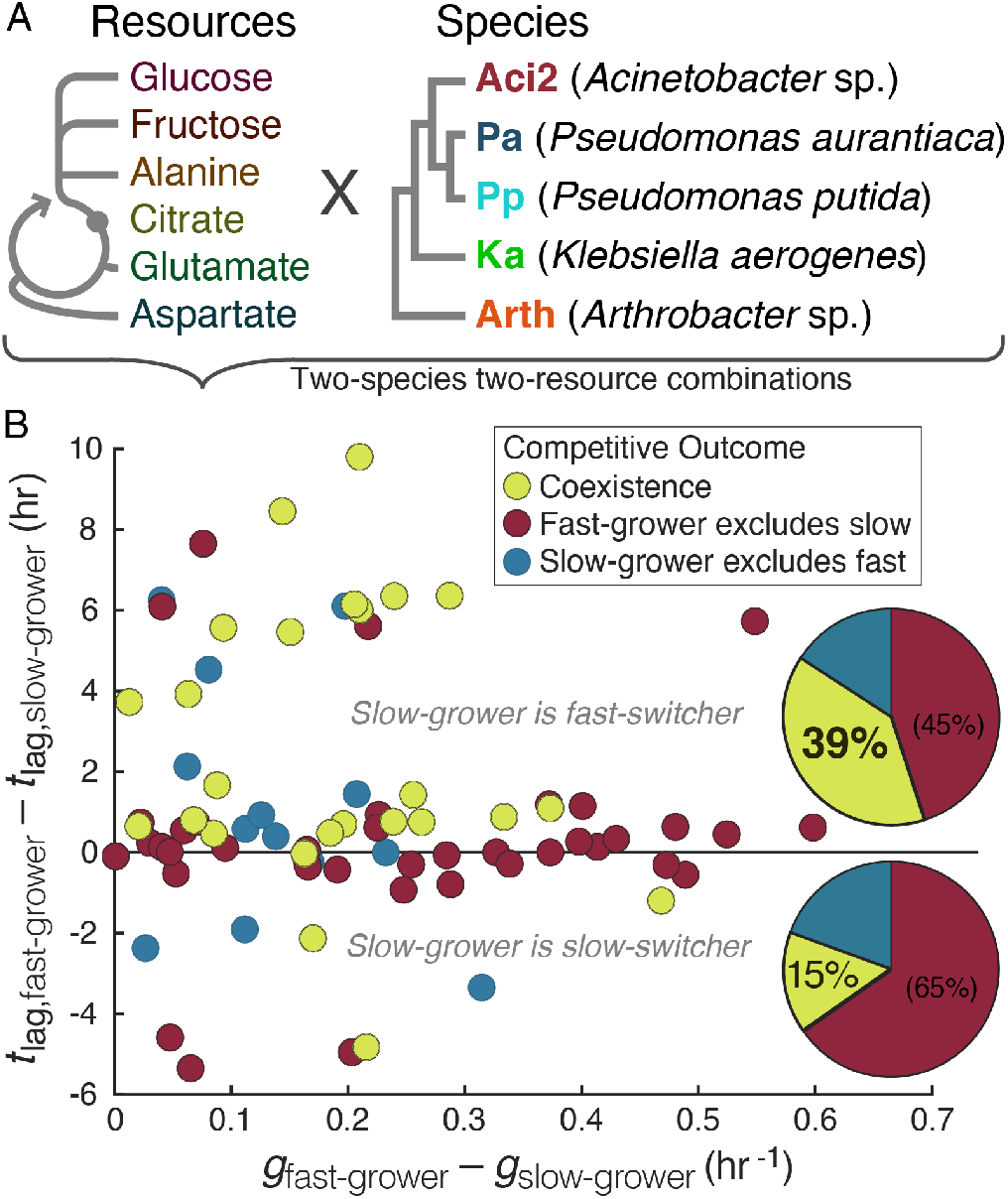
Slow-growers are more likely to coexist with fast-growers when the slow-growers are fast-switchers. (*A*) We selected six resources and five species for additional experiments. Resources are shown in relation to glycolysis and the citric acid cycle. Species are shown with a phylogenic tree with connections in the correct order but not to scale. (*B*) Monoculture experiments were performed to determine species’ growth rates and lag times for each two-resource environment. For each pair of species, coculture competition experiments were performed in each two-resource environment, and qualitative competitive outcomes were determined with a coexistence–exclusion cutoff at a species fraction of approximately 0.01. There were 77 competitions in which both species could be assigned growth rates and lag times. Each point represents one competition. Pie charts summarize the competitive outcomes for cases in which the slow-grower was the fast-switcher (above the Δ*t*_lag_ = 0 line) and in which the slow-grower was also the slow-switcher or neither species had a significant lag time (on and below Δ*t*_lag_ = 0).

This data also contained examples of species being the slow-switcher in some scenarios but the fast-switcher in others. For example, Aci2 had been the fast-grower and slow-switcher when competing against Pa on alanine and glutamate but became the slow-grower and fast-switcher when competing against Pp on alanine and aspartate. Pp and Aci2 coexisted in this competition with Pp performing much better than would have been predicting by averaging the single-resource outcomes, obtaining a population fraction of 0.65 compared to the prediction of 0.07 from averaging (Supp. Fig. 42*D*, 43*A*, and 43*C*). In the other direction, Pa became the fast-grower and slow-switcher when competing against Ka on fructose and aspartate, another competition in which species coexisted (Supp. Fig. 42H, 43B, and 43*D*). These examples of flips in the relative orderings of growth rates and lag times highlight how important the specific environment is to the growth dynamics of species with potentially significant impacts on community assembly.

## Discussion

In this paper we began by investigating a case of unexpected coexistence in a two-resource environment with Pa surviving on the combination of alanine and glutamate despite being competitively excluded on either resource alone and despite being the consistent slow-grower. We discovered that, while Pa was the slow-grower compared to Aci2, it was the fast-switcher. We developed a model based on the tradeoff between Aci2’s fast growth and Pa’s short diauxic lag and showed that this tradeoff was sufficient to explain the coexistence result. The model even predicted how the community composition would respond to environmental changes. We finished by showing that across at set of 77 two-species two-resource competitions the slow-grower being the fast-switcher made coexistence nearly three times as likely to occur. Our results establish the power of diauxic lags to effect coexistence and suggest such a mechanism may be common.

Previous studies have identified metabolic tradeoffs whereby fast-growers optimize their growth strategies for their current environment at the expense of being able to quickly adapt after resource depletions^31,32,39^. When such a tradeoff exists, slow-growers will tend to be fast-switchers while fast-growers will be slow-switchers. In our focal example of Aci2 and Pa, a tradeoff between Aci2’s fast growth and Pa’s short diauxic lags produced stable coexistence. We also showed that slow-growers are more likely to coexist with fast-growers when the slow-growers are fast-switchers. We did not investigate whether such a tradeoff was indeed responsible for coexistence in each of the cases we identified. A similar tradeoff has, however, been seen between different strains of E. coli growing on glucose and acetate^33,34^, establishing that Aci2 and Pa on alanine and glutamate is not the only case in which coexistence is the demonstrable result of a tradeoff between growth and lag. The combination of previous results and those presented here suggests tradeoffs between growth rate and diauxic lag time may be a common mechanism for the coexistence of species, although the relevancy of this mechanism depends on future work establishing the mechanistic connection in more cases.

The relevancy of tradeoffs between growth rate and lag time will also depend on the extension to more natural communities and environments. We considered the case of two species growing on two resources whereas natural ecosystems contain many species and resources. With more than two resources supplied, species may undergo multiple diauxic shifts, which could help larger numbers of species coexist. But, in the many-resource limit, a diauxic lag with every resource depletion would leave species endlessly trapped in lag phases, suggesting that this framework may lose realism in some cases. A recent theoretical study has, however, shown that even short diauxic lags can accelerate an ecosystem’s convergence to a coexistence steady-state^6^. Understanding when discrete growth and lag phases are an accurate picture of microbial growth will be a necessary step in applying tradeoffs between growth rate and lag time to more complex environments.

Cross-feeding is thought to be one of the most important factors in the assembly of natural communities but is not an interaction we sought to study in this paper. There are, however, interesting connections between diauxic lags and cross-feeding. Diauxic lags can occur when a microbe switches from a supplied resource to one of its own byproducts^40–42^. In the case of a primary degrader eating a supplied resource and producing a byproduct and a cross-feeder eating that byproduct, if the primary degrader consumes the byproduct after the supplied resource runs out and grows faster on both resources than the cross-feeder on the byproduct, then in the absence of lags the primary degrader will always be growing faster and always increasing its population fraction relative to the cross-feeder. But if the primary degrader has a diauxic lag, the cross-feeder will have a chance to catch up and maintain its population fraction. In the case of multi-step degradations, a species capable of consuming resources at every step along the way could entirely miss the chance to grow on certain resources if the species can’t switch to a resource before it runs out. In successional dynamics on complex carbon sources there are occasionally early colonizers that decline in relative abundance and later reappear^12^, a phenomenon for which a partial explanation could be diauxic lags. These intersections between diauxic lags and cross-feeding are explored in the Supplemental Information and may turn out to be some of cases in which tradeoffs between growth rates and diauxic lag times are the most ecologically relevant.

In addition to diauxic lags, microbes often display initial lags, which occur when previously stationary species are presented with fresh resources. Theoretical work has shown that tradeoffs between growth rate and initial lag could also be a source of coexistence between species^43,44^. Coexistence derived from initial lags would require only a single growth phase or supplied resource but would occur within comparatively narrow parameter regions and require species to have significantly different yields. This initial-lag model would not have predicted coexistence for our focal example because Aci2 and Pa had nearly equal yields (as well as little-to-no initial lag (Fig. 2*A*)). Even if modeling modifications had been made to allow for initial lags as a source of coexistence in our example, such a model would not have reproduced the experimental response to changes in the resource supply ratio. However, if initial lags were combined with our diauxic lag model, they may have given Pa enough of a boost to allow for coexistence at some of the low-dilution conditions were a discrepancy with the experimental data was observed. Both initial and diauxic lags are potentially important, and likely interacting, sources of coexistence, and future work should explore initial lags more deeply.

Our work adds to a growing body of research exploring the cases in which the fastest-growing species is not necessarily the strongest competitor in an ecosystem. Previous work has shown that decreasing the dilution factor or mortality rate favors slower-growing species. This universal property was originally derived from a Lotka-Volterra model and demonstrated experimentally^45^. Our focal example of Aci2 and Pa on alanine and glutamate maintained this property, and our model predicts this to be true whenever the slow-grower is the fast-switcher, thus providing a resource-explicit mechanism for this trend. In spatially structured ecosystems, it is known that a slow-grower can outcompete a fast-grower by being the fast-disperser and colonizing new locations before the fast-grower can arrive^46,47^. The tradeoff between growth rate and dispersal is a direct parallel to the tradeoff between growth rate and lag time: Aci2 could be loosely thought of as outcompeting Pa on the original alanine island while Pa survives by colonizing the glutamate island faster. We explore in the Supplemental Information whether differing resource preferences could be another tradeoff that allows a slow-grower to survive and in a simple diauxie model discover that without lags differing resource preferences are not enough for a consistent slow-grower to survive. This modeling highlights how diauxic lags produce mechanisms for coexistence beyond those possible in lag-less diauxie. Future research can continue to elucidate the various roles of diauxie and diauxic lags in supporting the coexistence of diverse communities in complex multi-resource environments.

## Materials and Methods

### Species and Media

Aci2 and Arth were previous isolated from soil and identified as an *Acinetobacter* species and an *Arthrobacter* species. Pa, Pp, and Ka were obtained from ATCC. Pa is *Pseudomonas aurantiaca* (ATCC 33663), Pp is *Pseudomonas putida* (ATCC 12633), and Ka is *Klebsiella aerogenes* (ATCC 13048). Species were streaked from glycerol freezer stocks onto agar plates and single colonies were picked for each experiment. Agar plates contained 15 g/L agar, 5 g/L peptone, and 3 g/L yeast extract. This recipe was also used for the colony counting plates.

All experiments were performed in an M9 media. The media contained final concentrations of 11.28g/L Sigma Aldrich M9 Minimal Salts (3g/L KH_2_PO_4_, 0.5g/L NaCl, 6.78g/L Na_2_HPO_4_, 1g/L NH_4_Cl), 1.9mM MgSO_4_, 0.95mM CaCl_2_, 3.8mg/L FeSO_4_-7H_2_O, 3.8mg/L MnCL_2_-4H_2_O, 2.1mg/L CuSO_4_-5H_2_O, 1.2mg/L ZnSO_4_-7H_2_0, 240μg/L NaMoO_4_-2H_2_O, 240μg/L CoCl_2_-6H_2_O, 9.5mg/L Na_2_EDTA, and a total of 0.1%w/v supplied carbon source. Fructose was supplied as D-(–)-fructose, glucose as D-(+)-glucose, citrate as citric acid, alanine as L-alanine, glutamate as L-glutamic acid monosodium salt, and aspartate as L-aspartic acid monopotassium salt. Weight per volume calculations excluded the weight of any metal ions.

### Competition Experiments

Two days before the start of the competitions experiments individual colonies were picked off agar plates and grown in monoculture for 24 hours in LB media. The day before the start of the experiments, the monocultures were diluted 1:1000 and transferred into M9 media containing equal parts alanine and glutamate when experiments related to the case of Aci2 and Pa on alanine and glutamate were being run and into M9 media containing equal parts fructose, glucose, citrate, alanine, glutamate, and aspartate when the survey of additional species and resources was being run. These monocultures were grown for an additional 24 hours before being used to inoculate the experiments. To initialize the coculture experiments, monocultures were mixed at 90:10, 75:25, 25:75, and 10:90 ratios, and the resulting mixtures were diluted by factors matching the experimental conditions and used to inoculate the competition media. 75:25 and 25:75 inoculum ratios were only used for experiments involving Aci2 and Pa on alanine and/or glutamate.

Competitions were incubated at 25°C on 96-well plates with 200uL in each well and orbital shaking at 350 rpm. Every 24 hours, the cocultures were diluted by a factor corresponding to the experimental condition. At the first, third, fifth, and seventh dilutions, the media was diluted by a factor of 106, and 10uL droplets were plated onto agar plates. Between five and ten replicates of the plating were performed. This yielded 100-200 total colonies from most wells. These colonies were counted to determine species’ population fractions.

### Monoculture Growth Experiments

The same two-step starter-culture procedure as used in the coculture experiments was followed in the 48 hours before the start of monoculture experiments. Monoculture experiments involving Aci2 and Pa on alanine and/or glutamate were inoculated with a 10^−4^ dilution. The other experiments were inoculated at 2*10^−3^. (This dilution factor was used instead of 10^−4^ due to the slower growth rates of some species.) Population size was measured as optical density at either 400nm or 600nm every 5 minutes. Plates were kept at 25°C with orbital shaking in between OD measurements. To extract growth rates over time, a linear least square fit was performed on population sizes from a 30-minute rolling window and the slope of this fit was divided by the population size at the center of the window. Replicates were then median filtered.

### Modeling

Growth before any resource depletion was exponential with equal yields. The equations governing this behavior were

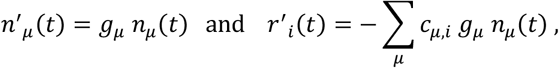

where *n_μ_*(*t*) is the population size of species *μ*, *g_μ_* is the growth rate of species *μ*, *r_i_*(*t*) is the concentration of resource *i*, and *c_μi_* is species *μ*’s consumption fraction of resource *i*. The growth rates used were *g*_Aci2_ = 0.88 hr^−1^ and *g*_Pa_ = 0.67 hr^−1^. The resource consumption fractions in the two resource environments were *c*_Aci2,Ala_ = 1, *c*_Aci2,Glu_ = 0, *c*_Pa,Ala_ = 0.4, and *c*_Pa,Glu_ = 0.6. In single resource simulations consumption fractions were *c_μ_,i* = 1. Note that *r*′_*i*_(*t*) has no dependence on *r*_*i*_(*t*).

After a resource depletion, growth rates recovered as

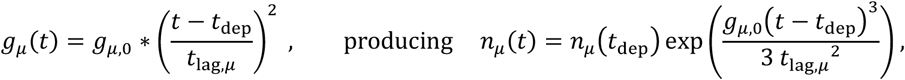

where t_dep_ is the time of the resource depletion (and on set of diauxic lag) and t_lag_ is the lag time. The lag time was constant for Pa at 1 hr and for Aci2 was a linear interpolation of the values presented in Fig. 2*F*, which were 12, 12, 12, 11.7, 10.8, 9.375, 7.875, 6, and 2.475 hours for resource supply ratios of 1:16, 1:8, …, 16:1 alanine:glutamate. Once *g*_μ_(*t*) = *g*_μ,0_ was reached, exponential growth resumed. Consumption of remaining resource *i* after depletion of resource *j* remained governed by *r*′_*i*_ (*t*) = − Σ_μ_ *n*′_μ_ (*t*) during and after the lag phase. All growth stopped after both resources had been depleted.

In the monoculture simulations, population sizes were started at *n*_μ_(0) = 10^−4^ and resource concentrations were started with *r*_Ala_(0) + *r*_Glu_(0) = 1. There was only one species present in the monoculture simulations.

In the competition simulations, resources were supplied at the beginning of each day with *r*_Ala_(0) + *r*_Glu_(0) = 1. At the start of each day, population sizes from the end of previous day were divided by DF. There was a slight carryover such that the total population size after saturation (at *t*_sat_) was 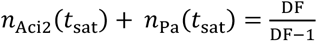 On the first day of the competition simulations, population sizes were started at 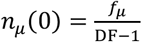. Reported in Fig. 4*B* are the population fractions at the end of each day, *f*_Pa_ = *n*_Pa_(t_sat_)/(*n*_Acl2_(t_sat_) + *n*_Pa_(t_sat_)).

Simulations were performed by directly solving a sum-of-exponentials equation for the time at which resources would be depleted and then calculating species population sizes using that value. No numerical integration of differential equations was involved, so all simulation results are precise.

### Statistical analysis

The uncertainties on the fractions of competitions that produced coexistence came from a beta distribution with

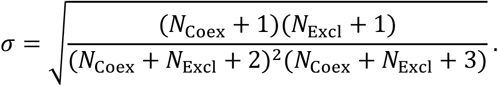

To calculate growth rates and their uncertainties, we selected a 4.5-hour window of monoculture growth data (OD_400_ sampled every 5 minutes) and calculated the slope of a log-linear fit through every possible pair of points. We performed these calculations for three replicates of Aci2 monoculture data and four replicates of Pa monoculture data. The reported growth rates are the median growth rate for each species across all the two-point fits. The reported uncertainties on those growth rates are the standard errors of the distributions of growth rates.

## Acknowledgements

This work was supported by Simons Foundation grant 542385 to J.G. within the Principles of Microbial Ecosystems Collaborative.

