## Supplemental Information for "Diauxic lags explain unexpected coexistence in multi-resource environments"

*for*

#### Contents

**Supplemental Figures** 3

**Supp. Fig. 1.** Aci2 is the fast-grower in both single-resource environments.

**Supp. Fig. 2.** Model predictions for population size at which diauxic shift will begin.

**Supp. Fig. 3.** Additional detail on diauxic lag time fits.

**Supp. Fig. 4.** Growth rates do not vary with resource supply ratio.

**Supp. Fig. 5.** All time-series and final fraction data for Aci2 and Pa competitions at various resource supply ratios, including pure alanine and pure glutamate conditions.

**Supp. Fig. 6.** Comparison of model from paper to using weighted average of single-resource results to predict competitive outcomes in the multi-resource environments.

**Supp. Fig. 7.** Model prediction vs observed outcome across all dilution factors and resource supply ratios tested.

**Supp. Fig. 8.** Comparison of model predictions for Pa as an alanine specialist, as a 2:3 generalist, and as a glutamate specialist.

**Supp. Fig. 9.** Addition of cross-feeding to the model allows for single resource coexistence at low dilution factors

**Key results from exploring simple diauxie models** 9

Without lags, diauxie models do not allow a consistent slow-grower to survive 9
The mapping of population fractions from one day to the next is continuous 9
Having one resource be cross-fed changes calculations and outcomes surprisingly little 9
Multi-stability arise through various mechanisms that require some form of anomalous preference 9

**Simplest model of a tradeoff between growth rate and diauxic lag time** 10

Dynamic equations and simplification to algebraic expressions 10
Fixed points with both species surviving 11
Monoculture steady-states and their invasibility by the other species 13
Qualitative competitive outcomes 14
Species fractions 15
Nonequal resource supplies 17
With opposite resource preferences 18
With different yields 21
No steady-state oscillations 22

*(contents continued on next page)*

**Simple diauxie model of two species on two resources without lags but with different growth rates for each resource** **24**

Species have the same resource preference 24
Species have opposite resource preferences 29
Five qualitatively distinct resource ratio vs dilution factor competitive outcome phase spaces 37
Nonequal yields 38
Separate two-resource co-utilization states 39
Continuity of the day-to-day map 40
Continuity and diauxic lags 43
Continuity and co-utilization 44

**Combination of diauxic lags and species having different growth rates for each resource** **45**

A tri-stability when species have opposite, anomalous resource preferences 45

**Diauxic lags and cross-feeding** **48**

Fast-growing, slow-switching primary degrader vs slow-grower cross-feeder 48
Fast-grower vs fast-switcher during sequential degradation 50

**Co-culture competitive outcomes and growth/lag characterizations for the survey of addition species and resources** **52**

Overview of Supp. Fig. 42 and 43 contents. 52
Supp. Fig. 42*A*–*B*. Summary of how two-resource competition outcomes related to single-resource outcomes 53
Supp. Fig. 42*C*–*L*. Competition data 55
Supp. Fig. 43. Growth rate and lag time characterization 65

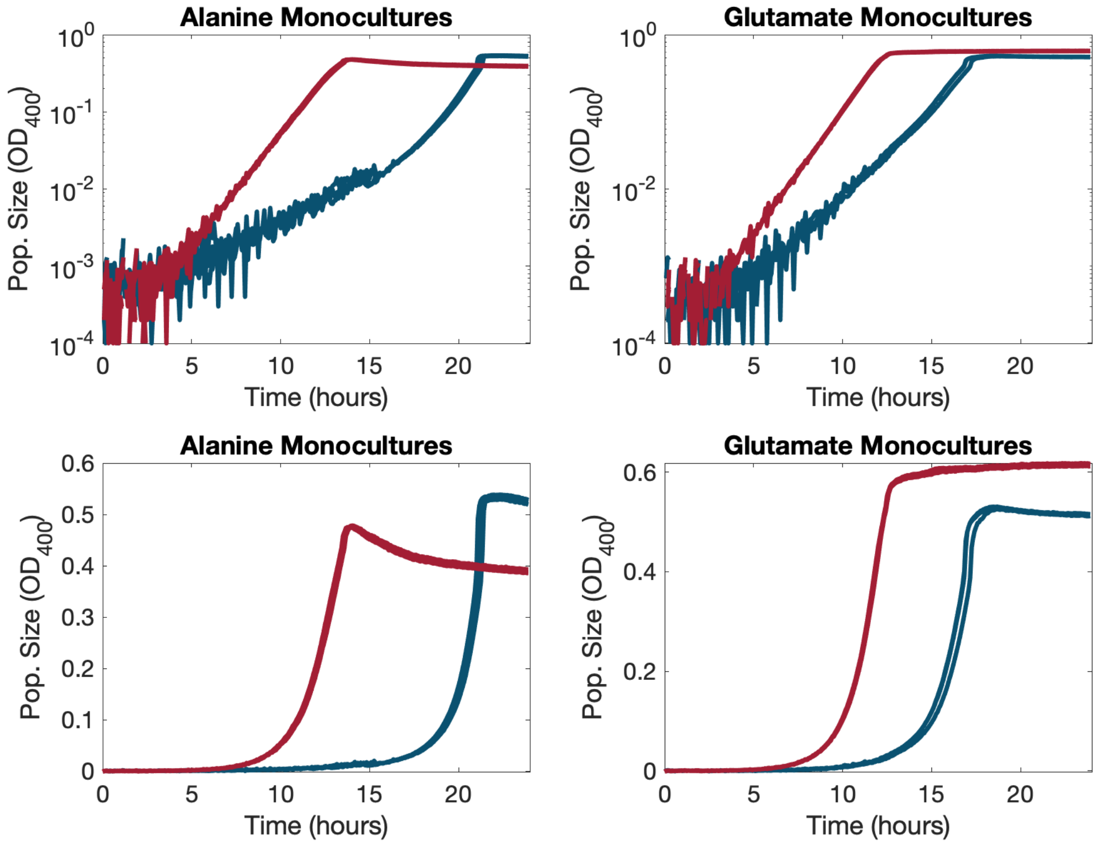


**Supp. Fig. 1.** Aci2 is the fast-grower in both single-resource environments. In all plots Aci2 is in red and Pa is in blue. Top row and bottom row are the same data just on different y-axis scales.

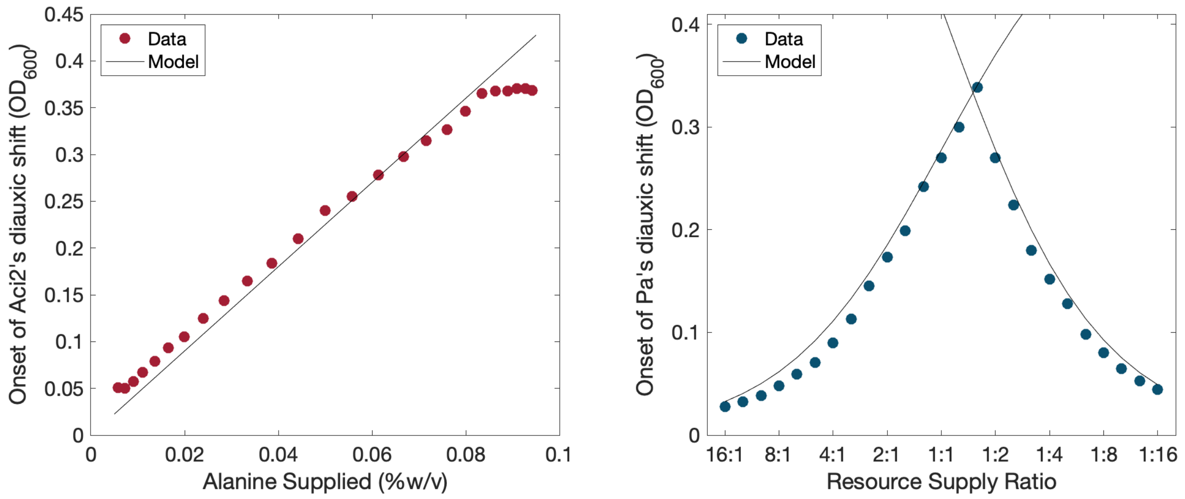


**Supp. Fig. 2.** Model predictions for population size at which diauxic shift will begin. In the Pa plot one line shows the onset associated with alanine running out first and the other shows the onset associated with glutamate running out first. In the Aci2 data, the slight divergence from the linear fit suggests that Aci2 may actually be eating alanine and glutamate in a ~10:1 alanine:glutamate ratio.

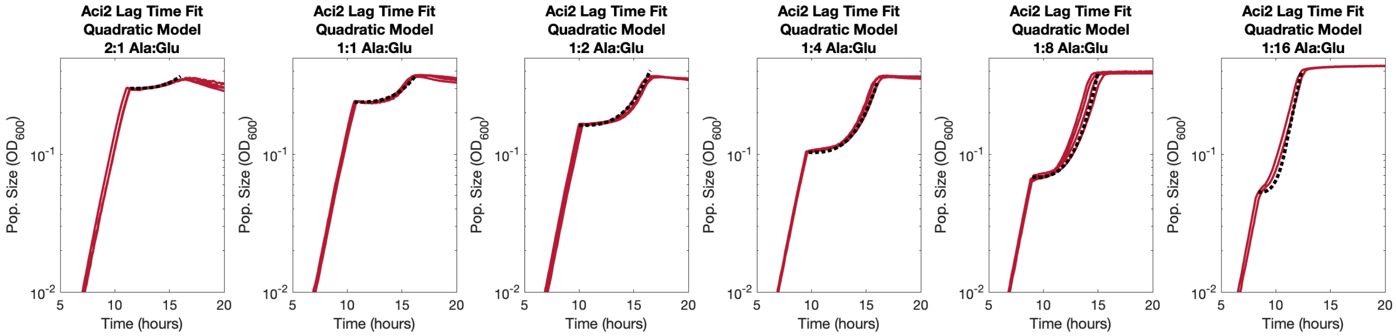


(3$A$) Aci2 lag time fits using quadratic growth rate recovery model used in paper.

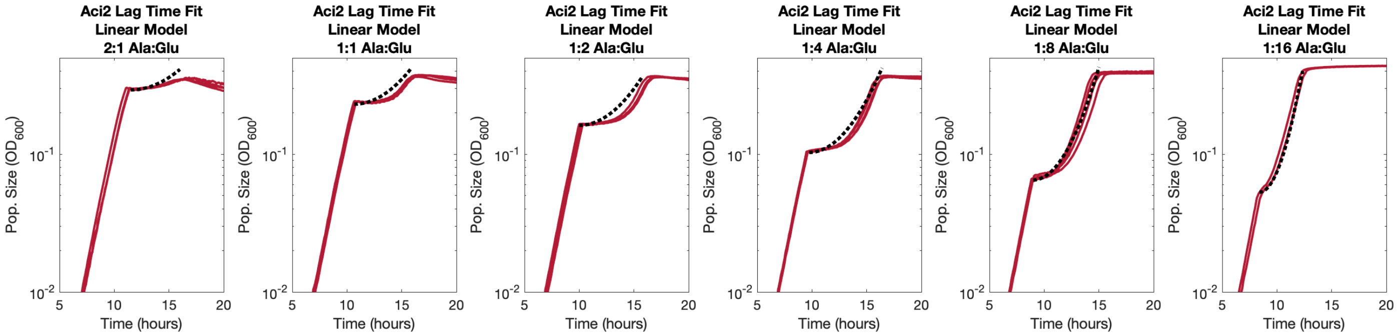


(3$B$) Aci2 lag time fits using alternative linear growth rate recovery model in which growth rate is linearly proportional to the time since the onset of the diauxic shift. Not that for conditions with large alanine supplies, the linear model struggles to capture the long period of essentially no growth immediately after the onset of the diauxic shift while still converging towards the data at later times.

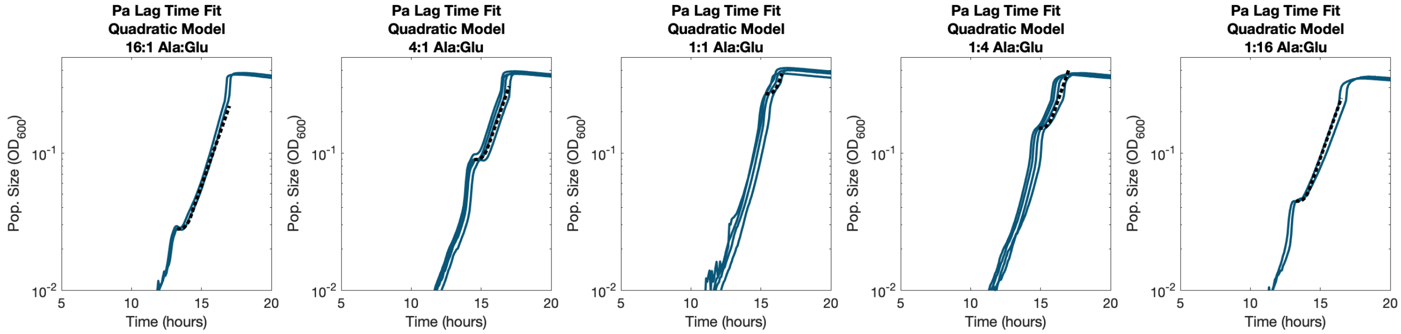


(3$C$) Pa lag time fits using quadratic model and constant value of 1 hour.

**Supp. Fig. 3.** Additional detail on diauxic lag time fits.

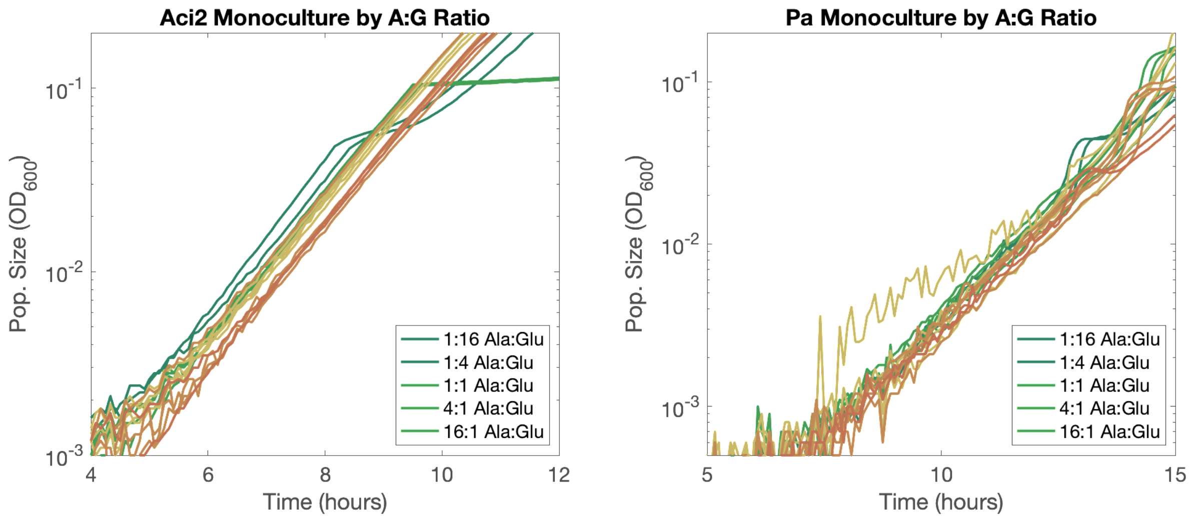


**Supp. Fig. 4.** Growth rates do not vary with resource supply ratio. 2-4 replicates for each condition. Same data as in Main Text Fig. 2.

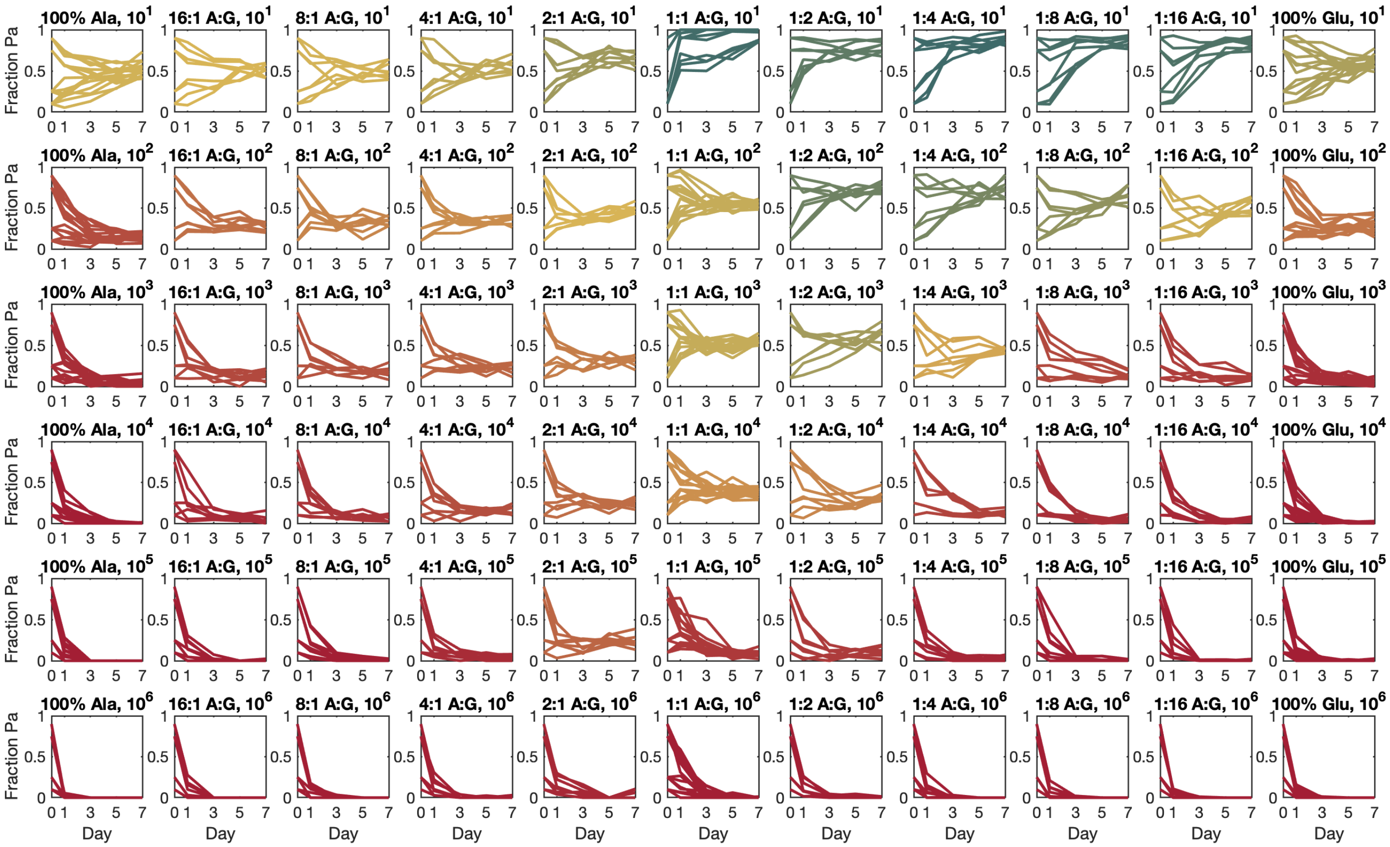


**Supp. Fig. 5.** All time-series and final fraction data for Aci2 and Pa competitions at various resource supply ratios, including pure alanine and pure glutamate conditions. Including the single-resource conditions, 146,566 colonies were counted to produce this data.

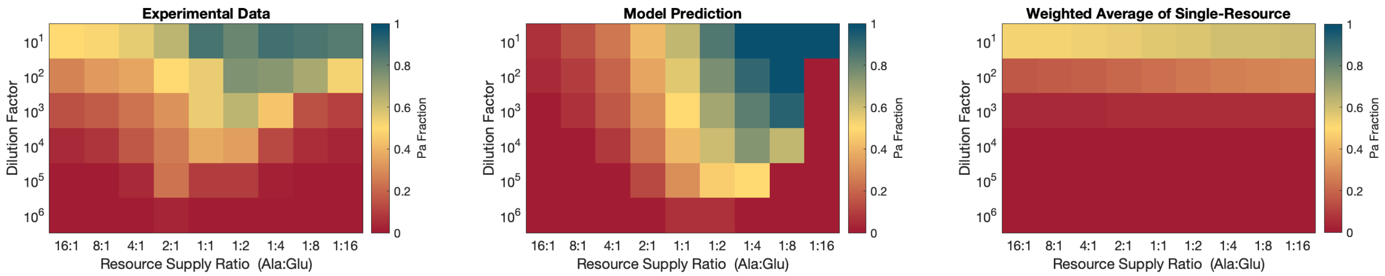


**Supp. Fig. 6.** Comparison of model from paper to using weighted average of single-resource results to predict competitive outcomes in the multi-resource environments. Note the significant amount of structure captured by the model but not by the weighted average alternative.

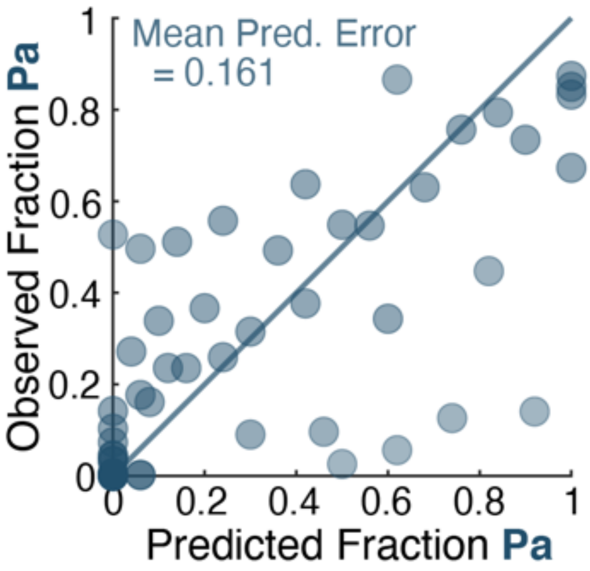


**Supp. Fig. 7.** Model prediction vs observed outcome across all dilution factors and resource supply ratios tested. Model had a mean absolute value prediction error of 0.161, a root-mean-square error of 0.243, and positive correlation to the observed outcomes with $p < 10^{-9}$.

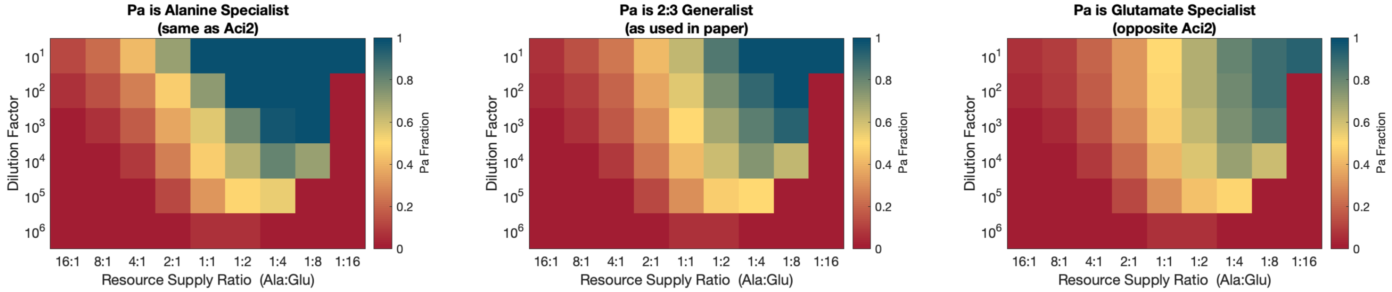


**Supp. Fig. 8.** Comparison of model predictions for Pa as an alanine specialist (same as Aci2), 2:3 generalist (as used in paper), and glutamate specialist. Note that competition outcomes – despite varying considerably with resource supply ratio – are relatively insensitive to Pa's resource consumption ratio. That Pa's resource consumption ratio was relatively inconsequential illustrates that the model's prediction of coexistence was not a result of the two species having different resource consumption ratios.

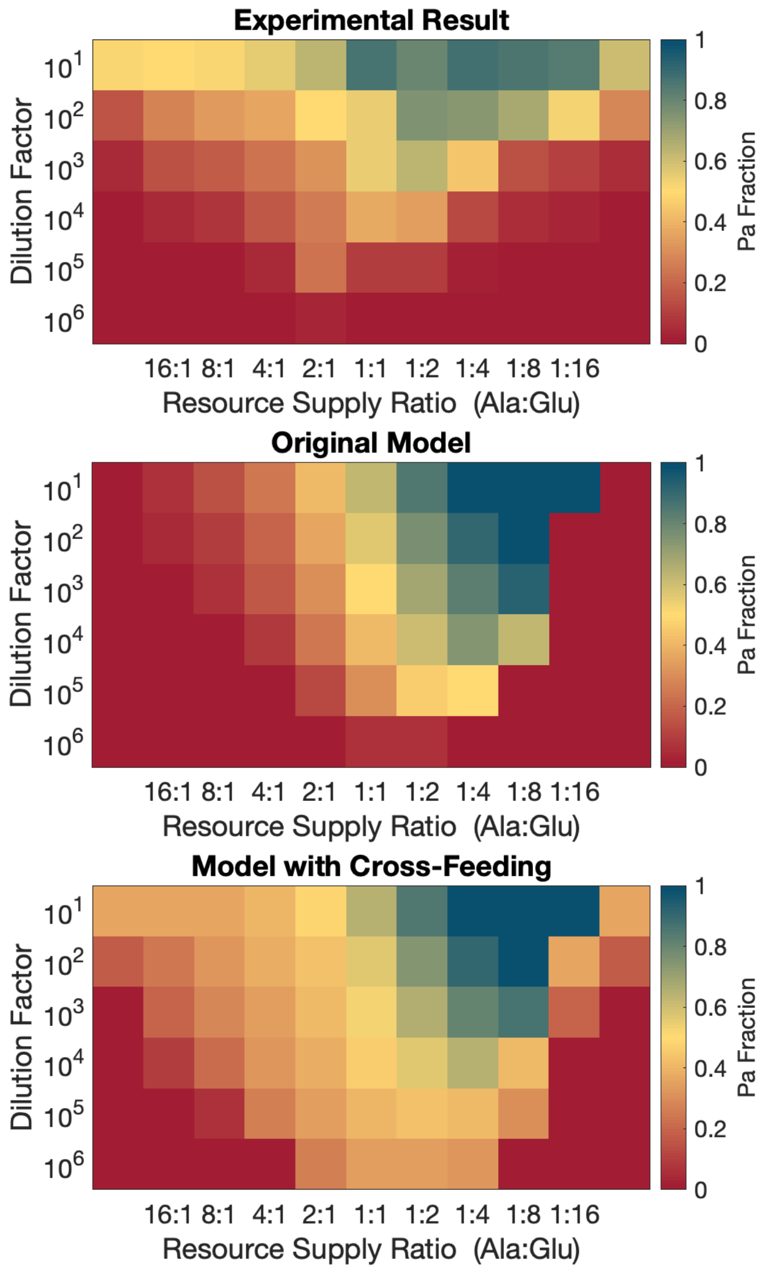


**Supp. Fig. 9.** Addition of cross-feeding to the model allows for single resource coexistence at low dilution factors and generally improves fit slightly at lower dilution factors. Top plot is experimental results with leftmost and rightmost columns displaying pure alanine and pure glutamate results. Middle plot is the model presented in the paper. Bottom plot is the same model with additional cross-feeding after growth on alanine and glutamate. In the cross-feeding model, 20% of the 'biomass potential' of each resource is excreted as a cross-feeding product, with one generalized cross-feeding product excreted from species. After alanine and glutamate are depleted, each species next eats the cross-feeding product that came from the other species and then the cross-feeding product that came from itself. Aci2 has a 3 hour diauxic lag at the start of each cross-feeding stage, and Pa has a 1 hour diauxic lag. Each species grows at its full growth rate on the cross-feeding product that came from the other species and at half its full growth rate on the cross-feeding product that came from itself.

### Key results from exploring simple diauxie models

#### *Without lags, diauxie models do not allow a consistent slow-grower to survive*

In the main text, we used a model focused on diauxic lag times to explain the unexpected coexistence of Aci2 and Pa in an alanine-glutamate environment despite Pa being the slow-grower on both resources and in the two-resource environment and competitively excluded by Aci2 in both single-resource environments. Could a diauxie model without lag times have explained the coexistence of Aci2 and Pa? We explore this question and conclude lag times were essential to our main-text modeling results.

The modeling sections of this Supplemental Information on pages 24-39 examine a simple diauxie model, in which species have a growth rate for each resource and an order in which they consume those resources but no lags. We see that, although coexistence is possible in such a model, in order to survive a species must be the fast-grower on one of the two resources or in the two-resource environment. We add additional factors such as yields and separate two-resource co-utilization states and find that this remains the case. An infinite number of modeling additions could be tested, but we believe that sampled reasonable choices and determine that none of the model variations considered could have produced the coexistence of Aci2 and Pa despite Pa being the slow-grower on both single resources and in the two-resource environment without lags being included in the model.

#### *The mapping of population fractions from one day to the next is continuous*

This is an important property of the model for the validity of various analytical tools. Across pages 40-44, we provide a proof that the simple diauxie model is continuous and that, under some assumptions, this continuity holds even with diauxic lags.

#### *Having one resource be cross-fed rather than supplied changes calculations and outcomes surprisingly little*

In the final modeling section of this Supplemental Information (pages 48-51) we explore the case of one or more resources that are cross-fed. We were surprised to see how little the calculations of whether species would coexist changed. This suggests much of the analysis performed here could be applicable and beneficial to the study of cross-feeding dynamics.

#### *Multi-stabilities arise through various mechanisms in diauxie models of two species on two resources and generally require some form of anomalous preference*

Although tangential to this paper, bistabilities in ecological models are often of interest, so we highlight some cases in which bi-, and even tri-, stabilities emerge:

- Bistabilities do not occur in the lag-and-single-growth-rate model (pages 10-21).
- A bistability can occur if a species prefers the resource it grows slowest on (pages 31-37).
- …if a species prefers the resource it has the lowest relative yield on (page 21).
- …if the fast-grower/slow-switcher is faster on the second resource by a larger factor (page 45).
- A tri-stability can even occur if both species prefer the resource they grow slowest on and the initial fast-grower has a diauxic lag (pages 45-47).

### Simplest model of a tradeoff between growth rate and diauxic lag time

This section provides a more in-depth look at the model presented in the main paper.

Species α and β are growing on resources $R_1$ and $R_2$. Each species has the same growth rate for both resources. These growth rates are $g_\alpha$ and $g_\beta$ with $g_\alpha > g_\beta$.

Both species eat resource $R_1$ first. When $R_1$ runs out, species α has lag $t_{\text{lag},\alpha}$, while β has no lag. Species α's lag is "sharp" in the sense that α does not have any growth until time $t_{\text{lag},\alpha}$ has passed then resumes growing at its full growth rate. Species α having a lag (rather than species β having a lag) is the interesting case to consider because if β is the species with a lag then α will always be growing faster than β and will therefore always being increasing its population fraction and will eventually outcompete β. If both species have a lag, all that matters is the difference in lag times, because the period of time during which neither species is growing can simply be ignored.

After saturation species are diluted by the dilution factor DF and resources are replenished. For now, we assume resources are supplied in equal amounts and all yields are equal. We will refer to the period of time from a dilution and resource replenishment to saturation as a "dilution cycle" or a "day" interchangeably.

#### *Dynamic equations and simplification to algebraic expressions*

Species grow exponentially and resources are depleted as the species grow. The model is therefore governed by underlying differential equations

$$n'_\mu(t) = \begin{cases} 0 & \text{if during } \mu's \text{ lag} \\ g_\mu \, n_\mu(t) & \text{else} \end{cases}$$

$$c'_i(t) = -\sum_\mu h_{\mu,i} \, n'_\mu(t) \quad \text{with} \quad h_{\mu,i} = \begin{cases} 1 & \text{if } \mu \text{ is eating } \mathrm{R}_i \\ 0 & \text{else} \end{cases}$$

where $n_\mu(t)$ is species μ's population size, $g_\mu$ is species μ's growth rate, $c_i(t)$ is the concentration of resource $R_i$, and $h_{\mu,i}$ reflects which resource each species is eating. The model has the discontinuity that

$$n_\mu \to \frac{1}{\mathrm{DF}} n_\mu \quad \text{and} \quad c_i \to s_i \quad \text{when } \forall i \;\; c_i(t) = 0$$

where DF is the dilution factor and $s_i$ is the supply concentration of resource $R_i$.

When species are both growing exponentially starting at time $t = 0$ then $n_\mu(t) = n_\mu(0) \exp(g_\mu t)$, and if both are eating $R_i$ then

$$c_i(t) = s_i - \sum_\mu \left(n_\mu(t) - n_\mu(0)\right) = s_i - \sum_\mu n_\mu(0)\left(\exp(g_\mu t) - 1\right).$$

From here it can be determine that $R_i$ is depleted a time $t_i$ such that

$$\sum_\mu n_\mu(0)\left(\exp(g_\mu t_i) - 1\right) = s_i .$$

And, each species' population size at this time is then calculable from $n_\mu(t_i) = n_\mu(0) \exp(g_\mu t_i)$.

(We note that each term $n_\mu(0)\left(\exp\left(g_\mu\, t_i\right)-1\right)$ is zero at time $t=0$ and then smoothly and strictly monotonically increasing (i.e. with a first time derivative strictly greater than zero) and also smoothly and strictly monotonically increasing with increases to $n_\mu(0)$ and $g_\mu$. Therefore the sum $\sum_\mu n_\mu(0)\left(\exp\left(g_\mu\, t_i\right)-1\right)$ is also monotonically increasing from zero, and the expression for the time at which $R_i$ is depleted,

$$t_i : \sum_\mu n_\mu(0)\left(\exp\left(g_\mu\, t_i\right)-1\right)=s_i\,,$$

has a solution $t_i$ that varies smoothly with all its parameters (by the invertibility of strictly monotonic functions).

These solutions to the differential equations turn out to be the only solutions we need to treat the model from an entirely algebraic perspective for the majority of our analysis going forwards.

#### *Fixed points with both species surviving*

We are primarily interested in studying this model to determine under which conditions species are able to coexist. We define coexistence as needing to be stable – a species going extinction very slowly does not count. Stable coexistence can take the form of (i) a fixed point at which population sizes do not change from one day to the next, (ii) an oscillation in which population sizes change day-to-day but return to the same values every $N$ days, or (iii) a chaotic, pseudo-periodic, or similar state in which population sizes never return to the exact same values but never leave some subrange of values. Fortunately for the simplicity of our analysis, in the case of two species on two resources the only possibility is a fixed point at which population sizes do not change from day-to-day (as is proven later on). Because species have their population sizes divided by the dilution factor DF at the start of each dilution cycle, they must both grow by exactly DF during the growth period of the dilution cycle.

The day-to-day dynamics are a map[1] whereby species population fractions $\boldsymbol{f}$ at the end of one day are a function of the species population sizes at the end of previous day $\boldsymbol{f}_{\text{day }i}=P(\boldsymbol{f}_{\text{day }i-1})$. (It is noted below that species fractions and population sizes at the end of a day are interchangeable under $n_i=\frac{\text{DF}}{\text{DF}-1}f_i$.) We refer to any species fractions $\boldsymbol{f}^*$ with all species surviving ($\forall_i\ f_i>0$) and the fractions not changing day-to-day ($\boldsymbol{f}^*=P(\boldsymbol{f}^*)$) as a fixed point. **For simplicity, when "fixed point" is used in this Supplemental Information it will always imply a fixed point with *all* species surviving.** All stable coexistence states must be fixed points, but fixed points may be stable coexistence states or the point that separates two basins of attraction when a bistability occurs.

At steady-state, it is possible for α to finish its lag in time to eat $R_2$ or for resource $R_2$ to run out before α's lag is over such that α only ever grows on $R_1$. We treat these cases separately as we look for the fixed points of the dynamics. These fixed points, if they correspond to both species surviving, must have each

[1] Specifically, this is a continuous Poincare map of the underlying dynamics with domain $\boldsymbol{f}\in\{\mathbb{R}^{N_{\text{sp}}} : \sum_i f_i=1\ \&\ \forall_i\ f_i\geq 0\}$. A proof for continuity is provided later in the Supplemental Information.

species growing by exactly the dilution factor on each cycle or else one species will be falling behind while the other pulls ahead (in terms of population fractions). There can be a feedback between population sizes and resource depletion times, so a fixed point can have a "fine-tuned" set of depletion times, but cannot require fine-tuned growth rates, lag times, or dilution factors as these are constant for each specific co-culture competition.

**In the case of α finishing its lag in time to grow on $R_2$**, α grows by a factor of $\exp(g_\alpha t_1)$ on $R_1$ and by a factor of $\exp(g_\alpha(t_2 - t_1 - t_{\text{lag},\alpha}))$ on $R_2$, where $t_1$ and $t_2$ are the times at which $R_1$ and $R_2$ run out. Combining these two growth phases, species α grows (on log scale now) by

$$\Delta \log(n_\alpha) = g_\alpha(t_2 - t_{\text{lag},\alpha}) .$$

Meanwhile species β will grow by a factor of $\exp(g_\beta t_1)$ on $R_1$ and a factor of $\exp(g_\beta(t_2 - t_1))$ on $R_2$ for a total (again switching to a log scale) of

$$\Delta \log(n_\beta) = g_\beta t_2 .$$

Each species' total growth depends only on the time at which the second resource runs out. Therefore (unless $t_{\text{lag},\alpha} = (1/g_\beta - 1/g_\alpha)\log(\text{DF})$ exactly) there is no set of resource depletion times which will allow both species to grow by DF on each cycle and there can be no fixed point with both species surviving (assuming α gets a chance to eat $R_2$).

**In the case of $R_2$ running out before α has finished its lag**, the species' total growths are

$$\Delta \log(n_\alpha) = g_\alpha t_1 \quad \text{and} \quad \Delta \log(n_\beta) = g_\beta t_2 .$$

It is now possible to solve the fixed point condition $\Delta \log(n_\alpha) = \Delta \log(n_\beta) = \log(\text{DF})$.

At any fixed point with both species surviving

$$t_1 = \frac{1}{g_\alpha}\log(\text{DF}) \quad \text{and} \quad t_2 = \frac{1}{g_\beta}\log(\text{DF}) .$$

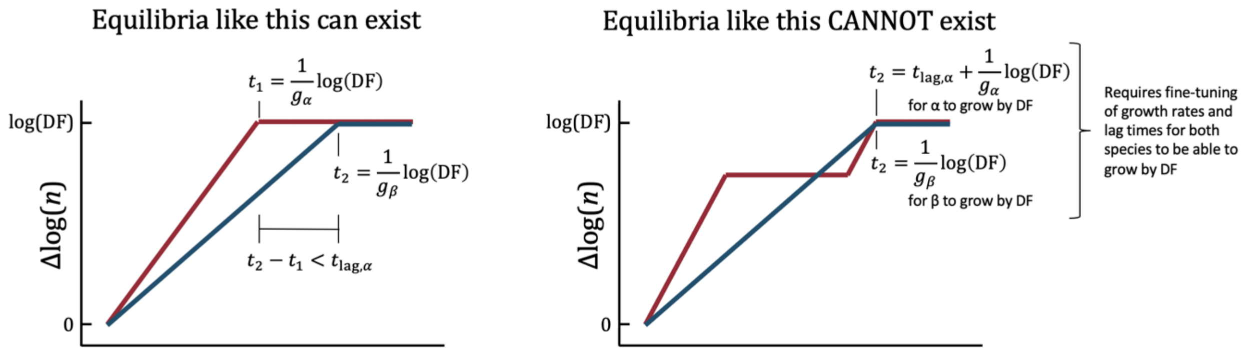


**Supp. Fig. 10.** At a fixed point (with α and β both surviving) α does not finish its lag in time to grow on $R_2$ because doing so would place a second constraint on $t_2$, which is already fully constrained by β's growth-by-DF requirement. If α is able to finish its lag one of the species will be outpacing the other.

It will be shown later on that with an equal number of surviving species and supplied resource (e.g. two of each), a set of resource depletion times corresponds to a unique fixed point.

##### *Monoculture steady-states and their invasibility by the other species*

The previous subsection tells us that there is at most one fixed point with both species coexisting. We can also assume the day-to-day mapping is continuous (as is proven in the section "Continuity of the day-to-day map" later on). This means we can determine qualitative competitive outcomes solely from looking at invasibility of monoculture steady-states by the other species.

Mathematically, invasibility of species μ's monoculture steady-state by species ν is determined by considering the case of $f_\mu = 1 - \epsilon$ and $f_\nu = \epsilon$ with $\epsilon \ll 1$ (i.e. when there's just a tiny fraction of species ν). In this case species ν's population size is considered to be so small that it has no impact on the resource depletion times, and those times are therefore identical to the resource depletion times in species μ's monoculture steady state. If over the course of one dilution cycle $\Delta\log(n_\nu) < \log(\mathrm{DF})$ then species ν's population size converges to zero. If over the course of one dilution cycle $\Delta\log(n_\nu) > \log(\mathrm{DF})$ then species ν population size increases and species μ is unable to drive species ν to extinction and ν survives.

If we know that neither species can drive the other to extinction there must be coexistence. If one species can drive the other to extinction but not the other way around (and if there is not at least two fixed points in between) then the outcome is competitive exclusion. For this reason, the invasibility of monoculture steady-states is relevant to competitive outcomes.

**In α's monoculture steady-state,** α will always get a chance to eat $R_2$ because there is no other species to finish $R_2$ during α's lag. Therefore, the time at which α finishes the second resource is simply the time α needs to spend growing to keep up with the dilution factor plus the time it loses to its lag:

$$t_{2|\alpha} = \frac{1}{g_\alpha}\log(\mathrm{DF}) + t_{\mathrm{lag},\alpha}$$

(Notation $t_{i|\mu}$ indicates the time at which resource *i* runs out in species $\mu$'s monoculture steady state.)

Species β can invade α's monoculture steady state iff

$$g_\beta\left(\frac{1}{g_\alpha}\log(\mathrm{DF}) + t_{\mathrm{lag},\alpha}\right) > \log(DF)\,.$$

**In β's monoculture steady-state,** if β grows by exactly the dilution factor on each cycle then

$$t_{2|\beta} = \frac{1}{g_\beta}\log(DF)\,.$$

From here we might be tempted to naïvely write that species α can invade β iff:

$$g_\alpha\left(\frac{1}{g_\beta}\log(\mathrm{DF}) - t_{\mathrm{lag},\alpha}\right) > \log(\mathrm{DF})$$

This, however, is not entirely true because it assumes $t_{\mathrm{lag},\alpha} < t_{2|\beta} - t_{1|\beta}$ (i.e. that α grows at least a little on $R_2$). Deriving the condition more carefully, α can invade β iff the sum of its log-scale growth on $R_1$

and its log-scale growth on $R_2$ is greater than log(DF). Mathematically enforcing that α's population does not shrink on $R_2$ if it does not finish its lag gives the condition that α can invade β iff:

$$g_\alpha t_{1|\beta} + \max\left(g_\alpha\left(t_{2|\beta} - t_{1|\beta} - t_{lag,\alpha}\right), 0\right) > \log\,(\mathrm{DF})$$

$$g_\alpha \max\left(t_{2|\beta} - t_{lag,\alpha}, t_{1|\beta}\right) > \log\,(\mathrm{DF})$$

To proceed with this calculation, we must determine $t_{1|\beta}$. (Interestingly, this will be the first point in this derivation where the resource supply ratio is relevant, as will be revisited later.)

There's a small carry-over effect from one cycle to the next, so (with total resource supply equal to one and all yields also equal to one) steady-state population sizes saturate at $n_{\mathrm{sat}} = \frac{\mathrm{DF}}{\mathrm{DF}-1}$ and start each day at $n(0) = \frac{1}{\mathrm{DF}-1}$.[2] With the supply of $R_1$ being $s_1 = \frac{1}{2}$, β will have a population size of $n_\beta\left(t_{1|\beta}\right) = \frac{1}{\mathrm{DF}-1} + \frac{1}{2}$ when it finishes $R_1$. So it's total growth is

$$\log\left[n_\beta\left(t_{1|\beta}\right)\right] - \log\left[n_\beta(0)\right] = \log\left[\frac{\frac{1}{\mathrm{DF}-1} + \frac{1}{2}}{\frac{1}{\mathrm{DF}-1}}\right] = \log\left[\frac{1}{2}(\mathrm{DF}+1)\right].$$

Growing at a constant rate $g_\beta$ means

$$t_{1|\beta} = \frac{1}{g_\beta}\log\left[\frac{1}{2}(\mathrm{DF}+1)\right].$$

So, α can invade β's monoculture steady-state iff

$$g_\alpha\left(\frac{1}{g_\beta}\log(\mathrm{DF}) - t_{\mathrm{lag},\alpha}\right) > \log(\mathrm{DF}) \quad \text{or} \quad \frac{g_\alpha}{g_\beta}\log\left[\frac{1}{2}(\mathrm{DF}+1)\right] > \log(DF)\,.$$

Noticed that the first of these inequalities (corresponding to α growing on $R_2$ while invading β) is satisfied if and only if β cannot invade α. For any set of species and environmental parameters such that β survives, β can invade α. This means the first possibility for α being able to invade β (the one in which α gets to eat some of $R_2$) is not satisfiable in any parameter region in which β survives. This will be notable below because it means any boundary between a region of coexistence and a region of β excluding α will not depend on α's lag time because α does not finish its lag (so $t_{lag,\alpha}$ might as well be infinitely long, for example).

#### *Qualitative competitive outcomes*

Because the dynamics are continuous and there exists at most one fixed point, we can deduce qualitative competitive outcomes from invasibility. If both species can invade each other they coexist, and if one species can invade the other but cannot itself be invaded then that species excludes the other.

(If neither species were able to invade the other there would be a bistability with the fixed point separating the basins of attraction, but this does not occur in this model. Likewise, when one species

---

[2] One could take a large DF limit, but the dilution factor isn't necessarily large – researchers often perform experiments with $DF \approx 2$ – and, having tried things both ways, in our opinion it's easiest to just carry few the small number of extra terms than to keep track of appropriately and consistently taking the large DF limit to the same extent in all calculations.

can invade the other but cannot itself be invaded it is possible to have a bistability between an exclusion state and a coexistence state, but this requires two fixed points and this model produces at most one.)

Using invasibility requirements, we can plot qualitative competitive outcomes at $\mathrm{DF} = 10$ and $g_\alpha = 1$ across a range of $t_{\mathrm{lag},\alpha}$ and $g_\beta$. Setting $g_\alpha = 1$ defines an overall timescale. Different values of DF are explored in the next section “Species fractions”.

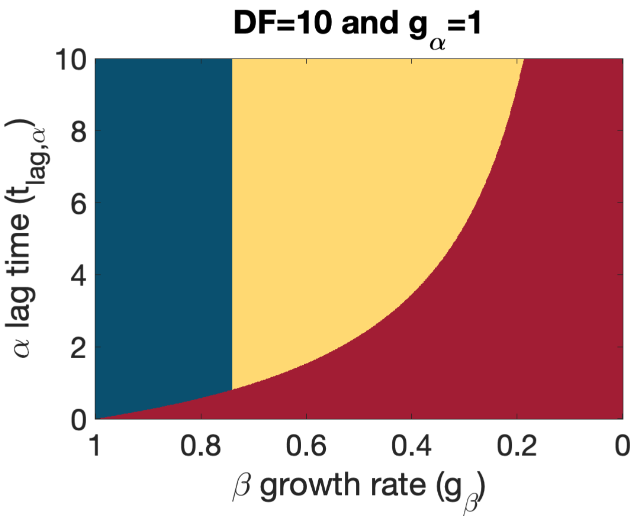


**Supp. Fig. 11.** Predicted *qualitative* outcomes between a fast-grower and a fast-switcher. Red represents the fast-grower α competitively excluding the fast-switcher β, blue represents the fast-switcher excluding the fast-grower, and yellow represents coexistence. Species fractions in the coexistence region are calculated below and plotted in Supp. Fig. 12.

As expected, the slow-grower β excludes α when its growth rate is only slightly behind that of the fast-grower α and is itself excluded when it is much slower. The slow-switcher α only ever loses its ability to survive and/or competitively exclude β if its lag time is increased. However, across much of the phase space α’s lag time appears irrelevant. In particular the boundary between β excluding α and coexistence is exactly vertical with no dependence on the fast-grower/slow-switcher’s lag time. This is because in any state in which β survives, α is not finishing its lag and getting a chance to grow on $R_2$, so α’s lag can be increased arbitrarily without any effect.

#### ***Species fractions***

Having established qualitative outcomes, we now move on to quantitative species fractions. We previously showed that if both species survive then $t_1 = \frac{1}{g_\alpha}\log(\mathrm{DF})$. We can use this and a constraint that $R_1$ must be exactly finished at time $t_1$ to calculate the species fractions when coexistence occurs:

$$\sum_{\mu=\alpha,\beta} \left(n_\mu(t_1) - n_\mu(0)\right) = \frac{1}{2}$$

$$\frac{1-f_\beta}{\mathrm{DF}-1}\left(\exp(g_\alpha t_1) - 1\right) + \frac{f_\beta}{\mathrm{DF}-1}\left(\exp(g_\beta t_1) - 1\right) = \frac{1}{2}$$

$$\frac{1-f_\beta}{\mathrm{DF}-1}(\mathrm{DF}-1) + \frac{f_\beta}{\mathrm{DF}-1}\left(DF^{g_\beta/g_\alpha} - 1\right) = \frac{1}{2}$$

$$f_\beta\left(\frac{DF^{g_\beta/g_\alpha}-1}{\mathrm{DF}-1}-1\right)=\frac{1}{2}-1$$

$$f_\beta=\frac{(\mathrm{DF}-1)}{2\ \mathrm{DF}-2\ \mathrm{DF}^{g_\beta/g_\alpha}}$$

We can plot this fraction in the region of coexistence previously identified:

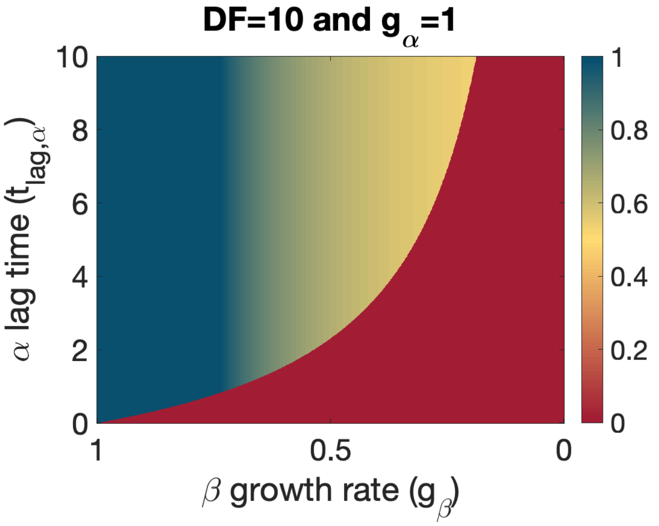


**Supp. Fig. 12.** Same as Supp. Fig. 11 but now plotting the slow-grower/fast-switcher's population fraction. Red is a population entirely composed of the fast-grower/slow-switcher α. Blue is a population entirely composed of the slow-grower/fast-switcher β. Yellow is coexistence at equal species population fractions (not appearing at these conditions), and orange and green are interpolations between those limits.

To further explore the model prediction, we can perform the same calculation for different dilution factors and plot the predicted species fractions:

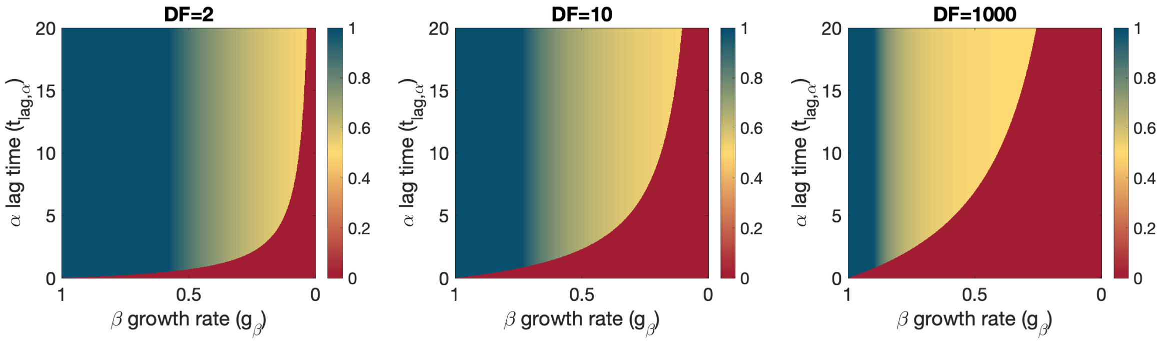


**Supp. Fig. 13.** Same as in Supp. Fig. 12, but with different dilution factors (and the y-axis extended from 10 to 20).

In Supp. Fig. 12 and 13 it is again apparent that once in the region of β surviving (either β excluding α or coexistence) increasing α's lag time has no impact on the population fractions. This is again because once in that region α doesn't finish its lag anyways so it can be made arbitrarily large without impacting anything.

It is also apparent and quite striking that, while the transition from β excluding α to coexistence happens smoothly, the transition from coexistence to α excluding β (and from β excluding α to α excluding β) happens suddenly. Mathematically, this is because there is no coexistence state in which α gets to eat any of $R_2$, which caps α's population at the supply of $R_1$. Intuitively, such a sudden change in population composition is able to occur because when the parameters have shifted just enough for α to get a chance to eat some of $R_2$ there is a sudden positive-feedback whereby (*i.*) α has a new period of growth and increases its population fraction, which (*ii.*) decreases β's population fraction and the amount of $R_2$ that β eats during α's lag, which then (*iii.*) lengthens α's new period of growth and further increases its population fraction.

#### *Nonequal resource supplies*

The resource supply ratios/fractions have no impact on β's ability to invade α, but they do affect α's ability to invade β via changes to the time at which $R_1$ runs out in β's steady state, which affects whether α finishes its lag while trying to invade β.

If the supply fractions are $s_1$ and $s_2$, the calculation for $t_{1|\beta}$ changes to:

$$t_{1|\beta} = \frac{1}{g_\beta}\log\left[\frac{\frac{1}{\mathrm{DF}-1}+s_1}{\frac{1}{\mathrm{DF}-1}}\right] = \frac{1}{g_\beta}\log[s_1(\mathrm{DF}-1)+1]$$

After carrying everything through, α can invade β's steady state iff

$$g_\alpha\left(\frac{1}{g_\beta}\log(\mathrm{DF}) - t_{\mathrm{lag},\alpha}\right) > \log(\mathrm{DF}) \quad \text{or} \quad \frac{g_\alpha}{g_\beta}\log[s_1(\mathrm{DF}-1)+1] > \log(DF)\,.$$

Again, the first condition represents α getting to eat some of $R_2$ while invading β, and the second condition represents α not getting to eat $R_2$ but growing enough of $R_1$ to outpace the dilution.

Meanwhile the species fractions when coexistence occurs are:

$$f_\beta = \frac{(1-s_1)(\mathrm{DF}-1)}{\mathrm{DF}-\mathrm{DF}^{g_\beta/g_\alpha}}\,.$$

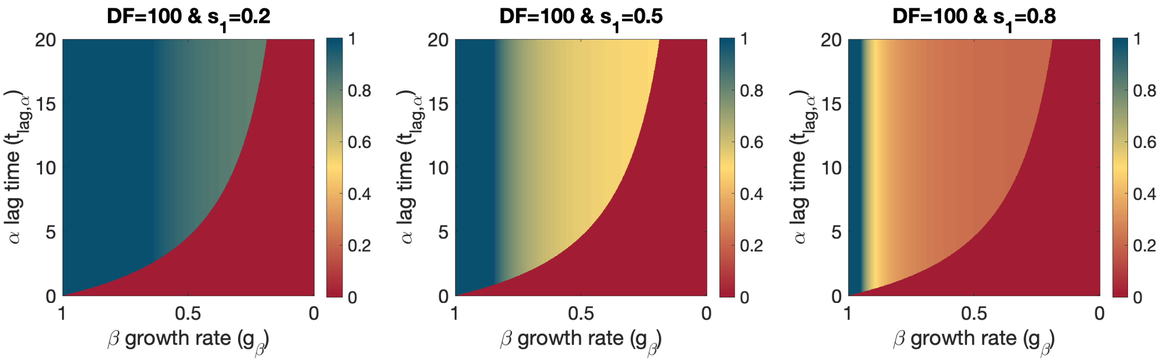


**Supp. Fig. 14.** Varying the resource supply fractions. We again see that α's population fraction in a coexistence state is capped by the supply of $R_1$ (because its lag doesn't end in time for it to eat any of $R_2$).

We continue to see that α's population fraction in any coexistence state is capped by $s_1$.

With changes to resource supply worked out we can also plot dilution factor vs resource supply ratio phase spaces equivalent to what was produced for Aci2 vs Pa in the main text:

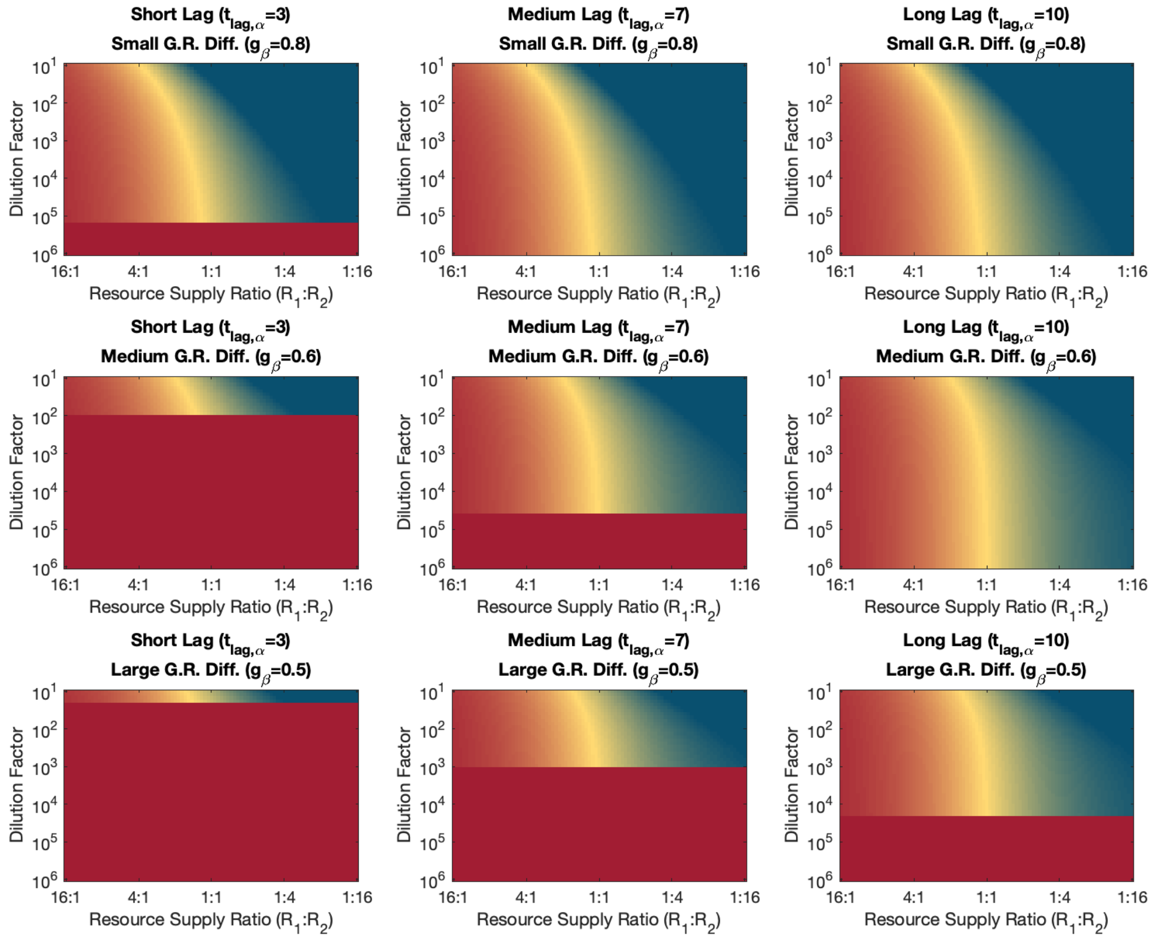


**Supp. Fig. 15.** Phase spaces. Same colormap as in previous figures. In all plots $g_\alpha$=1.

The sudden shift from the upper region of coexistence and β excluding α to the lower region of α excluding β occurs at $\log(\mathrm{DF}) = \frac{g_\alpha g_\beta t_{\mathrm{lag},\alpha}}{g_\alpha - g_\beta}$ (rearrangement of the condition for β invading α) and again appears because of the positive-feedback that occurs when parameters shift enough for α to get a chance to grow on $R_2$.

The sharpness of these phase spaces compared to the predicted Aci2-Pa phase space in the main text is due to the sharpness of Aci2's lag at high supply fractions of $R_1$ (left side; roughly equivalent to large alanine supply) and due to the lag time being constant at high supply fractions of $R_2$ (right side; roughly equivalent to large alanine supply).

#### *With opposite resource preferences*

**If the slow-grower/fast-switcher (β) now eats $R_2$ then $R_1$,** the most impactful change is that, because β depletes $R_2$ first in its monoculture steady state, α will never experience its lag when invading β and can therefore always invade β by being the fast-grower.

It remains the case that species β can invade α's monoculture steady state (and that coexistence will be observed) iff

$$g_\beta\left(t_{\text{lag},\alpha}+\frac{1}{g_\alpha}\log\left(\text{DF}\right)\right)>\log(\text{DF}).$$

So in this simple diauxic lag model, β's ability to survive has no dependence on its resource preference (nor on the resource supply ratio as we have already seen).

At a fixed point with both species surviving, it remains the case that α cannot finish its lag and have a chance to grow on $R_2$ because then each species' growth-by-DF condition would fully constrain $t_2$ to a different value. For a similar reason, β cannot finish $R_2$ before α finishes $R_1$ because then each species' growth-by-DF condition would fully constrain $t_1$ to a different value. Therefore, at any fixed point with both species surviving α is the only species to eat $R_1$ and β is the only species to eat $R_2$. Species fractions are therefore calculated as $f_\alpha = s_1$ and $f_\beta = s_2$.

Plotting these results, the $g_\beta$ vs $t_{\text{lag},\alpha}$ phase space becomes remarkably unfeatured:

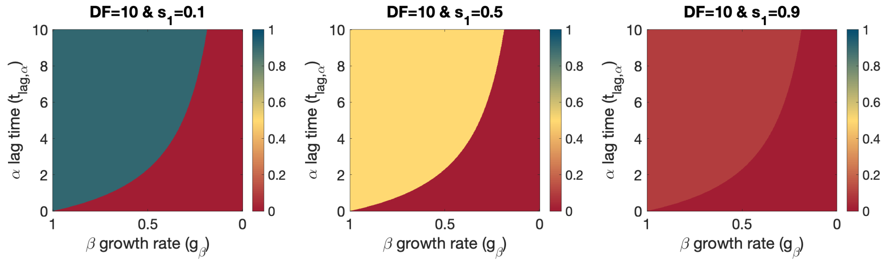


**Supp. Fig. 16.** Population fraction $f_\beta$ when species have opposite resource preferences. ($g_\alpha$=1 in all plots).

Plotting the dilution factor vs resource supply ratio phase space also highlights the drastic de-featuring that occurs when resource preferences are made opposite:

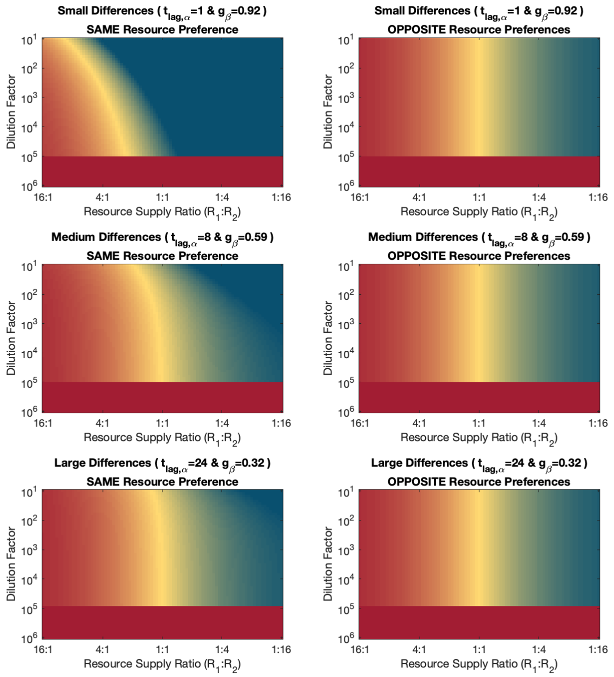


**Supp. Fig. 17.** Comparison of phase space when β has the same resource preference as α (left column) and when β has the opposite resource preference as α. The left column also highlights trends that were less obvious in Supp. Fig. 15. Colormap is same as in all previous figures.

We also notice in Supp. Fig. 17 that β's population is consistently lower when it has the opposite resource preference. This can be mathematically shown to always be true. We previously showed that when species have the same preferences:

$$f_\beta^{\text{same}} = s_2 \frac{(\text{DF}-1)}{\text{DF}-\text{DF}^{g_\beta/g_\alpha}}$$

The term $\frac{(\text{DF}-1)}{\text{DF}-\text{DF}^{g_\beta/g_\alpha}}$ is strictly greater than 1, so $f_\beta^{\text{same}} > f_\beta^{\text{opposite}}$ (because $f_\beta^{\text{opposite}} = s_2$ ). Therefore, having the opposite (compared to same) resource preference will only ever decrease β's population fraction.

**If species have opposite resource preferences and both have a lag time**, their lag times do not necessarily start at the same time so we cannot negate the period when both are lagging and only consider one species to have a lag time. We also don't need $t_{lag,\alpha} > t_{lag,\beta}$ for the problem to be interesting.

It remains the case that α can always invade β's steady-state and that β can invade α's steady state iff

$$g_\beta \left( t_{\text{lag},\alpha} + \frac{1}{g_\alpha} \log(\text{DF}) \right) > \log(DF).$$

The calculation for population fractions also doesn't change:

$$f_\alpha = s_1 \quad \text{and} \quad f_\beta = s_2$$

So, if species have opposite resource preferences, giving the slow-grower a diauxic lag – even one much longer than that of the fast-grower – has absolutely no impact on anything. The simple model of diauxic lags can allow a slow-grower that's also a slow-switcher to survive. This is actually unsurprising if we remember that it was already the case that in all coexistence states $R_1$ ran out first, which means β never experiences its lag anyways.

**Optimal resource preference for a slow-grower.** If we compare the results of the previous two subsections, we see that if β has no lag then its population fraction is only ever decreased by having the opposite resource previous as α. But, if β has the opposite resource preference it can be both the slow-grower and the slow-switcher and still survive.

If β has the same resource preference as α, β can invade α's steady state (and therefore survive) iff

$$g_\beta \left( \left( t_{\text{lag},\alpha} - t_{\text{lag},\beta} \right) + \frac{1}{g_\alpha} \log(\text{DF}) \right) > \log(DF).$$

But if β has the opposite resource preference it can invade α's steady state (and therefore survive) iff

$$g_\beta \left( t_{\text{lag},\alpha} + \frac{1}{g_\alpha} \log(\text{DF}) \right) > \log(DF).$$

So if the slow-grower β has any amount of lag and only cares about whether or not it survives then its best strategy will be to have the opposite resource preference as α.

If β cares about its population fraction, the strategy will depend on the environmental and species parameters.

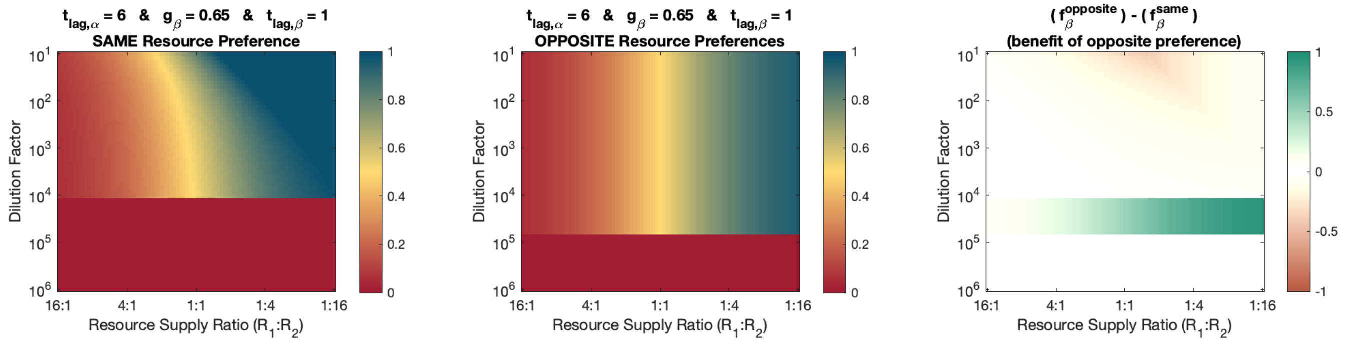


**Supp. Fig. 18.** If the slow-grower (β) has a small lag, it is more likely to survive if it has the opposite resource preference as the fast-grower. But, at certain environmental conditions (specifically small dilution factors) it will obtain a larger population fraction and even exclude α if it has the same preference. Colormap for left and center plots shows species β's population fraction. Colormap for right plot relative benefit of having the opposite vs same resource preference. Blue-green indicates an increased population fraction $f_\beta$ from having the opposite resource preference α. Red-brown indicates an increased population fraction $f_\beta$ from having the same resource preference as α.

We will see a similar case of an initial slow-grower being more likely to survive if it has the opposite preference but having a chance to exclude the fast-grower only if it has the same preference in the case of simple diauxie with no lags but different growth rates for each resource later on.

#### *With different yields*

For simplicity and consistency with our previous treatment of the species' growth being the same on each resource, we consider each species' yield to be the same for both resources (i.e. $y_{\alpha 1} = y_{\alpha 2}$ and $y_{\beta 1} = y_{\beta 2}$). When constrained in this manner, varying yields has no effect on qualitative competitive outcomes, as each species' monoculture carrying capacity will scale with changes to its yield such that it still finishes both resources at the times necessary for it to grow by DF on each cycle. These changes to monoculture carrying capacities carry through to changes to species fractions at coexistence steady-states that produce no qualitatively new features. If both species yields increase or decrease by the same factor, nothing changes, so we only need to consider a single yield $y_\beta$ defined such that if α and β both eat the same quantity of the supplied resource, β's population size will be $y_\beta$ times α's population size. The new species fractions $f_\beta(y_\beta)$ can be calculated as a simple function of the species fractions $f_\beta(1)$ that would have occurred if $y_\beta = 1$.

$$f_\beta(y_\beta) = \frac{y_\beta\, f_\beta(1)}{1 + (y_\beta - 1)\, f_\beta(1)}$$

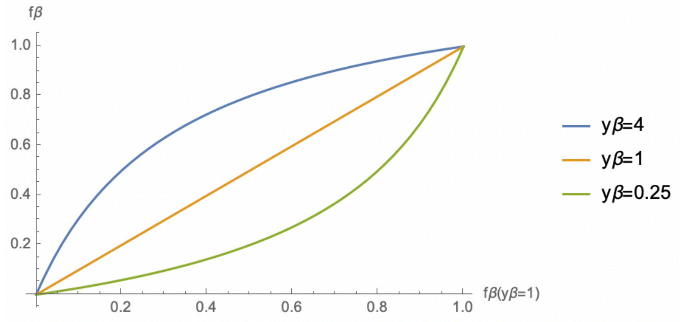


**Supp. Fig. 19.** Species fractions as a function of yield $y_\beta$ and the fractions from calculation assuming $y_\beta = 1$.

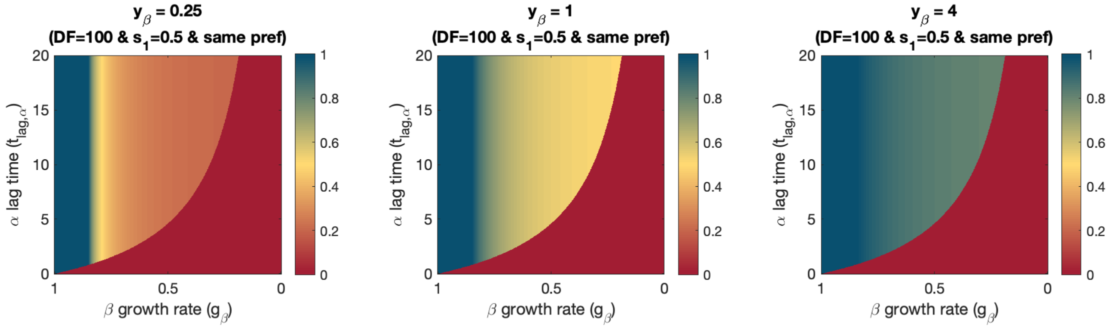


**Supp. Fig. 20.** Varying relative yields affects quantitative species fractions but not qualitative outcomes.

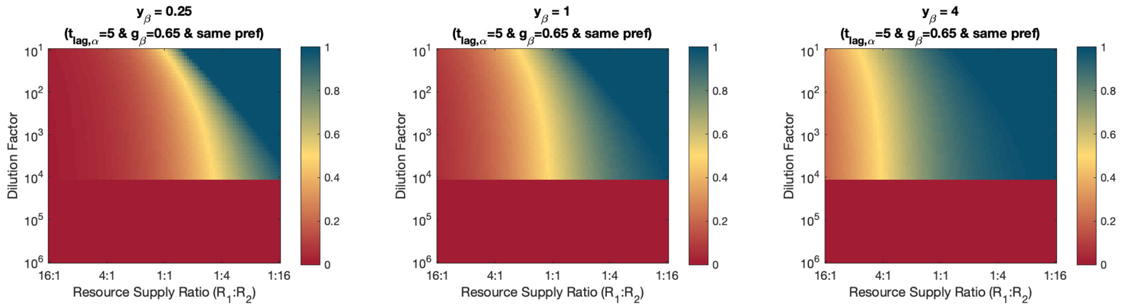


**Supp. Fig. 21.** Same as Supp. Fig. 20, but now plotting the dilution factor vs resource supply ratio phase space. Although varying relative yields does not technically affect qualitative outcomes, doing so can grow or shrink the regions in which a species population fraction is so small it might as well be considered excluded even if it's technically still coexistence.

#### *No steady-state oscillations*

**When species have the same resource preference.** In order to get an oscillation, there needs to be species fractions at which $f_\beta$ increases on the next cycle and species fractions at which $f_\beta$ decreases on the next cycle. Continuity of the map from one day to the next means that in between these two regions there must be a fixed point (i.e. a coexistence steady state). The oscillation will also require that on either side of the fixed point there are species fractions from which the fixed point will be overshoot on the next cycle. If we can show that such points cannot exist on at least one side of the fixed point, then no oscillations can exist. This is true regardless of resource preferences.

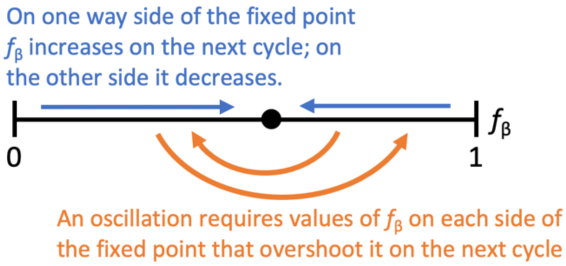


**Supp. Fig. 22.** In order to have an oscillation there must be species fractions on either side of the fixed point that overshoot the fixed point on the next cycle. If it can be shown that such points cannot exist on at least one side of the fixed point then no oscillations can occur.

We previously showed that at the fixed point, α does not finish its lag in time to grow on $R_2$. We now consider the fixed point $f_\beta^*$ and some point $f_\beta^* + \delta f_\beta$ at which β has a larger population fraction than at the fixed point $f_\beta^*$. Our previous arguments for stability established that β's population fraction will decrease on the next dilution cycle to move the system in the direction towards equilibrium, but we have yet addressed whether it's possible β's population can overshoot the equilibrium when starting from some $f_\beta^* + \delta f_\beta$. β's population cannot overshoot for the following reason:

(i) At $f_\beta^* + \delta f_\beta$, β's population fraction is larger and α's population fraction is smaller than at $f_\beta^*$, which means $R_1$ is being eaten slower by the combined population. This increases the time $t_1$ at which $R_1$ runs out. Because β both started with a larger population and grew for longer, β's population size at $t_1$ will be larger when starting from $f_\beta^* + \delta f_\beta$ than when starting from $f_\beta^*$.

(ii) Species β having a larger population size at $t_1$ means that β finishes $R_2$ more quickly. Considering that α could not finishes its lag in time to eat any of $R_2$ when population fractions started at $f_\beta^*$, further decreasing the time period $(t_2 - t_1)$ will ensure α still does not eat any of $R_2$ at $f_\beta^* + \delta f_\beta$. Therefore, β still gets all of $R_2$ to itself.

(iii) Putting everything together, if species fraction start at $f_\beta^* + \delta f_\beta$ then β will end the day with a larger population fraction than if its species fraction had started at the equilibrium $f_\beta^*$. More to the point, starting a day with β having a larger population fraction than at equilibrium never results in β ending that day with a population fraction smaller than the equilibrium. This means there cannot be any steady-state oscillation.

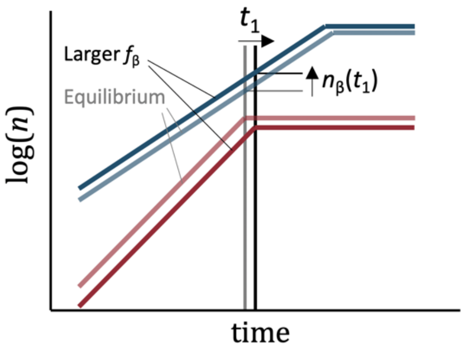


**Supp. Fig. 23.** If $f_\beta$ is perturbed above its equilibrium value, $t_1$ increases, as does $n_\beta(t_1)$, which means β ends the day with its population fraction still above the equilibrium value. There cannot, therefore, be an oscillation around the fixed point because β's population fraction is never able to move from above to below its equilibrium value.

(If the above argument sounds like it should also mean the fixed point is unstable, note that the argument was that β will end the day with a larger population size than it would have at equilibrium, not that it will end the day with a larger population size than it started the day with. Further note that increasing $t_1$ increases α's log-growth, which means $\Delta \log(n_\alpha) > \log(\text{DF})$ over the course of the cycle and α is recovering its population size, making the fixed point stable.)

**When species have the opposite resource preference.** Almost the exact same argument can be made against the possibility of steady-state oscillations, noting that it is still the case that at the fixed point, α is not finishing its lag in time to eat any of $R_2$.

### Simple diauxie model of two species on two resources without lags but with different growth rates for each resource

The primary motivation for this section is to explore whether a diauxie model without lags could have produced the unexpected coexistence of Aci2 and Pa. Specifically, we are interested in whether differing resource preferences or some other characteristic could allow for a species that is the slow-grower in both single-resource environments and in the two-resource environment to coexistence with the fast-grower. We conclude that out of the numerous model variations we explore, lags are essential to have such a result emerge from a diauxie model with two species and two resources.

Same as the previous section, we approach our understanding of the model by searching for fixed points, as their existence is essential for coexistence to occur. We'll be able to perform these calculations using a similar methodology: (i) The resource depletion times can be calculated from a linear condition that each species grows by exactly the dilution factor on each cycle. (ii) The species fractions can be calculated using those times and a condition that the resources are exactly depleted. In this case whether these fractions are all between 0 and 1 is sufficient to determine whether the fixed point exists. (iii) Invasibility of monoculture steady-states can be determined as a coherence check.

This section has some overlap to the Supplemental Information of "Complementary resource preferences spontaneously emerge in diauxic microbial communities" (Wang et al. bioRxiv 2021). These two works were undertaken simultaneously and largely independently, although some authors of Wang et al. had opportunities to preview earlier versions of the work presented here. In comparison to Wang et al. this Supplemental Information presents additional discussion on the effects of tuning experimentally available environmental parameters (dilution factor and resource supply), frames the conditions for coexistence and bistabilities from the perspective of slow-growers vs fast-growers, and provides a proof of the continuity of the day-to-day map (which is necessary for various analytical tools to maintain validity). In comparison to this work, the Supplemental Information for Wang et al. details a visual tool for understanding solutions to a linear constraint that at a fixed point species' growth must match the dilution factor, builds off that tool to take a more abstracted approach, and, in both main text and supplement, provides a discussion of the steady state that is reached when a very large number of species make sequential attempts at invading an ecosystem. We encourage anyone interested in diauxie models to read Wang et al. as well as the material presented here.

#### *Species have the same resource preference*

We'll continue to refer to our species as α and β with both species consuming $R_1$ then $R_2$ and with α as the initial fast-grower. Because growth rates can now vary with which resource is being consumed they now need two subscripts so that $g_{\mu i}$ is the growth rate of species μ on resource $R_i$.

That α is the initial fast-grower means $g_{\alpha 1} > g_{\beta 1}$. Although it isn't necessarily the case that $g_{\alpha 2} > g_{\beta 2}$, which would mean that α is the fast-grower on the second resource as well. Indeed **if both** $\boldsymbol{g_{\alpha 1} > g_{\beta 1}}$

**and $g_{\alpha 2} > g_{\beta 2}$ then α is always growing faster than β and α will exclude β** (because it is always increasing its population fraction relative to β).

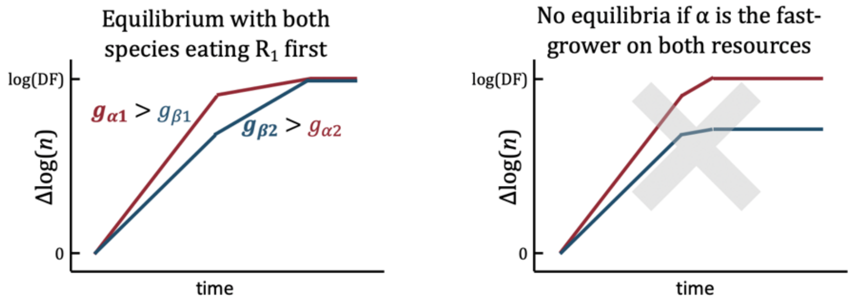


**Supp. Fig. 24.** An equilibrium with both species eating the same resource first (and no lags) requires the fast-grower on the first resource to be the slow grower on the second resource.

We therefore are only interested in continuing to consider the case of $g_{\alpha 1} > g_{\beta 1}$ and $g_{\beta 2} > g_{\alpha 2}$. In this case we know that $R_1$ will run out first because neither species is eating $R_2$ initially. Therefore, for each species to grow by DF on each cycle requires

$$g_{\alpha 1} t_1 + g_{\alpha 2}(t_2 - t_1) = \log(DF) \quad \text{and} \quad g_{\beta 1} t_1 + g_{\beta 2}(t_2 - t_1) = \log(DF)\,.$$

This pair of linear equations is solved for a unique $t_1$ and $t_2$ that can correspond to a fixed point:

$$t_1 = \frac{g_{\beta 2} - g_{\alpha 2}}{g_{\alpha 1} g_{\beta 2} - g_{\beta 1} g_{\alpha 2}} \log(DF) \quad \text{and} \quad t_2 - t_1 = \frac{g_{\alpha 1} - g_{\beta 1}}{g_{\alpha 1} g_{\beta 2} - g_{\beta 1} g_{\alpha 2}} \log(DF)\,.$$

We can now look for the species fractions that exactly finish $R_1$ at $t_1$:

$$n_\alpha(t_1) - n_\alpha(0) + n_\beta(t_1) - n_\beta(0) = s_1$$

$$\frac{f_\alpha}{\mathrm{DF}-1}\left(\exp(g_{\alpha 1} t_1) - 1\right) + \frac{f_\beta}{\mathrm{DF}-1}\left(\exp(g_{\beta 1} t_1) - 1\right) = s_1$$

(Remembering as we set up these equations that the day-to-day carryover means the total population saturates at $\frac{\mathrm{DF}}{\mathrm{DF}-1}$ and starts each day at $\frac{1}{\mathrm{DF}-1}$.)

With the linear constraint that $f_\alpha + f_\beta = 1$, we have two equations that are linear in $f_\alpha$ and $f_\beta$, meaning we will have a unique solution for $f_\alpha$ and $f_\beta$ once $t_1$ is known (which it is). This unique solution is

$$f_\alpha = \frac{s_1(\mathrm{DF}-1) + 1 - \exp(g_{\beta 1} t_1)}{\exp(g_{\alpha 1} t_1) - \exp(g_{\beta 1} t_1)}\,.$$

For this fraction to evaluate as positive ($f_\alpha > 0$) we need $s_1(\mathrm{DF}-1) + 1 > \exp(g_{\beta 1} t_1)$ and for the fraction to evaluate as less than 1 ($f_\alpha < 1$) we need $s_1(\mathrm{DF}-1) + 1 < \exp(g_{\alpha 1} t_1)$ so the existence of this fixed point requires

$$\exp(g_{\beta 1} t_1) < s_1(\mathrm{DF}-1) + 1 < \exp(g_{\alpha 1} t_1)\,,$$

which can be rewritten as

$$\frac{g_{\beta 1}(g_{\beta 2} - g_{\alpha 2})}{g_{\alpha 1} g_{\beta 2} - g_{\beta 1} g_{\alpha 2}} < \frac{\log(\, s_1(\mathrm{DF}-1) + 1\,)}{\log(\mathrm{DF})} < \frac{g_{\alpha 1}(g_{\beta 2} - g_{\alpha 2})}{g_{\alpha 1} g_{\beta 2} - g_{\beta 1} g_{\alpha 2}}\,.$$

Satisfying this expression is necessary and sufficient for the fixed point to exist. The condition is sufficient because we have just shown that with the right initial fractions $R_1$ will run out at the necessary $t_1$, while $R_2$ will then run out at the necessary $t_2$ because both species will have grown by exactly DF and together used up the entire resource supply. (Determining invasibility of monoculture steady-states as a check is straightforward and yields identical requirements.)

Returning to the expression for $f_\alpha$, we can show that $g_{\alpha 1} t_1$ and $g_{\beta 1} t_1$ can be written such that growth rate terms only appear in the ratios $(g_{\beta 1}/g_{\alpha 1})$ and $(g_{\beta 2}/g_{\alpha 2})$:

$$g_{\alpha 1} t_1 = \frac{g_{\alpha 1}(g_{\beta 2} - g_{\alpha 2})}{g_{\alpha 1} g_{\beta 2} - g_{\beta 1} g_{\alpha 2}} \log(\mathrm{DF}) = \frac{(g_{\beta 2}/g_{\alpha 2}) - 1}{(g_{\beta 2}/g_{\alpha 2}) - (g_{\beta 1}/g_{\alpha 1})} \log(\mathrm{DF}) \,.$$

The ability to rewrite everything in terms of the ratios of the growth rates will also carry through to the expression for the existence of the fixed point. Thus, outcomes depend only on $s_1$, DF, and the ratios $(g_{\beta 1}/g_{\alpha 1})$ and $(g_{\beta 2}/g_{\alpha 2})$. Assuming the fixed point is a stable coexistence state (demonstrated below), we can plot competitive outcomes as a function of $(g_{\alpha 1}/g_{\beta 1})$ vs $(g_{\beta 2}/g_{\alpha 2})$ (i.e. α's $R_1$ advantage and β's $R_2$ advantage) at few different dilution factors and supply fractions:

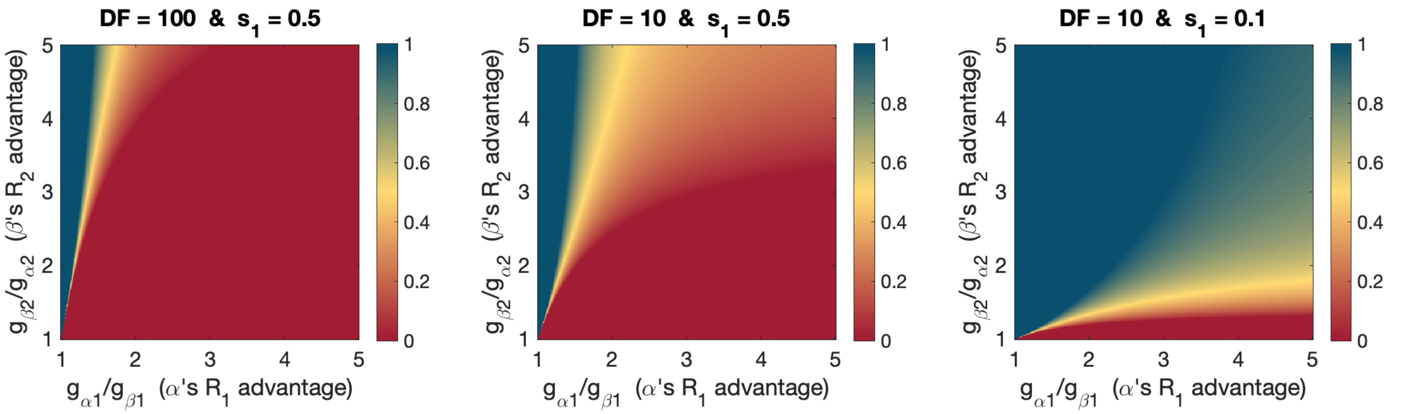


**Supp. Fig. 25.** Competitive outcomes between a two diauxic species with the same resource preference (and no lags). Colormaps shows the population fraction of the slow-grower β. Red represents α excluding β, blue represents β excluding α, and yellow represents coexistence with equal population fractions.

We note that for most conditions (or at least for conditions with $R_1$ and $R_2$ each being roughly equal fractions of the resource supply), the ratio of the growth rates on $R_1$ matters a lot more than the ratio of the growth rates on $R_2$. This is because the species have more log-growth on $R_1$ than on $R_2$. For example, at a dilution factor of 100 and supply fraction of $s_1 = s_2 = 0.5$, the total population size grows approximately 50-fold on $R_1$, but only 2-fold on $R_2$ (left side of Supp. Fig. 26):

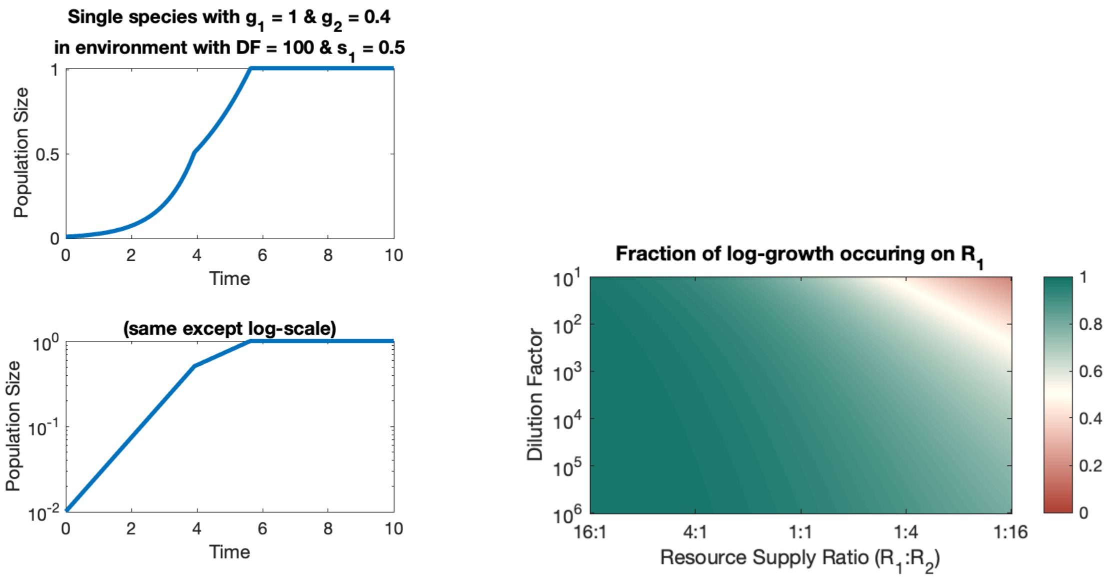


**Supp. Fig. 26.** Most growth happens on the first resource. Left: growth curve for a single species on two resources supplied in equal fractions started at a 1:100 dilution from carrying capacity. On a linear scale, an equal amount of growth appears to happen on both resource, but on a log-scale much more growth happens on the first resource. The time spent growing on each resource is proportional to the log-growth that happens on that resource, so the species grows for longer on the first resource even though both are supplied in equal fractions. In competition the growth that happens on that first resource will tend to have a larger impact than the growth that happens on the second. Right: That most growth happens on the first resource is true at all conditions except small dilution factors and large supply fractions of the second resource.

Returning to the condition for the fixed point to exist,

$$\frac{g_{\beta 1}(g_{\beta 2}-g_{\alpha 2})}{g_{\alpha 1}g_{\beta 2}-g_{\beta 1}g_{\alpha 2}} < \frac{\log(\, s_1(\mathrm{DF}-1)+1\,)}{\log\,(\mathrm{DF})} < \frac{g_{\alpha 1}(g_{\beta 2}-g_{\alpha 2})}{g_{\alpha 1}g_{\beta 2}-g_{\beta 1}g_{\alpha 2}},$$

we note that varying $s_1$ and DF can make $\log(\, s_1(\mathrm{DF}-1)+1\,)\,/\,\log\,(\mathrm{DF})$ arbitrarily close to zero (while remaining positive) or arbitrarily close to one. Therefore, for any set of growth rates $\{g_{\mu i}\}$ such that $g_{\alpha 1} > g_{\beta 1}$, and $g_{\beta 2} > g_{\alpha 2}$, we can find a $\{s_1, \mathrm{DF}\}$ that satisfies either or both halves of the condition for coexistence. Practically, this means that **any α will exclude any β at sufficiently large $s_1$ and DF and any β will exclude any α at sufficiently small $s_1$ and DF with a region of coexistence separating the two regions of competitive exclusions** (in the case of both species eating $R_1$ first, $g_{\alpha 1} > g_{\beta 1}$, and $g_{\beta 2} > g_{\alpha 2}$). This means any dilution factor vs resource supply ratio phase space will look like:

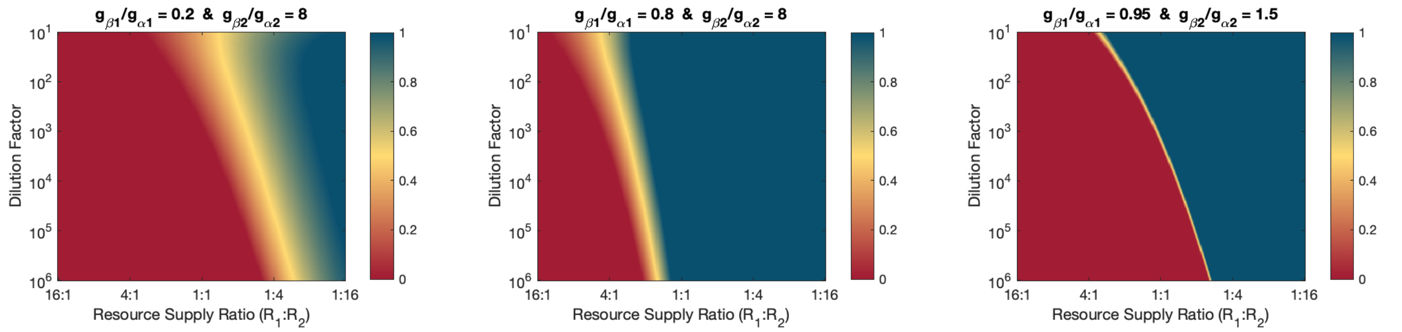


**Supp. Fig. 27** Competitive outcomes between two species with the same resource preference (and no lags). Species become more similar (i.e. growth rate ratios become closer to 1) from left to right across the three plots. Colormap is the steady-state population fraction of the initial slow-grower β. We note that (when the initial slow-grower is the fast-grower on the second resource) there are always regions of each species being excluding and of coexistence. If the same species was the slow-grower on both resources, it would always be competitively excluded by the fast-grower.

**Stability.** We can convince ourselves that this fixed point will be stable by an intuitive argument: If the fraction of the initial fast grower α increases then (i) the overall rate at which the community eats $R_1$ increases and $R_1$ runs out sooner, which hurts α's growth on that cycle, and (ii) the overall rate at which the community eats $R_2$ decreases and the time $(t_2 - t_1)$ increases, which hurts α's growth on that cycle relative to β's growth. Thus increasing α's population fraction above the equilibrium value decreases α's growth over the next dilution cycle. And the same argument can be made in reverse to show that decreasing α's population fraction causes α's growth on the next cycle to increase. This makes the equilibrium stable.[3]

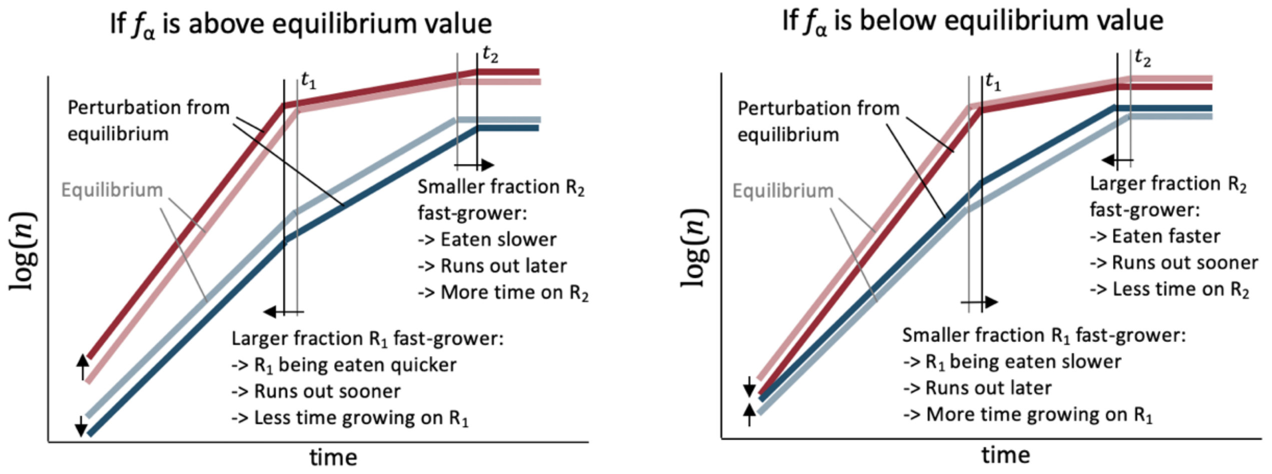


**Supp. Fig. 28.** When the fixed point exists, perturbing species fraction from the fixed point will result in the species fraction converging back towards the fixed point on the next cycle, making the fixed point a stable coexistence state. These cartoons are drawn with $t_2$ becoming earlier or later; this is done for the sake of illustration, but the real effect is that the period $(t_2 - t_1)$ becomes shorter or longer.

One can determine invasibility of monoculture steady states to check that this fixed point exists if and only if both species can invade the other. This is not presented here, but a similar calculation is done below for the case of opposite resource preferences.

While this version of the lag-less diauxie model is producing stable coexistence, we note that it could not have produced unexpected coexistence like what was seen with Aci2 and Pa, because β is the fast-grower on $R_2$ and would outcompete α in an $R_2$ single resource environment.

[3] One more detail is required to make the argument for stability work: If α's population fraction is increased, then even though R1 runs out sooner, α will still have a larger population fraction at this depletion event than it does at equilibrium. This is because β has started the dilution cycle at a lower population fraction and also had less time to grow on R1 meaning that β cannot have a larger population size at the depletion event than it would at equilibrium. (And this means α cannot have a smaller population size than it would at equilibrium because the total population size must still be $1/(DF-1)+s_1$.) That the recovery towards equilibrium cannot have overshot the equilibrium point by the first depletion event is necessary for us to confidently say that R2 will be eaten slower and $(t_2 - t_1)$ increased. The same argument can then be applied to the growth phase on R2 to establish that when perturbed from equilibrium the population fractions will recover back towards the equilibrium point without overshooting it. This means that, in the modeling, there is no chance of an growing, unstable oscillation about the fixed point, nor any other form of oscillation. Experimentally, however, some amount of fluctuation about the fixed point is inevitable due to the noisiness of any real experiment.

#### *Species have opposite resource preferences*

It is often assumed that species having opposite resource preferences is sufficient for coexistence[4], even if they can each eat the other's preferred resource after theirs runs out. From this line of thinking, one might propose that the coexistence between Aci2 and Pa originated not from the tradeoff between Aci2's fast growth and Pa's short lags but instead from the complementarity that Pa initially consumed mostly glutamate while Aci2 initially consumed entirely alanine. It turns out that with Pa being the slow-grower on both single resources and in the two-resource environment, this complementarity in resource preferences is not enough to support coexistence.

If the species have different resource preferences (i.e. if β eats $R_2$ then $R_1$ while α still eats $R_1$ then $R_2$), the resources can run out in either order, yielding two ways of satisfying the requirement that species must grow by the dilution factor on each cycle. We will see below that both these fixed points are possible and that, assuming α is still the initial fast-grower (i.e. $g_{\alpha 1} > g_{\beta 2}$), the fixed point with $R_1$ running out first will be a stable fixed point at which the species coexist and the fixed point with $R_2$ running out first will be an unstable fixed point separating the regions of attraction between bistable outcomes.

**Existence of the stable ($R_1$-first) fixed point.** If $R_1$ runs out first then for both species to grow by the dilution factor, the times at which the resources run out at a fixed point with both species surviving are determined by

$$g_{\alpha 1} t_1 + g_{\alpha 2}(t_2 - t_1) = \log(DF) \quad \text{and} \quad g_{\beta 2} t_2 = \log(DF)\,,$$

yielding,

$$t_1 = \frac{g_{\beta 2} - g_{\alpha 2}}{g_{\alpha 1} - g_{\alpha 2}} \frac{\log\,(\mathrm{DF})}{g_{\beta 2}} \quad \text{and} \quad t_2 = \frac{\log\,(\mathrm{DF})}{g_{\beta 2}}.$$

This calculation assumed that $R_1$ ran out before $R_2$ (i.e. $t_1 < t_2$), and the solution being realistic requires that $R_1$ runs out at a positive time (i.e. $0 < t_1$). These requirements enforce

$$0 < \frac{g_{\beta 2} - g_{\alpha 2}}{g_{\alpha 1} - g_{\alpha 2}} < 1.$$

If we are assuming α is the initial fast-grower ($g_{\alpha 1} > g_{\beta 2}$), this inequality will be satisfied by

$$g_{\alpha 1} > g_{\beta 2} > g_{\alpha 2}\,.$$

This makes intuitive sense because it's simply saying that if α is the initial fast-grower, it needs to become the slow-grower after $R_1$ runs out to give β a chance to catch up:

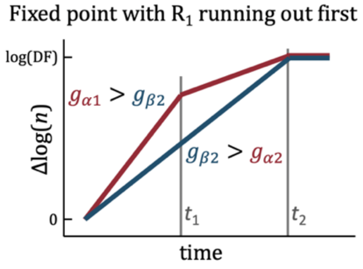


**Supp. Fig. 29.** If $g_{\alpha 1} > g_{\beta 2}$, a fixed point with $R_1$ running out first requires $g_{\beta 2} > g_{\alpha 2}$ so β can catch up to α.

[4] This could be inferred by a cursory reading of Wang et al., although the authors of that paper are clear that species also need to be fast-growers on their preferred resources in the steady-state they demonstrate.

Solving for species fractions is straight-forward using the fact that α eats all of $R_1$ between $t = 0$ and $t = t_1$:

$$\frac{f_\alpha}{\mathrm{DF}-1}\left(\exp(g_{\alpha 1} t_1) - 1\right) = s_1$$
$$f_\alpha = \frac{s_1(\mathrm{DF}-1)}{DF^{\left(\frac{g_{\alpha 1}(g_{\beta 2}-g_{\alpha 2})}{g_{\beta 2}(g_{\alpha 1}-g_{\alpha 2})}\right)} - 1}$$

For DF $> 1$, the above will always produce $f_\alpha > 0$. To have $f_\alpha < 1$ will require

$$\frac{g_{\alpha 1}(g_{\beta 2} - g_{\alpha 2})}{g_{\beta 2}(g_{\alpha 1} - g_{\alpha 2})} > \frac{\log\left(s_1(\mathrm{DF}-1)+1\right)}{\log(\mathrm{DF})}.$$

There will be a fixed point with $R_1$ running out first if and only if $g_{\alpha 1} > g_{\beta 2} > g_{\alpha 2}$ and the above equation is satisfied. (The inequality conditions are sufficient because their being satisfied means a species fractions such that both species grow by log(DF) exists.) Notice that there's a single inequality relating growth rates to environmental parameters compared to the two-sided inequality we had for the coexistence state to exist in the case of species having the same resource preference. This is related to this coexistence region being bordered on one side by a region of the fast-grower excluding the slow-growing but not having a region of the slow-grower excluding the fast-grower on the other side.

This fixed point is stable. For example, if $f_\alpha$ increases from its equilibrium value then $t_1$ must decrease (because a larger population eating $R_1$ finishes it sooner). This causes α to slow down sooner, decreases the time when α is growing faster than β, and increases the time when β is growing faster than α. This returns the system to equilibrium. A similar argument can be made if $f_\alpha$ decreases.

The fraction appearing in the above equation for $f_\alpha$ can be rearranged to be a function of just the ratios $(g_{\beta 2}/g_{\alpha 1})$ and $(g_{\alpha 2}/g_{\beta 2})$, which are the two steps down in the inequality $g_{\alpha 1} > g_{\beta 2} > g_{\alpha 2}$:

$$\frac{g_{\alpha 1}(g_{\beta 2} - g_{\alpha 2})}{g_{\beta 2}(g_{\alpha 1} - g_{\alpha 2})} = \frac{g_{\alpha 1}g_{\beta 2} - g_{\alpha 1}g_{\alpha 2}}{g_{\alpha 1}g_{\beta 2} - g_{\beta 2}g_{\alpha 2}} = \frac{1 - g_{\alpha 2}/g_{\beta 2}}{1 - g_{\alpha 2}/g_{\alpha 1}} = \frac{1 - (g_{\alpha 2}/g_{\beta 2})}{1 - (g_{\beta 2}/g_{\alpha 1})(g_{\alpha 2}/g_{\beta 2})}$$

This rearrangement allows us to plot competitive outcomes across a two dimensional phase space of relative growth rates (under the assumption that the other fixed point does not exist, which can be accomplished with a sufficiently small $g_{\beta 1}$):

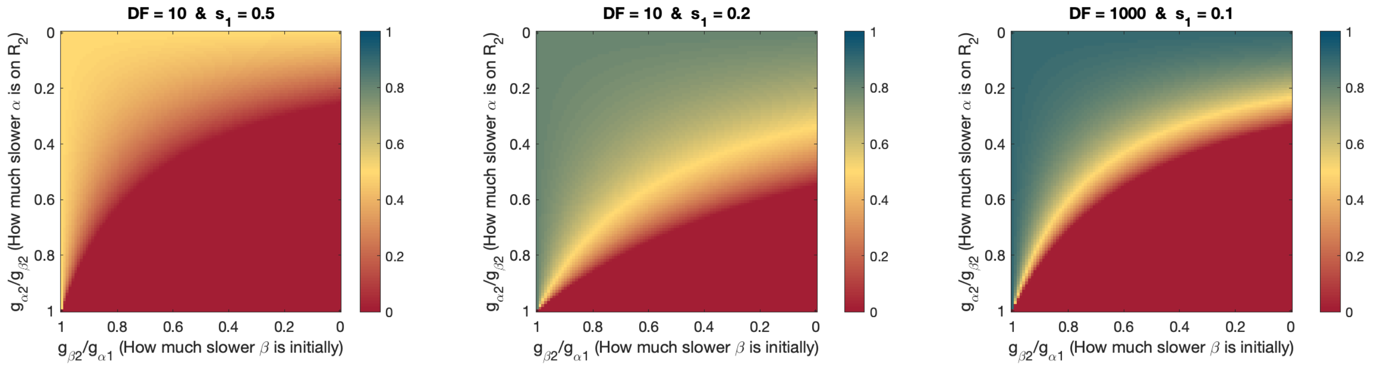


**Supp. Fig. 30.** Competitive outcomes when α eats $R_1$ first and β eats $R_2$ first and growth rates are ordered $g_{\alpha 1} > g_{\beta 2} > g_{\alpha 2} > g_{\beta 1}$. For any environmental parameters, the initial fast-grower α will outcompete β if it is the initial fast-grower by a large enough factor and not too much slower on $R_2$, but the initial slow-grower never outcompetes the initial fast-grower regardless of how much faster it is on the second resource. The population size of any initial slow grower under any environmental conditions is at most the supply concentration of $R_2$. Comparison to Supp. Fig. 25 is also interesting as that figure showed wide parameter regions in which an initial slow-grower with the same resource preference as the fast-grower could outcompete and competitively exclude the fast-grower.

Proceeding to phase spaces across the two environmental parameters, we note that $\log(s_1(\mathrm{DF}-1)+1)\,/\,\log(\mathrm{DF}) \to 0$ when $s_1 \to 0$, meaning that for any fixed dilution factor, the coexistence can be realized at some resource supply ratio. Additionally, $\log(s_1(\mathrm{DF}-1)+1)\,/\log(\mathrm{DF})$ decreases slightly with decreasing DF, so the region in which the fixed point exists will be larger at small dilution factors. We will see below that if $g_{\alpha 1} > g_{\beta 1}$ the other fixed point cannot exist and there is no region in which β excludes α. With $g_{\alpha 1} > g_{\beta 2} > g_{\alpha 2}$ and $g_{\alpha 1} > g_{\beta 1}$, we will therefore have dilution factor vs resource supply phase spaces that look like:

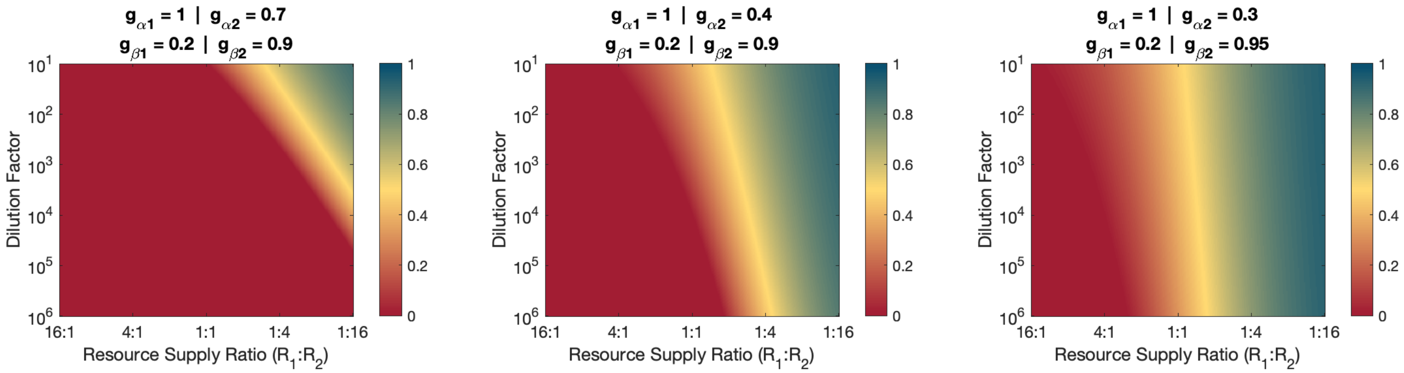


**Supp. Fig. 31.** Competitive outcomes between two species with opposite resource preferences (and no lags). Species α is the initial fast-grower and eats $R_1$ first while β has the opposite resource preference and eats $R_2$ first. Colormap is the steady-state population fraction of the initial slow-grower β. In order for a region of coexistence to exist, α must grow slower than β after it finishes $R_1$ and switches to $R_2$ (i.e. $g_{\alpha 1} > g_{\beta 2} > g_{\alpha 2}$). There is no region of β totally excluding α when both resources are supplied, although β would exclude α if only $R_2$ were supplied in a single-resource competition. This can be compared to Supp. Fig. 27 for similar conclusion as the comparison of Supp. Fig. 30 to Supp. Fig. 25.

**Existence of the unstable ($R_2$-first) fixed point.** If we follow to same procedure to solve for a fixed point with $R_2$ running out first, we get

$$t_2 = \frac{g_{\alpha 1} - g_{\beta 1}}{g_{\beta 2} - g_{\beta 1}} \frac{\log(\mathrm{DF})}{g_{\alpha 1}} \quad \text{and} \quad t_1 = \frac{\log(\mathrm{DF})}{g_{\alpha 1}}.$$

With the above solution the condition $t_2 < t_1$ requires

$$0 < \frac{g_{\alpha 1} - g_{\beta 1}}{g_{\beta 2} - g_{\beta 1}} < 1\,.$$

Species α still being the initial fast grower ($g_{\alpha 1} > g_{\beta 2}$), satisfying the above requires

$$g_{\beta 1} > g_{\alpha 1} > g_{\beta 2}\,.$$

Note that fixed point requires $g_{\beta 1} > g_{\beta 2}$ even though β eats $R_2$ first, giving β what is sometimes called an "anomalous resource preference". That $g_{\beta 1} > g_{\alpha 1}$ is also interesting and means that **there can only be a fixed point with $R_2$ running out first if the initial slow-grower β would have been the initial fast-grower had it chosen $R_1$ instead of $R_2$ as its top preference** (in the case of pure no-lag diauxie, α eating $R_1$ first, β eating $R_2$ first, and $g_{\alpha 1} > g_{\beta 2}$). This growth rate ordering actually makes intuitive sense because it is necessary for β to be able to catch up to α after $R_2$ runs out, as is illustrated in the left panel of Supp. Fig. 32:

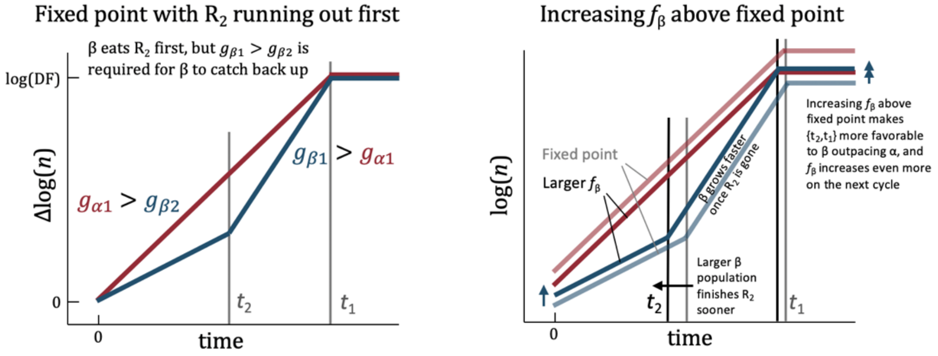


**Supp. Fig. 32.** A fixed point with $R_2$ running out first requires $g_{\beta 1} > g_{\alpha 1} > g_{\beta 2}$ (left), and is unstable (right).

The fixed point with $R_2$ running out first is unstable. This is because increasing β's population fraction above the fixed point causes $R_2$ to run out more sooner, allowing β to speed up sooner and giving β a further increased advantage over α. This is illustrated in the right panel of Supp. Fig. 23. In the other direction, increasing α's population fraction causes $R_2$ to run out later and gives α a further advantage. The characterization of this fixed point as unstable is also confirmed by invasibility calculations in the next subsection.

The species fractions at which the unstable fixed point occurs can be calculated in the same way we calculated the species fractions for the stable fixed point. The result is

$$f_\beta = \frac{s_2(\mathrm{DF}-1)}{DF^{\left(\frac{g_{\beta 2}(g_{\beta 1}-g_{\alpha 1})}{g_{\alpha 1}(g_{\beta 1}-g_{\beta 2})}\right)}-1}.$$

And the necessary and sufficient condition for the fixed point to exist is

$$g_{\beta 1} > g_{\alpha 1} > g_{\beta 2} \quad \text{and} \quad \frac{g_{\beta 2}(g_{\beta 1}-g_{\alpha 1})}{g_{\alpha 1}(g_{\beta 1}-g_{\beta 2})} > \frac{\log\left((1-s_1)(\mathrm{DF}-1)+1\right)}{\log(\mathrm{DF})}.$$

Notice that the RHS is now smallest as $s_1 \to 1$, meaning that, while the stable fixed point occurred at small $s_1$ and large $s_2$, the unstable fixed point occurs at large $s_1$ and small $s_2$.

If we choose $g_{\beta 1} > g_{\alpha 1} > g_{\alpha 2} > g_{\beta 2}$, the unstable fixed point can exist but the stable fixed point cannot. Plotting the bifurcation diagram for a changing resource supply ratio:

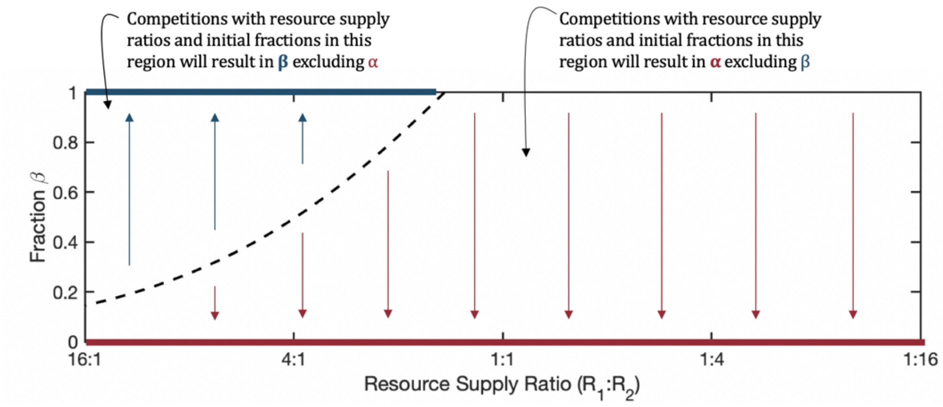


**Supp. Fig. 33.** Example bifurcation diagram for bistability between Pa. Calculated for DF = 10, $g_{\beta 1} = 1$, $g_{\alpha 1} = 0.6$, $g_{\alpha 2} = 0.55$, and $g_{\beta 2} = 0.5$.

The bistability occurs at large supplies of $R_1$ and small supplies of $R_2$. This is because a small supply fraction $s_2$ leads to $R_2$ running out sooner and β getting to speed up sooner. The bistability occurs because an increased fraction $f_\beta$ causes $R_2$ to run out sooner, further benefitting β, and creating a positive feedback loop. The unstable fixed point occurs with $R_2$ running out first and not at the point at which both resources run out at the same time because β needs some minimum time growing on $R_1$ to catch back up to α.

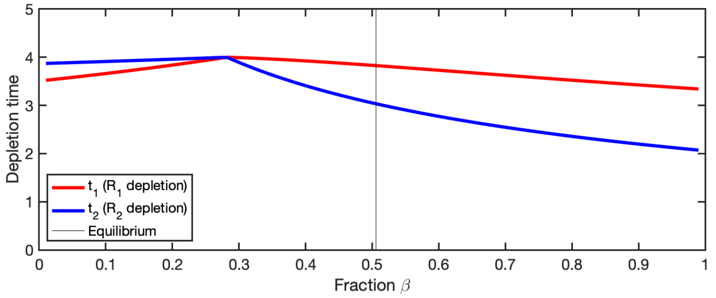


**Supp. Fig. 34.** Resource depletion times vary with initial fraction $f_\beta$. The unstable equilibrium is $f_\beta = 0.4943$ with $R_2$ running out first. Calculated for $s_1 = 0.8$ (i.e. 4:1 $R_1$:$R_2$). Same dilution factor and growth rates as Supp. Fig. 24. (DF = 10, $g_{\beta 1} = 1$, $g_{\alpha 1} = 0.6$, $g_{\alpha 2} = 0.55$, and $g_{\beta 2} = 0.5$)

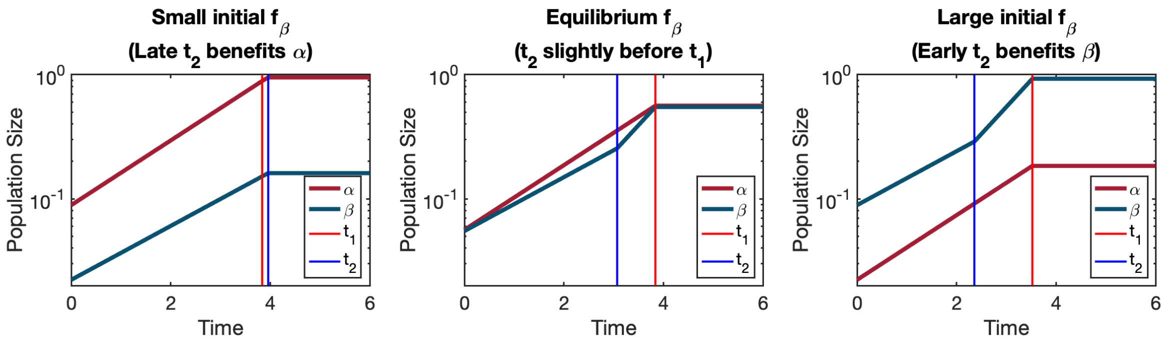


**Supp. Fig. 35.** Population sizes over the course of one dilution cycle for different initial $f_\beta$. At small $f_\beta$, $R_1$ runs out first, so β never grows on $R_2$ and never has a chance to catch back up to α, leading to α further increasing its species fraction and over additional dilution cycles eventually competitively excluding β. By contract, at large $f_\beta$, $R_2$ runs out first, so β has a chance to catch back up to and even overtake α, eventually leading to β competitively excluding α. Calculated for same parameters as Supp. Fig. 25 ($s_1 = 0.8$ , DF = 10, $g_{\beta 1} = 1$, $g_{\alpha 1} = 0.6$, $g_{\alpha 2} = 0.55$, and $g_{\beta 2} = 0.5$).

If $g_{\beta 1} > g_{\alpha 1} > g_{\beta 2} > g_{\alpha 2}$, both the stable and the unstable fixed point are possible. This is can be seen by lowering the value of $g_{\alpha 2}$ from the Supp. Fig. 33–35 example to be below $g_{\beta 2}$:

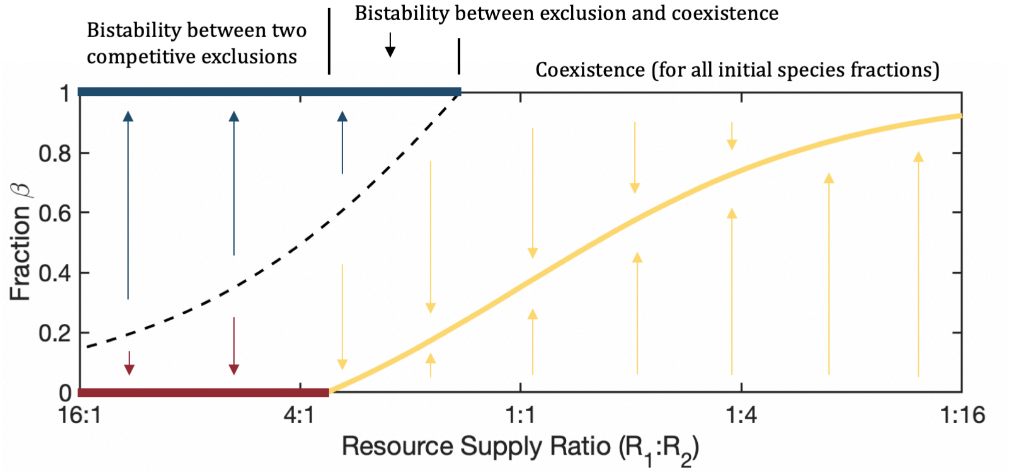


**Supp. Fig. 36.** Bifurcation diagram for DF = 10, $g_{\beta 1} = 1$, $g_{\alpha 1} = 0.6$, $g_{\beta 2} = 0.5$, and $g_{\alpha 2} = 0.2$.

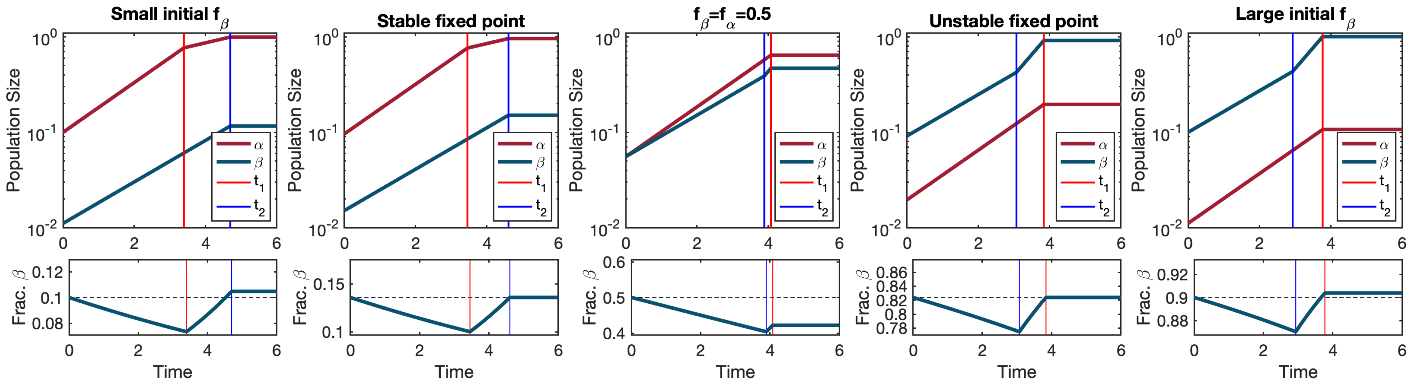


**Supp. Fig. 37.** Population sizes (top) and fraction β (bottom) over the course of one dilution cycle for different initial $f_\beta$. Calculated for $s_1 = 2/3$ (2:1 $R_1$:$R_2$) and same growth rates and dilution factor as Supp. Fig. 36 (DF = 10, $g_{\beta 1} = 1$, $g_{\alpha 1} = 0.6$, $g_{\beta 2} = 0.5$, and $g_{\alpha 2} = 0.2$). In the bottom row of plots, the y-axis scale is different in each plot.

The intuition behind the bistability between β excluding α and coexistence is as follows: Because α slows down when $R_1$ runs out and β speeds up when $R_2$ runs out, β performs best when the time between resource depletions is greatest. This occurs at very small and very large fractions $f_\beta$ (because at these fractions there is a much larger population of the species eating one resource than the species eating the other). Conversely, β performs worse at roughly equal species fractions when the two resources run out at roughly the same time. With increasing $f_\beta$, these two trends create a region of $f_\beta$ increasing, a region of $f_\beta$ decreasing, and then a region of $f_\beta$ increasing. Continuity places a stable and then an unstable fixed point in between these two regions.

It is interested to note that **when $g_{\beta 1} > g_{\alpha 1} > g_{\beta 2} > g_{\alpha 2}$ β would exclude α in both single resource environments (because $g_{\beta 1} > g_{\alpha 1}$ and $g_{\beta 2} > g_{\alpha 2}$), but α can survive and even exclude β in the two-resource environment** (with α eating $R_1$ first and β eating $R_2$ first). This is the closest thing to "unexpected coexistence" possible in the simple diauxic model without lags, but even if α survives without being the fast-grower in either single-resource environment it is still the fast-grower in two-resource environment. This is also a very strange and particular case whereas coexistence derived from diauxic lags is much more plausible to be a common phenomenon.

Dilution factor vs resource supply ratio phase spaces can be calculated:

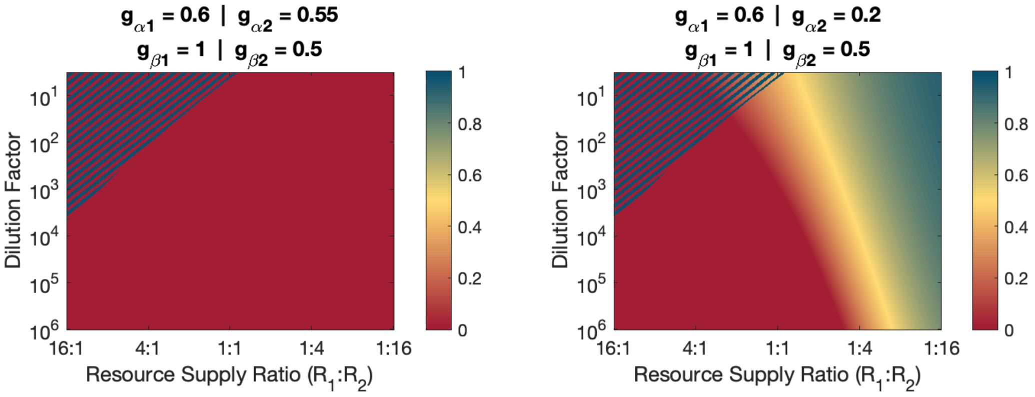


**Supp. Fig. 38.** Dilution factor vs resource supply ratio phase spaces. Colormap is β's population fraction with striped regions indicating bistability. Left plot has same growth rates as Supp. Fig. 33–35. Right has same growth rates as Supp. Fig. 36–37.

(When a set of growth rates allows for both the stable fixed point and the unstable fixed point, it may or may not be the case that there exists a dilution factor and resource supply ratio combination at which both fixed point are realized.)

The last question to ask in the case of having both stable and unstable fixed points is whether the unstable fixed point always occurs at a larger $f_\beta$ than the stable fixed point occurs at (with fixed growth rates and environmental parameters). The answer is yes with a simple explanation. At the unstable fixed point, β *eats all* of $R_2$ so its population fraction is at least $f_{\beta|\text{unstable pt.}} > s_2$. At the stable fixed point, β *only eats* $R_2$ so β's population fraction cannot be larger than $f_{\beta|\text{stable pt.}} < s_2$. Therefore, $f_{\beta|\text{unstable pt.}} > f_{\beta|\text{stable pt.}}$.

**Invasibility as a check of the above calculations.** We can use invasibility as a check of the above solutions for fixed points in the case of α eating $R_1$ first and β eating $R_2$ first and as check of our characterizations of the points as stable and unstable.

Let's first consider α's monoculture steady state and β's ability to invade it. We can calculate when $R_1$ must run out from the condition that α exactly finishes $R_1$:

$$\frac{1}{\text{DF}-1}\left(\exp\left(g_{\alpha 1} t_{1|\alpha}\right)-1\right) = s_1$$

$$t_{1|\alpha} = \frac{1}{g_{\alpha 1}}\log\left(s_1(\text{DF}-1)+1\right)$$

And for α to grow by DF, we need:

$$g_{\alpha 1} t_{1|\alpha} + g_{\alpha 2}\left(t_{2|\alpha} - t_{1|\alpha}\right) = \log(\text{DF})$$

$$t_{2|\alpha} = \frac{1}{g_{\alpha 2}}\left(\log(\text{DF}) - (g_{\alpha 1} - g_{\alpha 2})t_{1|\alpha}\right)$$

$$t_{2|\alpha} = \frac{1}{g_{\alpha 2}}\log(\mathrm{DF}) - \frac{g_{\alpha 1} - g_{\alpha 2}}{g_{\alpha 1}\, g_{\alpha 2}}\log(s_1(\mathrm{DF}-1)+1)$$

From this we determine that β can invade α's monoculture steady-state iff:

$$g_{\beta 2} t_{2|\alpha} > \log(\mathrm{DF})$$

$$g_{\beta 2}\left(\frac{1}{g_{\alpha 2}}\log(\mathrm{DF}) - \frac{g_{\alpha 1} - g_{\alpha 2}}{g_{\alpha 1}\, g_{\alpha 2}}\log(s_1(\mathrm{DF}-1)+1)\right) > \log(\mathrm{DF})$$

This rearranges to:

$$g_{\beta 2}(g_{\alpha 1} - g_{\alpha 2})\log(s_1(\mathrm{DF}-1)+1) < g_{\alpha 1}(g_{\beta 2} - g_{\alpha 2})\log(\mathrm{DF})$$

In order for β to invade α it must grow faster than α at some point during the dilution cycle. Because $g_{\alpha 1} > g_{\beta 2}$, this requires $g_{\beta 2} > g_{\alpha 2}$. This resolves issues with otherwise unknown signs and allows simplification to:

$$\beta \text{ invades } \alpha \quad \text{iff} \quad \frac{\log(s_1(\mathrm{DF}-1)+1)}{\log(\mathrm{DF})} < \frac{g_{\alpha 1}(g_{\beta 2} - g_{\alpha 2})}{g_{\beta 2}(g_{\alpha 1} - g_{\alpha 2})}.$$

This is exactly the condition for the existence of a fixed point with $R_1$ running out. Therefore, β can invade α if and only if there exist a fixed point with $R_1$ running out first.

We now move on to considering β's ability to invade α's steady state. Following the same calculations:

$$t_{2|\beta} = \frac{1}{g_{\beta 2}}\log(s_2(DF-1)+1)$$

$$t_{1|\beta} = \frac{1}{g_{\beta 1}}\log(\mathrm{DF}) - \frac{g_{\beta 2} - g_{\beta 1}}{g_{\beta 1}\, g_{\beta 2}}\log(s_2(DF-1)+1)$$

From these depletion times α can invade β's monoculture steady state if and only if:

$$g_{\alpha 1}\left(\frac{1}{g_{\beta 1}}\log(\mathrm{DF}) - \frac{g_{\beta 2} - g_{\beta 1}}{g_{\beta 1}\, g_{\beta 2}}\log(s_2(DF-1)+1)\right) > \log(DF)\,.$$

And this can be rearrange to be exactly the opposite of the condition for the $R_2$-first fixed point to exist. Therefore, α can invade β if and only if there does not exist a fixed point with $R_2$ running out first.

Summarizing, the linking of the conditions for the existence of the fixed points to the conditions invasibility exactly agrees with the necessary parity and confirms our characterizations of them as stable and unstable:

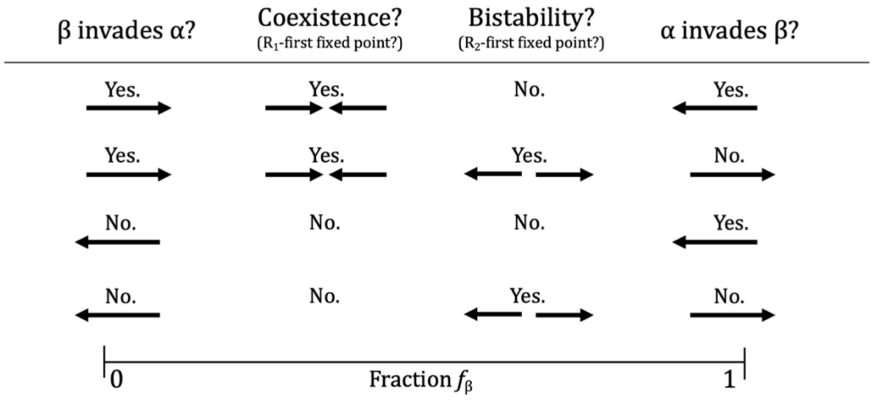


**Supp. Fig. 39.** All allowed combinations of fixed point existence and invasibility.

#### *Five qualitatively distinct resource ratio vs dilution factor competitive outcome phase spaces for two simple-diauxic species*

At this point we have indexed all possible cases of the simple diauxie model of two species on two resources without lags. We summarize the results below:

| Phase Space | Description | Growth Rate Orderings if Same Preference | Growth Rate Orderings if Opposite Preference |
|---|---|---|---|
| 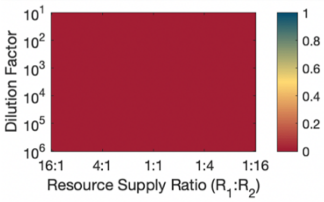 | Fast-grower α always excludes β | $g_{\alpha 1} > g_{\beta 1}$<br>$g_{\alpha 2} > g_{\beta 2}$ | $g_{\alpha 1} > g_{\beta 1}$<br>$g_{\alpha 1} > g_{\beta 2}$<br>$g_{\alpha 2} > g_{\beta 2}$ |
| 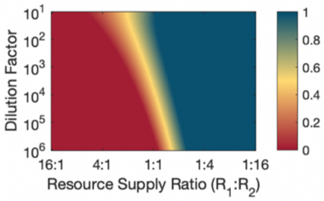 | Regions of coexistence and of each species excluding the other | $g_{\alpha 1} > g_{\beta 1}$<br>$g_{\beta 2} > g_{\alpha 2}$ | *not possible* |
| 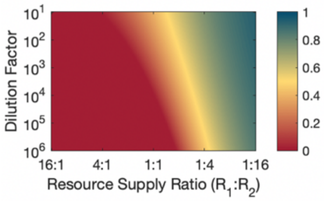 | Regions of coexistence and of fast-grower α excluding β, but not region of β excluding α | *not possible* | $g_{\alpha 1} > g_{\beta 2} > g_{\alpha 2}$<br>$g_{\alpha 1} > g_{\beta 1}$ |
| 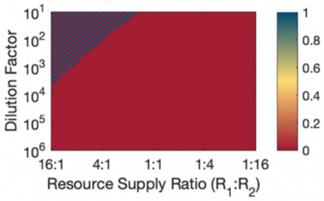 | Regions of fast-grower α excluding β and of bistability between competitive exclusions | *not possible* | $g_{\beta 1} > g_{\alpha 1} > g_{\alpha 2} > g_{\beta 2}$ |
| 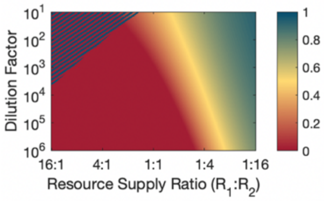 | Regions of coexistence, of fast-grower α excluding β, and of bistability (with or without coex/bi overlap) | *not possible* | $g_{\beta 1} > g_{\alpha 1} > g_{\beta 2} > g_{\alpha 2}$ |

**Supp. Fig. 40.** All qualitatively distinct dilution factor vs resource supply ratio competitive outcome phase spaces for two simple-diauxic species (no lags) on two resources. Dilution factor and resource supply ratio are chosen as the axes for this summary as these are (i) environmental variables that can be easily tuned experimentally than growth rates and (ii) equal relevant for all the different cases where each case has a different set of relevant growth rate ratios. Varying the resource supply ratio primarily affects outcomes via changing the relative timing of when the resources run out. Varying the dilution factor primarily affects outcomes via changing how much growth happens before either resource has run out yet.

Notably, any species the survives in any of the above cases must be the fast-grower on at least one resource or the fast-grower in the two-resource environment. Simple diauxie on two resources without lags therefore cannot explain the survival of a consistent slow-grower.

#### *Nonequal yields*

We now implements yields $\{y_{\mu i}\}$ such that if species $\mu$ eats one unit of resource *i*, it gains $y_{\mu i}$ units of population size. In the order direction, this means that if species $\mu$ has grown by one unit of population size on resource *i* it has consumed $\frac{1}{y_{\mu i}}$ units of resource *i*.

For brevity, we will only consider the case of species having the same resource preference. At a fixed point it is still the case that

$$g_{\alpha 1}t_1 + g_{\alpha 2}(t_2 - t_1) = \log(DF) \quad \text{and} \quad g_{\beta 1}t_1 + g_{\beta 2}(t_2 - t_1) = \log(DF)\,.$$

And, these equations are solved by

$$t_1 = \frac{g_{\beta 2} - g_{\alpha 2}}{g_{\alpha 1}g_{\beta 2} - g_{\beta 1}g_{\alpha 2}}\log(DF) \quad \text{and} \quad t_2 - t_1 = \frac{g_{\alpha 1} - g_{\beta 1}}{g_{\alpha 1}g_{\beta 2} - g_{\beta 1}g_{\alpha 2}}\log(DF).$$

The calculations for the population sizes are now more complicated, however. In order to solve for the population sizes at the fixed point we would need to use constraints on both $R_1$ being finished at $t_1$ and $R_2$ being finished at $t_2$:

$$\frac{n_\alpha}{\mathrm{DF}}\frac{(\exp(g_{\alpha 1}t_1) - 1)}{y_{\alpha 1}} + \frac{n_\beta}{\mathrm{DF}}\frac{\left(\exp\left(g_{\beta 1}t_1\right) - 1\right)}{y_{\beta 1}} = s_1$$

$$\frac{n_\alpha \exp(g_{\alpha 1}t_1)}{\mathrm{DF}}\frac{(\exp(g_{\alpha 2}(t_2 - t_1)) - 1)}{y_{\alpha 2}} + \frac{n_\beta \exp\left(g_{\beta 1}t_1\right)}{\mathrm{DF}}\frac{\left(\exp\left(g_{\beta 2}(t_2 - t_1)\right) - 1\right)}{y_{\beta 2}} = s_2$$

These equations will have a messy solution (and especially so once $t_1$ and $t_2$ are plugged in), but note that they are linear in $n_\alpha$ and $n_\beta$, so there will be at most one fixed point. This allows us to proceed with invasibility criteria without having to worry about having multiple fixed points.

Considering α's monoculture steady-state: α's steady-state population size will be

$$n_{\alpha|\alpha} = \frac{\mathrm{DF}}{\mathrm{DF} - 1}(y_{\alpha 1}s_1 + y_{\alpha 2}s_2)$$

That α finishes $R_1$ at time $t_{1|\alpha}$ yields:

$$\frac{n_{\alpha|\alpha}}{\mathrm{DF}}\left(\exp\left(g_{\alpha 1}t_{1|\alpha}\right) - 1\right) = y_{\alpha 1}s_1$$

$$t_{1|\alpha} = \frac{1}{g_{\alpha 1}}\log\left(\frac{y_{\alpha 1}s_1 DF + n_{\alpha|\alpha}}{n_{\alpha|\alpha}}\right)$$

$$t_{1|\alpha} = \frac{1}{g_{\alpha 1}}\log\left(\frac{y_{\alpha 1}s_1(DF - 1) + (y_{\alpha 1}s_1 + y_{\alpha 2}s_2)}{(y_{\alpha 1}s_1 + y_{\alpha 2}s_2)}\right)$$

And that α has matched the dilution factor at time $t_{2|\alpha}$ yields:

$$g_{\alpha 1}t_{1|\alpha} + g_{\alpha 2}\left(t_{2|\alpha} - t_{1|\alpha}\right) = \log(\mathrm{DF})$$

$$t_{2|\alpha} - t_{1|\alpha} = \frac{1}{g_{\alpha 2}}\log(\mathrm{DF}) - \frac{g_{\alpha 1}}{g_{\alpha 2}}t_{1|\alpha}$$

β's ability to invade α's monoculture steady state is determined by:

$$g_{\beta 1}t_{1|\alpha} + g_{\beta 2}\left(t_{2|\alpha} - t_{1|\alpha}\right) > \log(\mathrm{DF})$$

$$g_{\beta 1}t_{1|\alpha} + g_{\beta 2}\left(\frac{1}{g_{\alpha 2}}\log(\mathrm{DF}) - \frac{g_{\alpha 1}}{g_{\alpha 2}}t_{1|\alpha}\right) > \log(\mathrm{DF})$$
$$\left(g_{\beta 1}\, g_{\alpha 2} - g_{\alpha 1}g_{\beta 2}\right) t_{1|\alpha} > \left(g_{\alpha 2} - g_{\beta 2}\right)\log(\mathrm{DF})$$
$$\log\left(\frac{y_{\alpha 1}s_1(DF-1) + (y_{\alpha 1}s_1 + y_{\alpha 2}s_2)}{(y_{\alpha 1}s_1 + y_{\alpha 2}s_2)}\right)/\log(\mathrm{DF}) < \frac{g_{\alpha 1}\left(g_{\beta 2} - g_{\alpha 2}\right)}{g_{\alpha 1}g_{\beta 2} - g_{\beta 1}\, g_{\alpha 2}}$$

If the above is satisfied β can invade α. And a similar expression will exist for whether α can invade β.

At this point, we now note that all the yield terms except one are redundant as far as invasibility is concerned. As we can see above all that β's ability to invade α actually depends on is the ratio $y_{\beta 2}/y_{\beta 1}$. Furthermore, scaling both $y_{\alpha 2}$ and $y_{\beta 2}$ by the same factor essentially just rescales the resource supply. (Both species having their yield on a resource doubled has the same effect as doubling the supply.) So we can simplify to:

$$\beta \text{ invades } \alpha \quad \text{iff} \quad \frac{\log(s_1(DF-1)+1)}{\log(\mathrm{DF})} > \frac{g_{\alpha 1}(g_{\beta 2} - g_{\alpha 2})}{g_{\alpha 1}g_{\beta 2} - g_{\beta 1}g_{\alpha 2}}$$

$$\text{and} \quad \alpha \text{ invades } \beta \quad \text{iff} \quad \frac{\log\left(\frac{s_1(DF-1) + \left(s_1 + y_{\beta 2}s_2\right)}{\left(s_1 + y_{\beta 2}s_2\right)}\right)}{\log(\mathrm{DF})} < \frac{g_{\beta 1}\left(g_{\beta 2} - g_{\alpha 2}\right)}{g_{\alpha 1}g_{\beta 2} - g_{\beta 1}g_{\alpha 2}}$$

When $y_{\beta 2}$ is large (i.e. when β has a larger yield on the second resource relative to α than it does on the first resource), a bistability is possible:

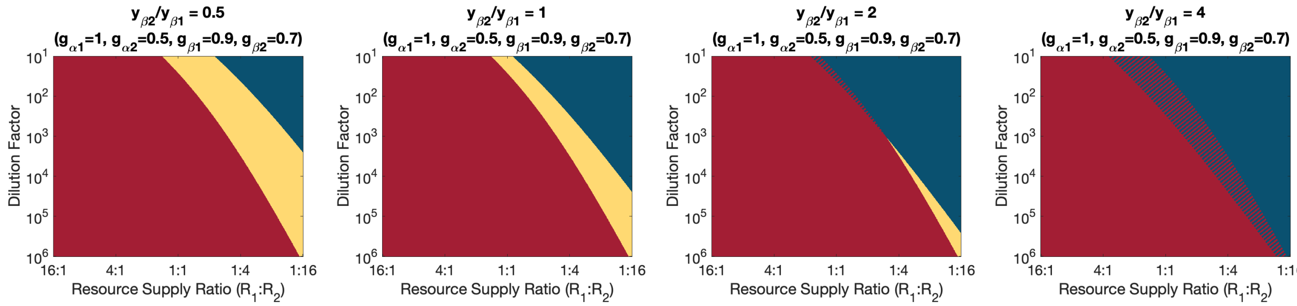


**Supp Fig. 41.** A bistability resulting from species having different yields on each resource. As the slow-grower has its relative yield on the second resource (its lower preference) increased, a coexistence region becomes a bistability.

With yields implemented this new bistability emerges, but it remains the case that species must be the fast-grower in at least one environment to survive.

#### *Separate two-resource co-utilizations states*

This creates more permutations and possibilities than is practical to analyze here. Two key points are:

- A species can easily be excluded in both single-resource environments but survive in the two-resource environment if it has small single-resource growth rates but a large two-resource growth rate.
- But, to survive in competition a species must still be the fast-grower in one single-resource environment or the two-resource environment or else its competitor will always be growing faster than it.

#### *Continuity of the day-to-day map*

Much of the analysis perform in this Supplemental Information relies on the day-to-day map being continuous in order to maintain strict mathematically validity. This section provides a proof that the model has the necessary continuity.

This proof will be done for an arbitrary number of species $N_{\text{sp}} \geq 1$ and an arbitrary number of resources $N_{\text{re}} \geq 1$ and initially for the case of no diauxic lags. Diauxic lags will be added after the no-lags case has been completed.

To avoid some difficult edge cases, we assume that (i) at any point in time there is at least one surviving species (i.e. $\exists \mu : n_\mu > 0$), (ii) each resource has at least one surviving species that eats it (i.e. $\forall i\ \exists \mu : n_\mu > 0\ \&\ g_{\mu i} > 0$), and that (iii) no species prefers a resource it does not grow on to a resource it can grow on (i.e. if μ consumes $R_i$ before $R_j$ then $g_{\mu j} > 0 \rightarrow g_{\mu i} > 0$.) These assumptions guarantee that there is always at least one resource that is being eaten and that eventually all resources are depleted.

At any point during the course of a day, the dynamics can be written in the form of dynamic equations,

$$\frac{d}{dt}\boldsymbol{n} = \boldsymbol{G}(\boldsymbol{c} > 0) \cdot \boldsymbol{n} \quad \text{and} \quad \frac{d}{dt}\boldsymbol{c} = -\boldsymbol{H}(\boldsymbol{c} > 0) \cdot \boldsymbol{n}\,,$$

where $\boldsymbol{G}$ is a diagonal matrix that depends on which resources are left (i.e. which $c_i > 0$) and whose elements are each species' growth rate on its top-preference remaining resources and $\boldsymbol{H}$ is a matrix that has elements

$$\boldsymbol{H}_{\mu i} = \begin{cases} g_{\mu i} & \text{if species μ is eating } \mathrm{R}_i \\ 0 & \text{else} \end{cases}.$$

Because the dynamic equations have discontinuities every time a resource is depleted, it would seem likely that the day-to-day would be discontinuous. Fortunately, this is not the case, as is proven below.

**For any period of time during which no additional resource are depleted,** $\boldsymbol{G}$ and $\boldsymbol{H}$ are constant and the dynamics are therefore smooth and continuous. During this time the system moves through a population-size-and-resource-concentration subspace $S_{(\boldsymbol{c}>0)}$,

$$\begin{pmatrix} \boldsymbol{n}(t) \\ \boldsymbol{c}(t) \end{pmatrix} \in \left\{\mathbb{R}^{(N_{\text{sp}}+N_{\text{re}})}\ :\ c_i(t) > 0 \text{ if } \mathrm{R}_i \text{ hasn't run out yet, else } c_i(t) = 0\right\} \equiv S_{(\boldsymbol{c}>0)}$$

where the set of $\mathrm{R}_i$ that haven't run out yet is constant while in $S_{(\boldsymbol{c}>0)}$.

(The notation $S_{(\boldsymbol{r}>0)}$ is used to reference the subspace being different for each combination of which terms in $\boldsymbol{c}(t)$ are still greater than zero. For example, subspace $S_{111}$ would correspond to no resources having been depleted yet, while subspace $S_{010}$ would correspond to $R_1$ and $R_3$ having been depleted.)

Because dynamics in subspace $S_{(\boldsymbol{c}>0)}$ are smooth and continuous, while the system trajectory remains in this space $\boldsymbol{n}(t)$ and $\boldsymbol{c}(t)$ vary smoothly and continuously with the population sizes and resource concentrations time when $S_{(\boldsymbol{c}>0)}$ was entered (as well as with growth rates $\{g_{\mu i}\}$ and the time when $S_{(\boldsymbol{c}>0)}$ was entered).

Another resource is depleted and the subspace $S_{(\boldsymbol{c}>0)}$ is left when the boundary defined by $\{\exists i : c_i(t) = 0 \text{ for } \mathrm{R}_i \text{ that hadn't run out yet}\}$ is hit. This boundary is continuous and piece-wise smooth, being made up of a set of orthogonal planes in resource-concentration space.

For each boundary plane, if it's the planes defined by $c_i = 0$ then the distance to the plane is $c_i(t)$, which varies smoothly and continuously with parameters as noted above and is monotonically decreasing because $\frac{d}{dt}\boldsymbol{c} = -\boldsymbol{H}(\boldsymbol{c} > 0) \cdot \boldsymbol{n}$ with all $\boldsymbol{H}_{\mu i} \geq \boldsymbol{0}$ (with the distance to at least one of the planes strictly monotonically decreasing because at least one resource is being eaten). The distance to the closest plane therefore varies continuously and is strictly monotonically decreasing with time. Thus, the time at which the boundary of $S_{(\boldsymbol{c}>0)}$ is hit and the next resource is depleted varies continuously with the population sizes and resource concentrations when $S_{(\boldsymbol{c}>0)}$ was entered (by the invertibility of strictly monotonic functions). Therefore:

**The time at which the next resource is depleted and the population sizes and resource concentrations when that happens vary continuously with the time at which the previous resource was depleted (if applicable) and the population sizes and resource concentrations when that previous resource was depleted (or the population sizes at the start of the day if no previous resource has run out yet).** This statement is true even across changes to which resource ran out previously and which resource runs out next.

**When the next resource is depleted,** a new subspace $S_{(\boldsymbol{c}>0)'}$ is entered. Under a continuous variation to the population sizes at the start of the day, if the resource depletion order does not change, the sequence of subspaces that are entered (e.g. $\langle S_{1111}, S_{1101}, S_{1001}, S_{1000}\rangle$) does not change. Therefore, the system passes through a well-defined series of subspaces that each have a continuous mapping of population sizes and resource concentrations upon entry to population sizes and resource concentration upon exit. A sequence of continuous mappings is itself a continuous map. The implementation of the dilution is also another continuous map. Therefore, if the resource depletion order does not change, the mapping from the start of one day to the start of the next is a continuous map. (Below we proceed to prove that this true even if the depletion order changes.)

Furthermore, **if the resource depletion order does not change, the mapping from the start of one day to the start of the next is a smooth and continuous map.** We can establish this by going back through the argument and noting that (i) if we know which boundary plane we're going to hit, we only care about the distance to that plane, and that distance is *smoothly* and strictly monotonically decreasing, (ii) the time at which we hit the plane is now smoothly varying with relevant quantities (by the smooth invertibility of smooth and strictly monotonic functions), (iii), because the plane itself is smooth, the location at which the plane is hit (in population-size-and-resource-concentration space) varies smoothly with the population size and resource concentration at the previous resource depletion (or start of the day), and (iv) that we now have sequence of smooth and continuous maps, which is itself a smooth and continuous map.

**If a continuous variation to population sizes cause the resource depletion order to change,** the sequence of spaces through which the system trajectory passes can change, which breaks the arguments in the previous two paragraphs. There must be at least two resources for this to happen; we'll assume they're $R_1$ and $R_2$. Before either of $R_1$ and $R_2$ have been depleted the system is travelling through subspace $S_{11}$ (which could also be the sequence of multiple subspaces). If $R_1$ is depleted first the system next enters subspace $S_{01}$, and if $R_2$ is depleted first the system next enters subspace $S_{10}$. After both $R_1$ and $R_2$ are depleted, the system enters subspace $S_{00}$ (which could also simply be the end of the day). If $R_1$ and $R_2$ are depleted simultaneously, the system enters subspace $S_{00}$ directly from $S_{11}$.

Suppose we start with some initial (i.e. start-of-day) species fractions $\boldsymbol{f}$ and start varying them from values that will result in $R_1$ running out before $R_2$ to values that result in $R_2$ running out before $R_1$. Because we can choose $\boldsymbol{f}$ from a continuum of values, we can choose an $\boldsymbol{f}_{12}$ such that $R_1$ runs out before $R_2$ and an $\boldsymbol{f}_{21}$ such that $R_2$ runs out before $R_1$ that are arbitrarily close to each other (i.e. each just across the boundary separating which resource out first from each other). Mathematically,

$$\forall \epsilon\ \exists\, \boldsymbol{f}_{12}, \boldsymbol{f}_{21} :\ \|\boldsymbol{f}_{12} - \boldsymbol{f}_{21}\| < \epsilon \ \text{ and } \mathrm{R}_i \text{ runs out before } \mathrm{R}_j \text{ when starting from } \boldsymbol{f}_{ij}\,.$$

The mapping from one day to the next is continuous across this change in depletion order iff as the initial fractions converge (i.e. as $\|\boldsymbol{f}_{12} - \boldsymbol{f}_{21}\| \to 0$) the population sizes at the end of the day given each initial fractions also converge (i.e. $\left\|\boldsymbol{n}_{|\boldsymbol{f}_{12}}(t_{\mathrm{sat}|\boldsymbol{f}_{12}}) - \boldsymbol{n}_{|\boldsymbol{f}_{21}}(t_{\mathrm{sat}|\boldsymbol{f}_{21}})\right\| \to 0$). This can be proven:

By the continuity within subspace $S_{11}$ that was shown above: As we choose $\boldsymbol{f}_{12}$ and $\boldsymbol{f}_{21}$ that become arbitrarily close (i.e. as $\|\boldsymbol{f}_{12} - \boldsymbol{f}_{21}\| \to 0$), the times at which subspace $S_{11}$ is left given each $\boldsymbol{f}$ converge (i.e. $\left|t_{1|\boldsymbol{f}_{12}} - t_{2|\boldsymbol{f}_{21}}\right| \to 0$). As this happens the population sizes and resource concentrations when subspace $S_{11}$ is left under each trajectory also converge (i.e. $\left\|\boldsymbol{n}_{|\boldsymbol{f}_{12}}(t_{1|\boldsymbol{f}_{12}}) - \boldsymbol{n}_{|\boldsymbol{f}_{21}}(t_{2|\boldsymbol{f}_{21}})\right\| \to 0$ and $\left\|\boldsymbol{c}_{|\boldsymbol{f}_{12}}(t_{1|\boldsymbol{f}_{12}}) - \boldsymbol{c}_{|\boldsymbol{f}_{21}}(t_{2|\boldsymbol{f}_{21}})\right\| \to 0$).

Because we know that $c_{1|\boldsymbol{f}_{12}}\left(t_{1|\boldsymbol{f}_{12}}\right) = 0$ and $c_{2|\boldsymbol{f}_{21}}\left(t_{2|\boldsymbol{f}_{21}}\right) = 0$, to have $\left\|\boldsymbol{c}_{|\boldsymbol{f}_{12}}(t_{1|\boldsymbol{f}_{12}}) - \boldsymbol{c}_{|\boldsymbol{f}_{21}}(t_{2|\boldsymbol{f}_{21}})\right\| \to 0$ requires the concentration of the resource that is the second to run out at the time the first runs out converges to zero (i.e. $c_{2|\boldsymbol{f}_{12}}\left(t_{1|\boldsymbol{f}_{12}}\right) \to 0$ and $c_{1|\boldsymbol{f}_{21}}\left(t_{2|\boldsymbol{f}_{21}}\right) \to 0$).

To have either resource be able to run out first means that while in subspace $S_{11}$ at least one species is eating each resource. Species don't stop eating a resource until it runs out, so we can be confident that after the first of $R_1$ and $R_2$ runs out at least one species is eating the remaining resources, and the concentration of that second resource is decreasing at a nonzero rate.

We've just shown that $c_{2|\boldsymbol{f}_{12}}\left(t_{1|\boldsymbol{f}_{12}}\right) \to 0$ and $c_{1|\boldsymbol{f}_{21}}\left(t_{2|\boldsymbol{f}_{21}}\right) \to 0$ as $\|\boldsymbol{f}_{12} - \boldsymbol{f}_{21}\| \to 0$ and that $\frac{d}{dt} c_{2|\boldsymbol{f}_{12}}(t) < 1$ for $t_{1|\boldsymbol{f}_{12}} < t < t_{2|\boldsymbol{f}_{12}}$ and $\frac{d}{dt} c_{1|\boldsymbol{f}_{21}}(t) < 1$ for $t_{2|\boldsymbol{f}_{21}} < t < t_{1|\boldsymbol{f}_{21}}$. Using these statements and that $t_{2|\boldsymbol{f}_{12}}$ and $t_{1|\boldsymbol{f}_{21}}$ are defined by $c_{2|\boldsymbol{f}_{12}}\left(t_{2|\boldsymbol{f}_{12}}\right) = 0$ and $c_{1|\boldsymbol{f}_{21}}\left(t_{1|\boldsymbol{f}_{21}}\right) = 0$, we can now conclude that $\left(t_{2|\boldsymbol{f}_{12}} - t_{1|\boldsymbol{f}_{12}}\right) \to 0$ and $\left(t_{1|\boldsymbol{f}_{21}} - t_{2|\boldsymbol{f}_{21}}\right) \to 0$ as $\|\boldsymbol{f}_{12} - \boldsymbol{f}_{21}\| \to 0$. (i.e. Because $\|\boldsymbol{f}_{12} - \boldsymbol{f}_{21}\| \to 0$ means that there's only an infinitesimal amount of the second resource to run out left when the first runs out and that resource concentration is strictly decreasing, the second resource to run out runs out infinitesimally soon after the first runs out when $\|\boldsymbol{f}_{12} - \boldsymbol{f}_{21}\| \to 0$.)

The population sizes and resource concentrations always vary continuously, so $\left(t_{2|\boldsymbol{f}_{12}} - t_{1|\boldsymbol{f}_{12}}\right) \to 0$ means $\left\|\boldsymbol{n}_{|\boldsymbol{f}_{12}}(t_{2|\boldsymbol{f}_{12}}) - \boldsymbol{n}_{|\boldsymbol{f}_{12}}(t_{1|\boldsymbol{f}_{12}})\right\| \to 0$ and $\left\|\boldsymbol{c}_{|\boldsymbol{f}_{12}}(t_{2|\boldsymbol{f}_{12}}) - \boldsymbol{c}_{|\boldsymbol{f}_{12}}(t_{1|\boldsymbol{f}_{12}})\right\| \to 0$ (and similar for the trajectory starting at $\boldsymbol{f}_{21}$).

We can now use the transitive property of convergence (which exists because $\boldsymbol{n} \in \mathbb{R}^{N_{\text{sp}}}$ and $\boldsymbol{c} \in \mathbb{R}^{N_{\text{re}}}$ and our distance metrics are assumed to be well-behaved choices) to conclude that $\left\|\boldsymbol{n}_{|\boldsymbol{f}_{12}}(t_{2|\boldsymbol{f}_{12}}) - \boldsymbol{n}_{|\boldsymbol{f}_{21}}(t_{1|\boldsymbol{f}_{21}})\right\| \to 0$ and $\left\|\boldsymbol{c}_{|\boldsymbol{f}_{12}}(t_{2|\boldsymbol{f}_{12}}) - \boldsymbol{c}_{|\boldsymbol{f}_{21}}(t_{1|\boldsymbol{f}_{21}})\right\| \to 0$.

The times $t_{2|\boldsymbol{f}_{12}}$ and $t_{1|\boldsymbol{f}_{21}}$ are when the second of $R_1$ and $R_2$ runs out, which is when subspace $S_{00}$ is entered. So what we have shown so far is that across a switch in the resource depletion order the state of the system when $S_{00}$ is entered varies continuously with the initial species fractions $\boldsymbol{f}$. And we know that the dynamics will continuously map the state when $S_{00}$ is entered to the final population fractions.

Therefore, **the day-to-day map is continuous even across a change in the resource-depletion order.** (More specifically, it is continuous and piecewise smooth.)

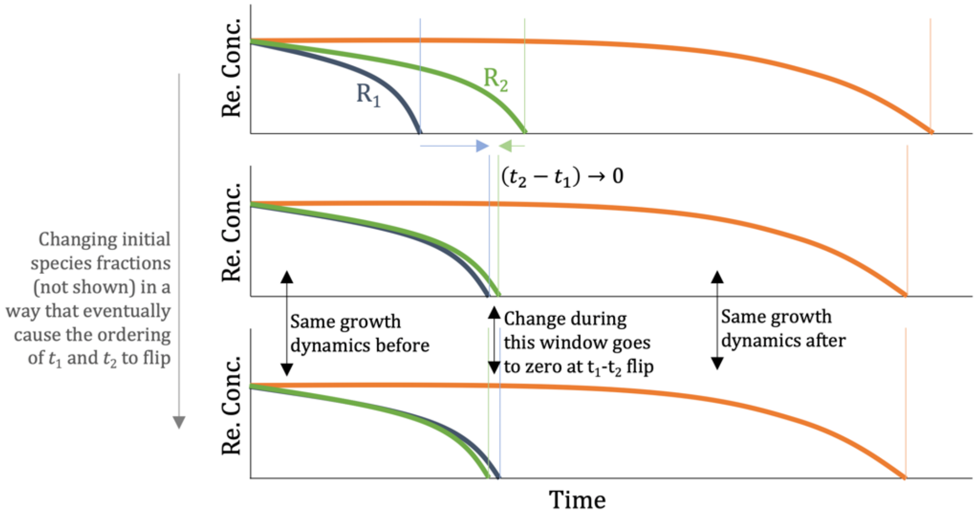


**Supp. Fig. 42.** Illustration of why the day-to-day map is continuous even when the resource depletion order changes. As the change in the ordering of two depletions, the time in between the depletion goes to zero as does the amount the system can change during that time. The dynamics after the two depletion do not depend on the order in which resources ran out.

#### *Continuity and lags*

Diauxic lag times can unfortunately break continuity. But, there are many formulations of diauxic lags that do maintain continuity. For example, continuity is maintained if lag times depend only on the resource that is being switched *to* (not the resource that is being switched *from*) and progress through a lag is reset if the resource that is being switched to changes before the species has finished its lag (e.g. if another species finishes it). These conditions maintain continuity by preventing a species from suddenly gaining or losing a significant lag when the ordering of resource depletions changes.

We can state the above more mathematically. If we are still considering a flip in the depletion order of $R_1$ and $R_2$, and $S_{00}$ is still the space that is entered, it remains that case that population sizes and resource concentrations when $S_{00}$ is entered still vary continuously with initial species fractions. But, population sizes and resource concentrations are no longer the complete state of the system. The system's state also includes how much time (if any) species have left in their diauxic lags. If species could accrue different amounts of lag based on whether $R_1$ runs out just before $R_2$ or the other way around then the state of the system when $S_{00}$ is entered would no longer vary continuously with initial species fractions. But these issues are avoided if lags only depend on the resource being switched to, because a species that would switched either from $R_1$ to $R_2$ to $R_3$ or from $R_1$ directly to $R_3$ depending on the depletion order experiences its switching-to-$R_3$ lag (if any) regardless of the depletion order.

In all our discussion of the diauxie models with lags, there are only two resources, so even if we occasionally refer to lags by the resource that has run out, which resource runs out defines which resource is being switched to, so the lags do only depend on which resource is being switched to. Therefore, **in all our discussion of diauxie models with lags in this Supplemental Information the models are indeed continuous.**

#### *Continuity and co-utilization*

This will not be discussed in depth here, but it is noted that co-utilization state have a surprising potential to break continuity across changes to the resource depletion order. For example, if the only species consuming $R_2$ was co-utilizing $R_1$ and $R_2$ but switches to a co-utilization state that doesn't involve $R_2$ after $R_1$ runs out then the part of the continuity argument that $\left|t_{2|\boldsymbol{f}_{12}} - t_{1|\boldsymbol{f}_{21}}\right| \to 0$ breaks down. This is, however, resolved if species never drop a resource from the set of resources they are co-utilizing as a result of another resource being depleted. But, if there are also lags continuity breaks again because if the only species eating $R_2$ has a lag after $R_1$ runs out then $\left|t_{2|\boldsymbol{f}_{12}} - t_{1|\boldsymbol{f}_{21}}\right| \to 0$ breaks down. This breakdown in continuity is solved if growth rate recoveries are smooth rather than sharp (i.e. if growth rate are zero only instantaneously). The use of smooth growth rate recoveries in the main text (as well as continuity being more readily maintained with only two species) allows Pa's co-utilization followed by lag to not break continuity.

#### Combination of diauxic lags and species having different growth rates for each resource

Combining diauxic lag with species having different growth rates for each resource creates a large set of permutations to survey. In many cases, the phenomena that occur are slight changes to what could already happen in the simpler version of the diauxie model that have already been explored. In the interest of brevity, we will not survey all possible outcomes in this version of the model, but will instead highlight just two points of interest:

First, in the case of species having the same resource preference a bistability can be created when the initial fast-grower is the slow-switcher and also the fast-grower on the second resource and is so by an even larger factor than it was on the first (i.e. $\frac{g_{\alpha 2}}{g_{\beta 2}} > \frac{g_{\alpha 1}}{g_{\beta 1}}$). Because the fast-grower has a lag, the slow-grower still has a chance at survival. The positive feedback loop that creates the bistability is as follows: (i) As the population fraction of α is increased and the fraction of β decreased, $R_1$ runs out earlier (because α eats it more quickly). The amount of $R_2$ that β eats during α's lag also deceases, leaving more available for α. Thus, increasing α's population fraction increases the time the population spends growing on $R_2$ relative to the time it spends on $R_1$. Because α increases its population fraction relative to β faster when growing on $R_2$ (assuming α is able to finish its lag), more time on $R_2$ benefits α, further increasing its population fraction and further increasing the amount it gets to grow on $R_2$. This creates a positive feedback and ultimately a bistability. The math behind this bistability is easy enough to work out if one follows the methodology used elsewhere in the Supplemental Information (but remember that α may or may not get to eat $R_2$ at fixed points and when invading β).

Second, it is possible to have a tri-stability between each species excluding the other and a coexistence state. This is explored in more detail below:

##### *A possible tri-stability when species have opposite resource preferences*

In the case of species having opposite resource preferences, there are five qualitatively distinct fixed points to consider:

| | | | |
|---|---|---|---|
| i. | $R_1$ runs out first. | α has no lag. | |
| ii. | $R_1$ runs out first. | α has lag. | α finishes lag in time to eat $R_2$ |
| iii. | $R_1$ runs out first. | α has lag. | α does not finish in time to eat $R_2$ |
| iv. | $R_2$ runs out first. | β has no lag. | |
| v. | $R_2$ runs out first. | β has lag. | β finishes lag in time to eat $R_1$ |
| vi. | $R_2$ runs out first. | β has lag. | β does not finish in time to eat $R_1$ |

With fixed points (i) and (iv), the species that finishes its preferred resource first does not have a lag time. It does not matter if the other species has a lag time because that species just continues eating it preferred resource. So these will have the same mathematical solutions and behavior as in the case of simple diauxie without lags.

Fixed point (v) won't have any qualitatively new behavior that fixed point (iv) did not. This is because it's still the case that β is falling behind α until it starts eating $R_1$, at which point if β is the $R_1$ fast-grower it has a chance to catch up to α. This will still be an unstable fixed point because it will still be the case that a positive feedback exists whereby an increase in β's population fraction causes $R_2$ to run out sooner and β to start growing on $R_1$ sooner.

Fixed point (vi) cannot exist because β is the initial slow-grower and then does not finish its lag, so it cannot possibly keep up with α.

Fixed points (ii) and (iii) will have the most interesting behavior. If $g_{\alpha 2} > g_{\alpha 1}$ it is possible to get both fixed points. If we also have $g_{\beta 1} > g_{\alpha 1} > g_{\beta 2}$ it is possible to get the bistability that occurred in the simpler diauxie model without lag times.

We therefore consider the case of α eating $R_1$ then $R_2$, β eating $R_2$ then $R_1$, α having a lag when $R_1$ runs out, α preferring the resource it grows slowest on ($g_{\alpha 2} > g_{\alpha 1}$), and β also preferring the resource it grower slowest on and not being the initial fast-grower because of that preference ($g_{\beta 1} > g_{\alpha 1} > g_{\beta 2}$).

As we solve for each of three possible fixed points, the numbering will maintain consistency with the above discussion. We will be solving for fixed points (ii), (iii), and (iv).

*Fixed point (ii)* has growth-by-DF requirements:

$$g_{\alpha 1} t_1 + g_{\alpha 2}\left(t_2 - t_1 - t_{\text{lag},\alpha}\right) = \log(\text{DF})$$
$$g_{\beta 2}\, t_2 = \log(\text{DF})$$

This is solved by:

$$t_1 = \frac{g_{\beta 2} - g_{\alpha 2}}{(g_{\alpha 1} - g_{\alpha 2}) g_{\beta 2}} \log(\text{DF}) + \frac{g_{\alpha 2}}{g_{\alpha 1} - g_{\alpha 2}} t_{\text{lag},\alpha} \quad \text{and} \quad t_2 = \frac{1}{g_{\beta 2}} \log(\text{DF})$$

Species fractions are:

$$f_\alpha^{\text{(FP ii)}} = \frac{s_1(\text{DF} - 1)}{\exp\left(\frac{g_{\alpha 1}(g_{\beta 2} - g_{\alpha 2})}{g_{\beta 2}(g_{\alpha 1} - g_{\alpha 2})} \log(\text{DF}) + \frac{g_{\alpha 1} g_{\alpha 2}}{g_{\alpha 1} - g_{\alpha 2}} t_{\text{lag},\alpha}\right) - 1}$$

And the fixed point exists if $t_1 > 0$, $t_2 - t_1 < t_{\text{lag},\alpha}$, and $0 < f_\alpha < 1$ using the above expressions. The condition that $t_2 - t_1 < t_{\text{lag},\alpha}$ simplifies to:

$$\left\{g_{\alpha 2} < g_{\alpha 1} \text{ and } \left(\frac{1}{g_{\beta 2}} - \frac{1}{g_{\alpha 1}}\right) \log(\text{DF}) > t_{\text{lag},\alpha}\right\} \quad \text{or} \quad \left\{g_{\alpha 2} > g_{\alpha 1} \text{ and } \left(\frac{1}{g_{\beta 2}} - \frac{1}{g_{\alpha 1}}\right) \log(\text{DF}) < t_{\text{lag},\alpha}\right\}$$

In the case of $g_{\alpha 2} > g_{\alpha 1}$ this point will be unstable (because increases α's species fraction will cause it to deplete $R_1$ sooner and then also finish its lag and start growing on $R_2$ at its faster growth rate sooner).

*Fixed point (iii)* has growth-by-DF requirements:

$$g_{\alpha 1} t_1 = \log(DF) \quad \text{and} \quad g_{\beta 2}\, t_2 = \log(\text{DF})$$

The fractions for this fixed point are simply $f_\alpha^{\text{(FP iii)}} = s_1$ and $f_\beta^{\text{(FP iii)}} = s_2$ and the existence requirement is that β finishes $R_2$ before α finishes its lag:

$$\left(\frac{1}{g_{\beta 2}} - \frac{1}{g_{\alpha 1}}\right) \log(\text{DF}) < t_{\text{lag},\alpha}$$

Notice how if $g_{\alpha 2} > g_{\alpha 1}$ both fixed point (*ii*) and fixed point (*iii*) are possible.

*Fixed point (iv)* is the same as was calculated in the case of no lags:

$$f_\alpha{}^{(\text{FP iv})} = 1 - \frac{(1 - s_1)(\text{DF} - 1)}{DF^{\left(\frac{g_{\beta 2}(g_{\beta 1} - g_{\alpha 1})}{g_{\alpha 1}(g_{\beta 1} - g_{\beta 2})}\right)} - 1}$$

Plotting all three fixed points on a bifurcation diagram as the resource supply is varied:

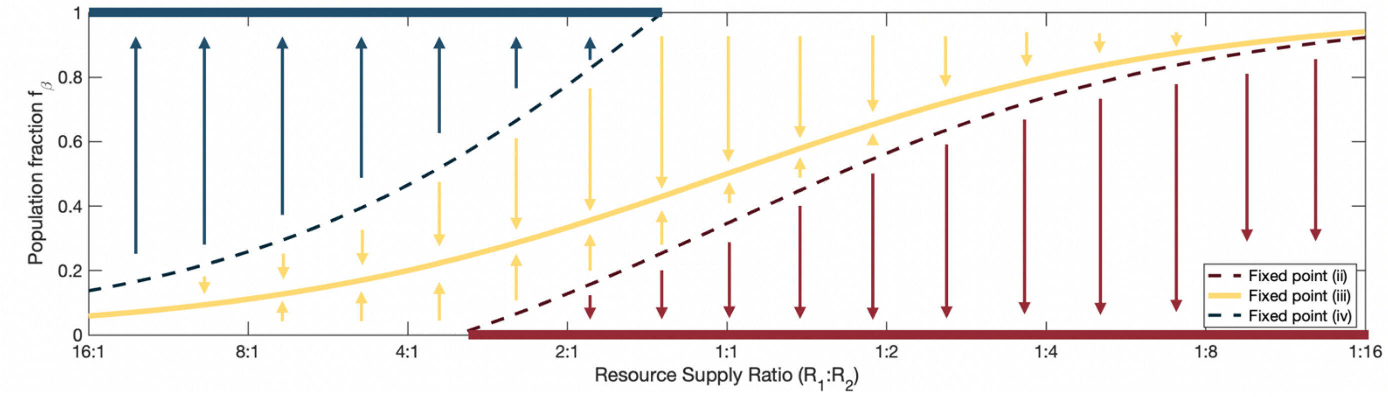


**Supp. Fig. 43.** Bifurcation diagram for a diauxie-driven tri-stability. Species α eats $R_1$ then $R_2$. Species β has the opposite resource preference, eating $R_2$ then $R_1$. Growth rates are $g_{\alpha 1} = 0.4$, $g_{\alpha 2} = 0.9$, $g_{\beta 1} = 1.5$, and $g_{\beta 2} = 0.3$. These growth rates are such that both species are anomalous (i.e. $g_{\alpha 2} > g_{\alpha 1}$ even though α eats $R_1$ first and $g_{\beta 1} > g_{\beta 2}$ even though β eats $R_2$ first). Species α has lag time $t_{\text{lag},\alpha} = 2.25$ when $R_1$ runs out. The dilution factor is 10.

These three outcomes occur by the following mechanisms:

- Coexistence occurs with $R_1$ running out first and α not finishing its lag such that each species only ever grows on its preferred resource.
- If $R_2$ runs out first (which happens when the supply fraction of $R_2$ is lowered and the population fraction of β is increased), β has a period of growing on $R_1$, during which time is grows faster than α, creating a possibility of β excluding α once its population is large enough.
- If α finishes its lag before $R_1$ runs out (which happens when the supply fraction of $R_1$ is lowered and the population fraction of β is decreased), it has a period of growing on $R_2$ when it grows faster than β by an even larger factor, creating the possibility of α excluding β once its population fraction is large enough.

It is unlikely this scenario could be realized experimentally (given its large set of requirements, including two species having anomalous resource preferences for the resource they grow slowest, both growing purely diauxically, and there being no other significant interactions even at the relatively low dilution factor), but it is an interesting modelling curiosity.

### Diauxic lags and cross-feeding

Modeling the interactions between diauxic lags and cross-feeding could be several papers of interesting material. Here, we very briefly look at two examples of simple scenarios in which diauxic lags and cross-feeding interact. The first demonstrates how little the analysis can need to change to incorporate cross-feeding into a diauxic lag model. The second suggests how diauxic lags could create a pressure towards specializing in specific steps in sequential degradations and other cross-feeding networks.

#### *Fast-growing, slow-switching primary degrader vs slower-growing cross-feeder*

In this example, we consider a primary degrader (α) consuming resource $R_1$ and releasing $R_2$ as a byproduct and a cross-feeder (β) consuming resource $R_2$. When $R_1$ runs out, α switches to $R_2$. After $R_2$ is depleted and the population has saturated, the population size is diluted and fresh resources are supplied.

Both species have a single growth rate ($g_\alpha$ and $g_\beta$ with $g_\alpha > g_\beta$). If there are no lags, α is always growing faster than β and always increasing its population fraction. Whenever both species are growing at their full growth rates (i.e. when β is not limited by waiting for α to release more $R_2$ – in which case α is only going to perform even better in comparison to β):

$$n'_\alpha(t) = g_\alpha n_\alpha(t) = g_\alpha\, f_\alpha(t)\, n_{\text{tot}}(t) \quad \text{and} \quad n'_\beta(t) = g_\beta f_\beta(t)\, n_{\text{tot}}(t)$$

$$f'_\alpha(t) = \frac{d}{dt}\left(\frac{n_\alpha(t)}{n_\alpha(t) + n_\beta(t)}\right) = \frac{n_\beta(t)\, n'_\alpha(t) - n_\alpha(t)\, n'_\beta(t)}{\left(n_\alpha(t) + n_\beta(t)\right)^2}$$

$$f'_\alpha(t) = \frac{\left(f_\beta(t)\, n_{\text{tot}}(t)\right)\left(g_\alpha\, f_\alpha(t)\, n_{\text{tot}}(t)\right) - \left(f_\alpha(t)\, n_{\text{tot}}(t)\right)\left(g_\beta\, f_\beta(t)\, n_{\text{tot}}(t)\right)}{n_{\text{tot}}(t)^2}$$

$$f'_\alpha(t) = \left(g_\alpha - g_\beta\right) f_\alpha(t)\, f_\beta(t)$$

(This equation would also apply to many cases within the "pure" diauxie modeling and holds any time two species are growing at constant exponential growth rates.)

Because α it is always increasing its species fraction, **in order to have β survive it is necessary to implement some feature that will give β a period in which it can catch back up to α. This could be various model additions, including α having a diauxic lag**, which is the addition we will be looking at in this subsection.

Before proceeding, we need to define how much $R_2$ α releases as it's growing on $R_1$. We will give α a ½ yield, defined by: For every unit of $R_1$ that α consumes, it gains half a unit of population size and releases half a unit of $R_2$. For every unit of $R_2$ consumed by either species, one unit of population size is gained.

**Consideration of whether β is limited by α's $R_2$ release.** As α is growing on $R_1$ and β is growing on the $R_2$ released by α, the flux-balance for the concentration $c_2(t)$ of $R_2$ is

$$c'_2(t) = g_\alpha n_\alpha(t) - n'_\beta(t)\,.$$

If β is not limited by the $R_2$ supply $n'_\beta(t) = g_\beta n_\beta(t)$. But, we cannot have $c_2(t) < 0$. So if $c_2(t) = 0$ (i.e. there is no remaining reservoir of $R_2$) and $g_\beta n_\beta(t) > g_\alpha n_\alpha(t)$ (i.e. β would be consuming $R_1$ faster than α produces it) then $n'_\beta(t) = g_\alpha n_\alpha(t) = {n_\alpha}'(t)$ (i.e. β grows at a rate determined by eating $R_2$ as quickly as it is produced).

The condition for whether β is limited by the $R_2$ supply rearranges to become a function of the species fractions:

$$\beta \text{ is limited by } R_2 \text{ supply} \quad \text{iff} \quad g_\beta n_\beta(t) > g_\alpha n_\alpha(t) \ \rightarrow \ f_\beta(t) > \frac{g_\alpha}{g_\alpha + g_\beta}$$

Because $f_\beta(t)$ will only ever be decreasing when α is growing at its full initial growth rate, once β is no longer limited by the α's release of $R_2$ β will never become limited by $R_2$ again.

If we start at $n_\alpha(0) = \frac{1-f_{\beta 0}}{\mathrm{DF}-1}$ and $n_\beta(0) = \frac{f_{\beta 0}}{\mathrm{DF}-1}$ and assume $f_{\beta 0}$ is such that β is initially limited by α's $R_2$ release then while β is still limited by $R_2$ supply:

$$n_\alpha(t) = \frac{1-f_{\beta 0}}{\mathrm{DF}-1}\exp(g_\alpha t) \quad \text{and} \quad n_\beta(t) = n_\beta(0) + \Delta n_\alpha = \frac{f_{\beta 0}}{\mathrm{DF}-1} + \frac{1-f_{\beta 0}}{\mathrm{DF}-1}(\exp(g_\alpha t) - 1).$$

Species β will stop being limited by $R_2$ supply when

$$g_\beta\left(\frac{f_{\beta 0}}{\mathrm{DF}-1} + \frac{1-f_{\beta 0}}{\mathrm{DF}-1}(\exp(g_\alpha t_{\mathrm{unlimd}}) - 1)\right) = g_\alpha\left(\frac{1-f_{\beta 0}}{\mathrm{DF}-1}\exp(g_\alpha t_{\mathrm{unlimd}})\right).$$

Solving this equation for the time $t_{\mathrm{unlimd}}$ at which β stops being limited by $R_2$ supply produces

$$t_{\mathrm{unlimd}} = \frac{1}{g_\alpha}\log\left(\frac{g_\beta}{(1-f_{\beta 0})\, g_\alpha}\right).$$

At which point population sizes will be

$$n_\alpha(t_{\mathrm{unlimd}}) = \frac{g_\beta}{(\mathrm{DF}-1)g_\alpha} \quad \text{and} \quad n_\beta(t_{\mathrm{unlimd}}) = \frac{(2f_{\beta 0}-1)g_\alpha + g_\beta}{(\mathrm{DF}-1)g_\alpha}.$$

We can calculate the condition for whether β stops being limited by $R_2$ before or after $R_1$ runs out by first calculating the time $t_1$ at which $R_1$ runs out and then setting up the inequality $t_1 < t_{\mathrm{unlimd}}$:

$$n_\alpha(t_1) - n_\alpha(0) = \frac{1-f_{\beta 0}}{\mathrm{DF}-1}(\exp(g_\alpha t_1) - 1) = \frac{1}{2} \quad \rightarrow \quad t_1 = \frac{1}{g_\alpha}\log\left(\frac{(\mathrm{DF}+1) - 2f_{\beta 0}}{2(1-f_{\beta 0})}\right)$$

$$\frac{1}{g_\alpha}\log\left(\frac{(\mathrm{DF}+1) - 2f_{\beta 0}}{2(1-f_{\beta 0})}\right) < \frac{1}{g_\alpha}\log\left(\frac{g_\beta}{(1-f_{\beta 0})\, g_\alpha}\right)$$

$$\frac{(\mathrm{DF}+1) - 2f_{\beta 0}}{2(1-f_{\beta 0})} < \frac{g_\beta}{(1-f_{\beta 0})\, g_\alpha}$$

$$\beta\text{'s growth rate will be limited by } R_2 \text{ throughout the cycle} \quad \text{iff} \quad f_{\beta 0} > \frac{g_\beta}{g_\alpha} + \frac{1}{2}(\mathrm{DF}+1).$$

However, $\mathrm{DF} > 1$, so $\frac{1}{2}(\mathrm{DF}+1) > 1$ and above reduces to requiring $f_{\beta 0} > 1$, so there is no initial population fraction $f_{\beta 0}$ (either corresponding to a fixed point or not) that will result in β being limited by α's release of $R_1$ for the entirety of the time until $R_1$ is depleted.

Returning to the condition that

$$\beta \text{ is limited by } R_2 \text{ supply} \quad \text{iff} \quad f_\beta(t) > \frac{g_\alpha}{g_\alpha + g_\beta},$$

we note that β being initially limited by $R_2$ supply requires $f_{\beta 0} > \frac{g_\alpha}{g_\alpha + g_\beta}$. When $g_\alpha > g_\beta$, the fraction $\frac{g_\alpha}{g_\alpha + g_\beta} > \frac{1}{2}$, so $f_{\beta 0} > 0$ must be true for β to be initially limited by $R_2$ supply. But, β only ever eats $R_2$ while α gets to eat all of $R_1$ and maybe even some of $R_2$, which means at a fixed point $f_\beta \le \frac{1}{2}$. This is a contradiction and means that **at a fixed point β's growth rate is never limited by α's release of $R_2$.**

**Coexistence between α and β.** We now turn to considering the possibility of coexistence between α and β. Similar to what we did in analyzing the diauxie model without cross-feeding, we start by looking for whether there can be fixed points. Because β's growth rate is never limited by α's release of $R_2$ at a fixed point, it turns out that the fact $R_2$ is a cross-fed resource and not a supplied resource makes no difference to the math.

α grows at rate $g_\alpha$ until the supply of $R_1$ is depleted (with an effective supply of $s_1 = ½$ being implemented to capture α's ½ yield) and then experiences its lag and switches to $R_2$. β grows at rate $g_\beta$ on $R_2$. The population saturates when the effective supply $s_2 = ½$ of $R_2$ is depleted. Way back near the start of this Supplemental Information, we saw that if α finishes its lag, the time $t_2$ at which $R_2$ runs out is over-determined by the species' growth-by-DF requirements, so the only fixed points are when α does not finish its lag.

The fixed point with α not finishing its lag and species have opposite resource preferences has population fractions $f_a = s_1 = ½$ and $f_\beta = s_2 = ½$. This species fraction agree with β never being limited by $R_2$. That this point exists with α never finishing its lag requires

$$t_{\text{lag},\alpha} > \left(\frac{1}{g_\beta} - \frac{1}{g_\alpha}\right) \log(DF) \,.$$

(That's it. This is the fixed point of the system. It really doesn't look any different than in the diauxie model without cross-feeding. And this fixed point requires $t_{\text{lag},\alpha} > 0$ for its existence.)

**If α's yield and $R_1$ cross-feeding release are not equal to ½.** To try to make things more interesting, let's now say that for every unit of $R_1$ that α consumes, it gains $y_{\alpha 1}$ units of population size and releases $(1 - y_{\alpha 1})$ units of $R_2$. So, for each unit of population size it gains from $R_1$, α releases $\frac{(1-y_{\alpha 1})}{y_{\alpha 1}}$ units of $R_2$. Both species gain one unit of population size for each unit of $R_2$ they consume.

Carrying this through some of the calculations:

$$\beta \text{ is limited by } \mathrm{R}_2 \text{ supply} \quad \text{iff} \quad g_\beta n_\beta(t) > \frac{(1 - y_{\alpha 1})}{y_{\alpha 1}} g_\alpha n_\alpha(t) \;\; \rightarrow \;\; f_\beta(t) > \frac{g_\alpha(1 - y_{\alpha 1})}{g_\alpha(1 - y_{\alpha 1}) + g_\beta y_{\alpha 1}}$$

With $g_\alpha > g_\beta$, the RHS cannot become smaller than $(1 - y_{\alpha 1})$, so for β to be initially limited by $R_2$ supply requires at least $f_{\beta 0} > (1 - y_{\alpha 1})$. But, at a fixed point, because α gets all of $R_1$ to itself $f_{\alpha 0} \geq y_{\alpha 1}$ and $f_{\beta 0} \leq 1 - y_{\alpha 1}$. This creates the same contradiction and means that it is still the case that at any fixed point β is never limited by α's $R_2$ release, and the only possible fixed point occurs at $f_\alpha = y_{\alpha 1}$ and $f_\beta = 1 - y_{\alpha 1}$ and requires

$$t_{\text{lag},\alpha} > \left(\frac{1}{g_\beta} - \frac{1}{g_\alpha}\right) \log(DF) \,.$$

#### *Fast-grower vs fast-switcher during sequential degradation*

In this example, we consider a simplified model of a three-step degradation of a complex resource. There will be two species in this example: the fast-grower α and the fast-switcher β.

Species grow first on $R_1$ while producing $R_2$ as a byproduct, then on $R_2$ while producing $R_3$, and finally on $R_3$ with no further byproduct production. These resources could be thought of as a polysaccharide, leftover monomers, and a generalized pool of amino acids and other metabolic byproducts. Resources become progressively slower to grow on such that $g_{\alpha 1} > g_{\alpha 2} > g_{\alpha 3}$ and $g_{\beta 1} > g_{\beta 2} > g_{\beta 3}$. In this example we will use:

$$g_{\alpha 1} = 1.0 \quad | \quad g_{\alpha 2} = 0.9 \quad | \quad g_{\alpha 3} = 0.3$$
$$g_{\beta 1} = 0.8 \quad | \quad g_{\beta 2} = 0.7 \quad | \quad g_{\beta 3} = 0.1$$

If both species excrete the same fraction of $R_1$ as $R_2$ and both excrete the same fraction of $R_2$ as $R_3$, then there is an effective resource supply $\boldsymbol{s} = (s_1 \quad s_2 \quad s_3)$ where $s_i$ captures the total (linear-scale) growth of the population on each resource. For this example we will use $\boldsymbol{s} = (0.2 \quad 0.2 \quad 0.6)$.

When a resource is depleted, β is able to switch to the next resource without any lag, while α has a lag lasting $t_{\text{lag},\alpha} = 2$. If α has not yet finished its first lag when $R_2$ runs out, its progression through its lag is lost and it does not grow until $t_{\text{lag},\alpha}$ has passed since the $R_2$ depletion.

In this example, we start the populations at equal species fractions and a total population size $10^{-2}$ times carrying capacity:

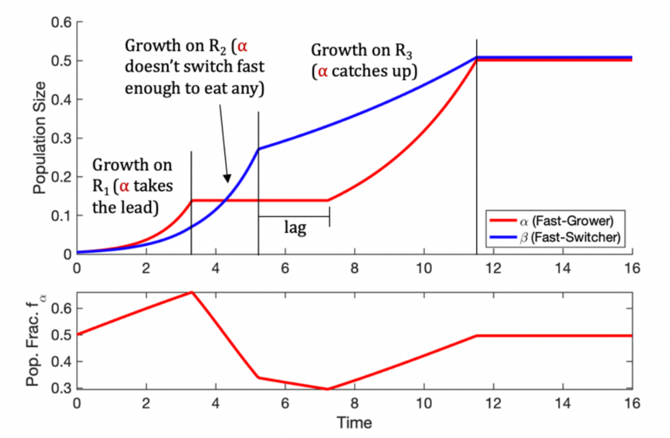


**Supp. Fig. 41.** In a three-step degradation of a complex resource, the fast-grower/slow-switcher initially becomes the dominant species (largest population fraction) then declines in population fraction as it fails to switch in time to eat intermediate cross-feeding products. When it finally switches in time to eat a resource it reappears as a more significant fraction of the community.

In this example, α appears as the dominant species early on then declines in population fraction as a result of its diauxic lags – not even getting to grow at all on $R_2$ – but later recovers once it switches to $R_3$ in time to grow on that resource before it runs out.

Thus, diauxic lags prevent α from consuming $R_2$ and give its population fraction a rise-fall-rise pattern. It is unclear how realistic this scenario is, but this illustrates how diauxic lags would interfere with a species trying to participate in every step in a degradation and create a pressure towards specializing in only a single step in such a cross-feeding network.

**Supp. Fig. 42 and 43.** These figures are spread across the remainder of the supplemental information. This data underlies the conclusions regarding the survey of additional species and resources (Main Text Figure 6).

Supp. Fig. 42 (*A*-*J*) contains competition data for 10 pairs of species in 6 single-resource environments and 15 corresponding two-resource environments. Data shows species fractions over time, colored by assigned qualitative outcome. (Colors corresponding to species are as colored in the titles; yellow indicates coexistence.) All competition were originally run with two replicates. Additional replicates were run for select competitions that needed additional data to clarify the outcome (usually clarifying being extinction and coexistence with a small fraction for one of the species).

Supp. Fig. 43 (*A*-*E*) contains monoculture data for each of the 5 species in each of the 15 two-resource environments. For most cases, three replicates were run. Black lines show fits of exponential growth, lag, then exponential growth. Cases with two exponential growth fits not connected by a horizontal line have no measurable lag time. When a species either (i) did not grow on both resources, (ii) did not grow fast enough in the two-resource environment for its lag to be observed before the end of the experiment, or (iii) had extra diauxic shifts*, growth rates and lag times were not fit. In the analysis presented in the main text cases in which both species had lag times less than half an hour were considered cases of neither species having a significant lag and the slow-grower not being the fast-switcher by default.

*For *Pseudomonas aurantiaca* and *Pseudomonas putida* growing on the combination of glucose and fructose, extra diauxic shifts were observed. One possibility for this would be initial fermentation followed by growth on a fermentation end-product such as acetate. Regardless of the explanation, it was impossible to fit a single lag time in these cases.

| Single-Resource Outcomes | Two-Resource Outcome | Observations | **Aci2** vs **Pa** (Supp. Fig. 42*C*) | **Aci2** vs **Pp** (Supp. Fig. 42*D*) | **Aci2** vs **Ka** (Supp. Fig. 42*E*) | **Aci2** vs **Arth** (Supp. Fig. 42*F*) | **Pa** vs **Pp** (Supp. Fig. 42*G*) | **Pa** vs **Ka** (Supp. Fig. 42*H*) | **Pa** vs **Arth** (Supp. Fig. 42*I*) | **Pp** vs **Ka** (Supp. Fig. 42*J*) | **Pp** vs **Arth** (Supp. Fig. 42*K*) | **Ka** vs **Arth** (Supp. Fig. 42*L*) |
|---|---|---|---|---|---|---|---|---|---|---|---|---|
| **A wins + A wins** | **A wins** | **49 (94%)** | 4 | | 6 | 6 | 6 | 6 | 6 | 6 | 6 | 3 |
| | B wins | | | | | | | | | | | |
| | Coex. | 3 (6%) | 2 | | | 1 | | | | | | |
| | Extinc. | | | | | | | | | | | |
| **A wins + B wins** | A wins | 8 (28%) | | | | | 4 | 2 | 2 | | | |
| | **Coex.** | **21 (72%)** | | 1 | | 8 | | 2 | 2 | 4 | 4 | |
| | Extinc. | | | | | | | | | | | |
| **A wins + Coex.** | A wins | 9 (26%) | | 1 | | | | 2 | 2 | | 4 | |
| | B wins | | | | | | | | | | | |
| | **Coex.** | **25 (74%)** | | 5 | 8 | | | 3 | 3 | 5 | 1 | |
| | Extinc. | | | | | | | | | | | |
| **Coex. + Coex.** | A wins | | | | | | | | | | | |
| | **Coex.** | **4 (100%)** | | 3 | 1 | | | | | | | |
| | Extinc. | | | | | | | | | | | |
| **Extinc. + Extinc.** | A wins | 2 (50%) | 1 | | | | | | | | | 1 |
| | Coex. | 1 (25%) | | | | | | | | | | 1 |
| | **Extinc.** | **1 (25%)** | | | | | | | | | | 1 |
| **Extinc. + A wins** | **A wins** | **18 (75%)** | 3 | 2 | | | 4 | | | | | 9 |
| | B wins | 1 (4%) | | | | | 1 | | | | | |
| | Coex. | 5 (21%) | 5 | | | | | | | | | |
| | Extinc. | | | | | | | | | | | |
| **Extinc. + Coex.** | A wins | | | | | | | | | | | |
| | **Coex** | **3 (100%)** | | 3 | | | | | | | | |
| | Ext | | | | | | | | | | | |

**Supp. Fig. 42** (*A*). Summary of the relationships between single-resource and two-resource outcomes. Highlighted are the predictions from a linear sum rule, which is 81% +/- 3% accurate in this data set (standard error of the binomial distribution reported).

**Frequency of single-resource coexistence was 17%** (9/53 cases which were not mutual extinctions).
**Frequency of two-resource coexistence was 42%** (62/149 cases which were not mutual extinctions).
Coexistence was 2.4x more likely in a two-resource environment than in a single-resource environment.

| Single-Resource Outcomes | Two-Resource Outcome | Observations | **Fructose** and **Glucose** | **Fructose** and **Citrate** | **Fructose** and **Alanine** | **Fructose** and **Glutamate** | **Fructose** and **Aspartate** | **Glucose** and **Citrate** | **Glucose** and **Alanine** | **Glucose** and **Glutamate** | **Glucose** and **Aspartate** | **Citrate** and **Alanine** | **Citrate** and **Glutamate** | **Citrate** and **Aspartate** | **Alanine** and **Glutamate** | **Alanine** and **Aspartate** | **Glutamate** and **Aspartate** |
|---|---|---|---|---|---|---|---|---|---|---|---|---|---|---|---|---|---|
| **A wins + A wins** | **A wins** | **49 (94%)** | 1 | 1 | | | | 1 | | | | 7 | 8 | 8 | 7 | 8 | 8 |
| | B wins | | | | | | | | | | | | | | | | |
| | Coex. | 3 (6%) | 1 | | | | | | | | | 1 | | | 1 | | |
| | Extinc. | | | | | | | | | | | | | | | | |
| **A wins + B wins** | A wins | 8 (28%) | | 2 | | 2 | | 1 | 1 | 1 | 1 | | | | | | |
| | **Coex.** | **21 (72%)** | | 3 | 5 | 3 | 5 | 1 | 2 | 1 | 1 | | | | | | |
| | Extinc. | | | | | | | | | | | | | | | | |
| **A wins + Coex.** | A wins | 9 (26%) | | | | | | 3 | 1 | 4 | 1 | | | | | | |
| | B wins | | | | | | | | | | | | | | | | |
| | **Coex.** | **25 (74%)** | 4 | 1 | 1 | 1 | 1 | 3 | 4 | 2 | 5 | 1 | | | 1 | 1 | |
| | Extinc. | | | | | | | | | | | | | | | | |
| **Coex. + Coex.** | A wins | | | | | | | | | | | | | | | | |
| | **Coex.** | **4 (100%)** | 1 | | | | | | | | | | 1 | 1 | | | 1 |
| | Extinc. | | | | | | | | | | | | | | | | |
| **Extinc. + Extinc.** | A wins | 2 (50%) | 1 | | | | | | | | | | | | 1 | | |
| | Coex. | 1 (25%) | | | | | | | | | | | | | | 1 | |
| | **Extinc.** | **1 (25%)** | | | | | | | | | | | | | | | 1 |
| **Extinc. + A wins** | **A wins** | **18 (75%)** | 2 | 2 | 3 | 2 | 3 | | 1 | 1 | 1 | 1 | 1 | 1 | | | |
| | B wins | 1 (4%) | | | 1 | | | | | | | | | | | | |
| | Coex. | 5 (21%) | | | | 1 | | 1 | 1 | 1 | 1 | | | | | | |
| | Extinc. | | | | | | | | | | | | | | | | |
| **Extinc. + Coex.** | A wins | | | | | | | | | | | | | | | | |
| | **Coex** | **3 (100%)** | | 1 | | 1 | 1 | | | | | | | | | | |
| | Ext | | | | | | | | | | | | | | | | |

**Supp. Fig. 42** (*B*). Continued summary of the relationships between single-resource and two-resource outcomes. Highlighted are the predictions from a linear sum rule.

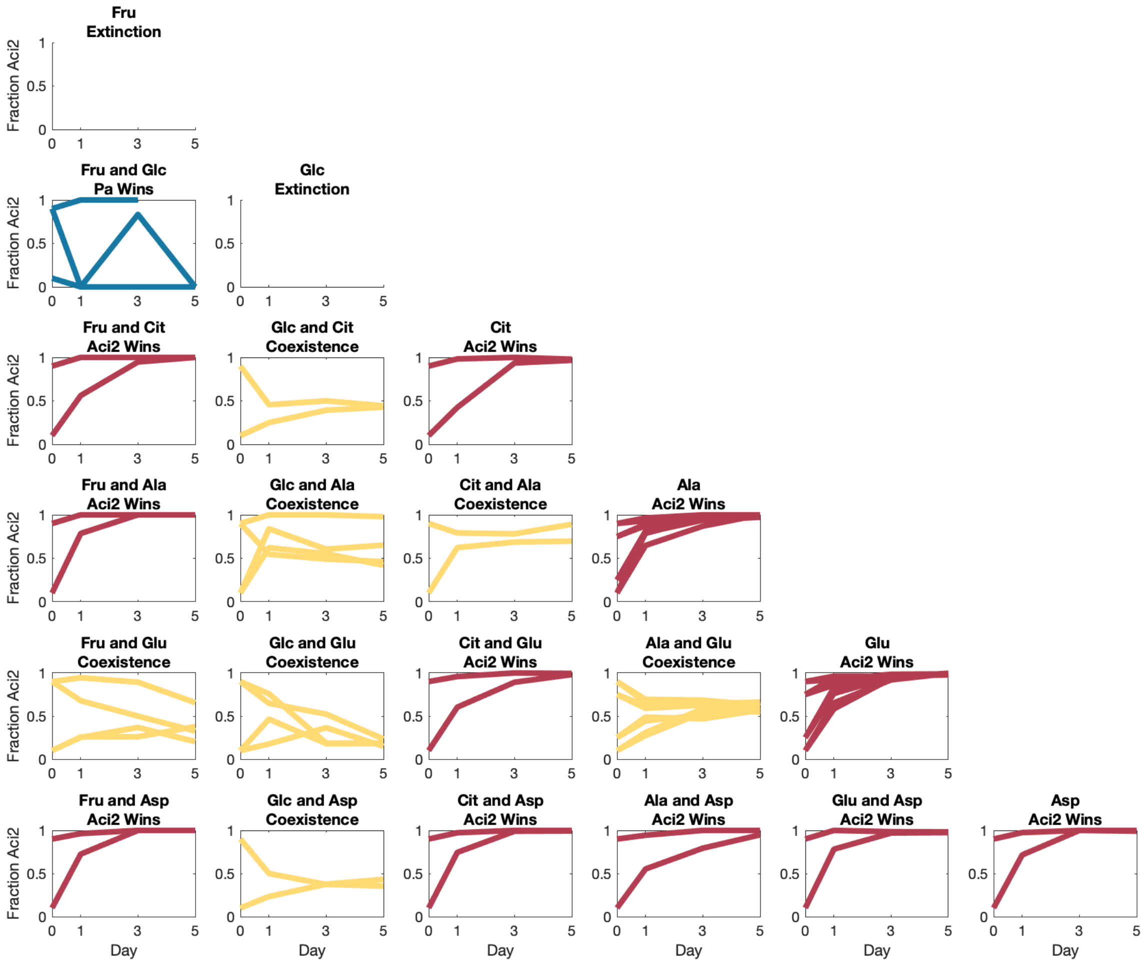


**Supp. Fig. 42** (*C*). **Aci2** vs **Pa** competition data.

On Fructose and Glucose, the two species were very close to extinction.

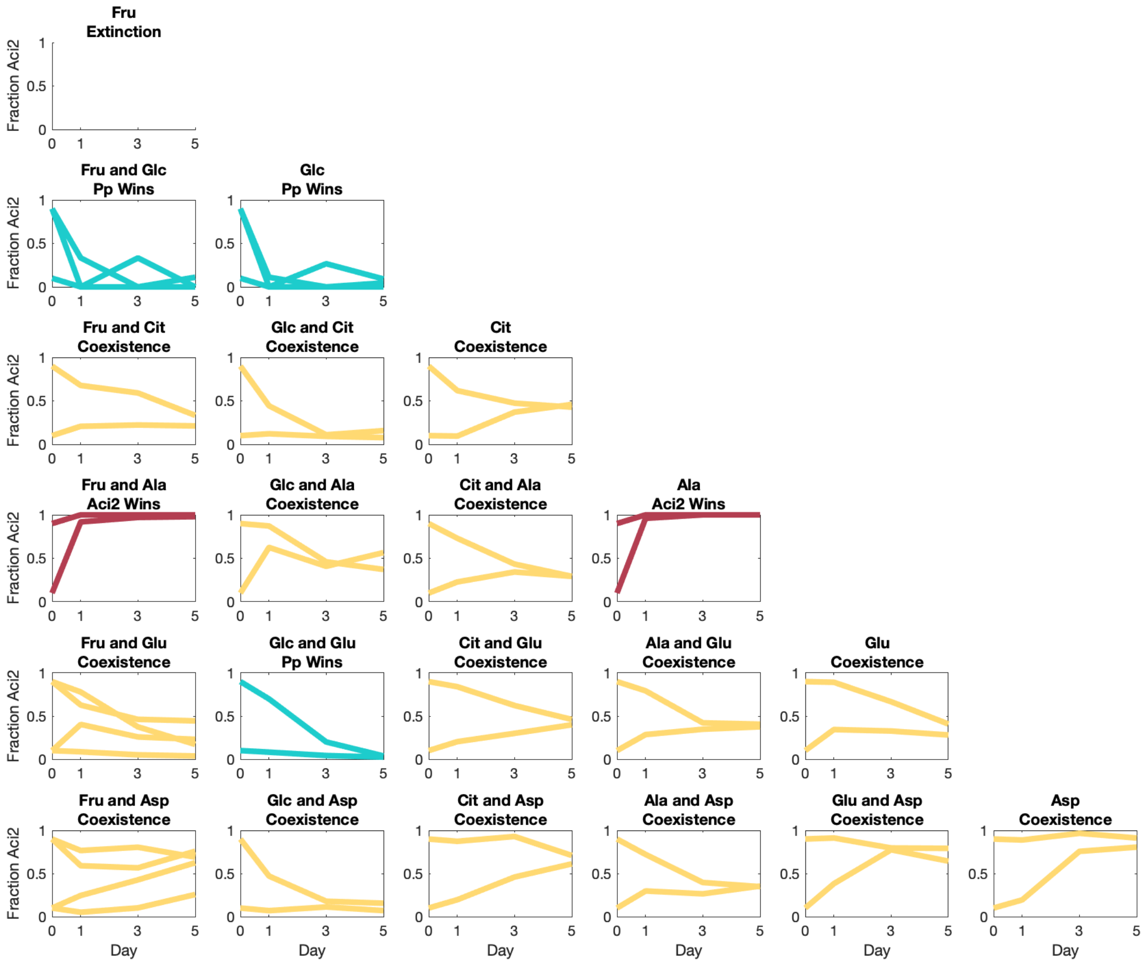


**Supp. Fig. 42** (*D*). **Aci2** vs **Pp** competition data.

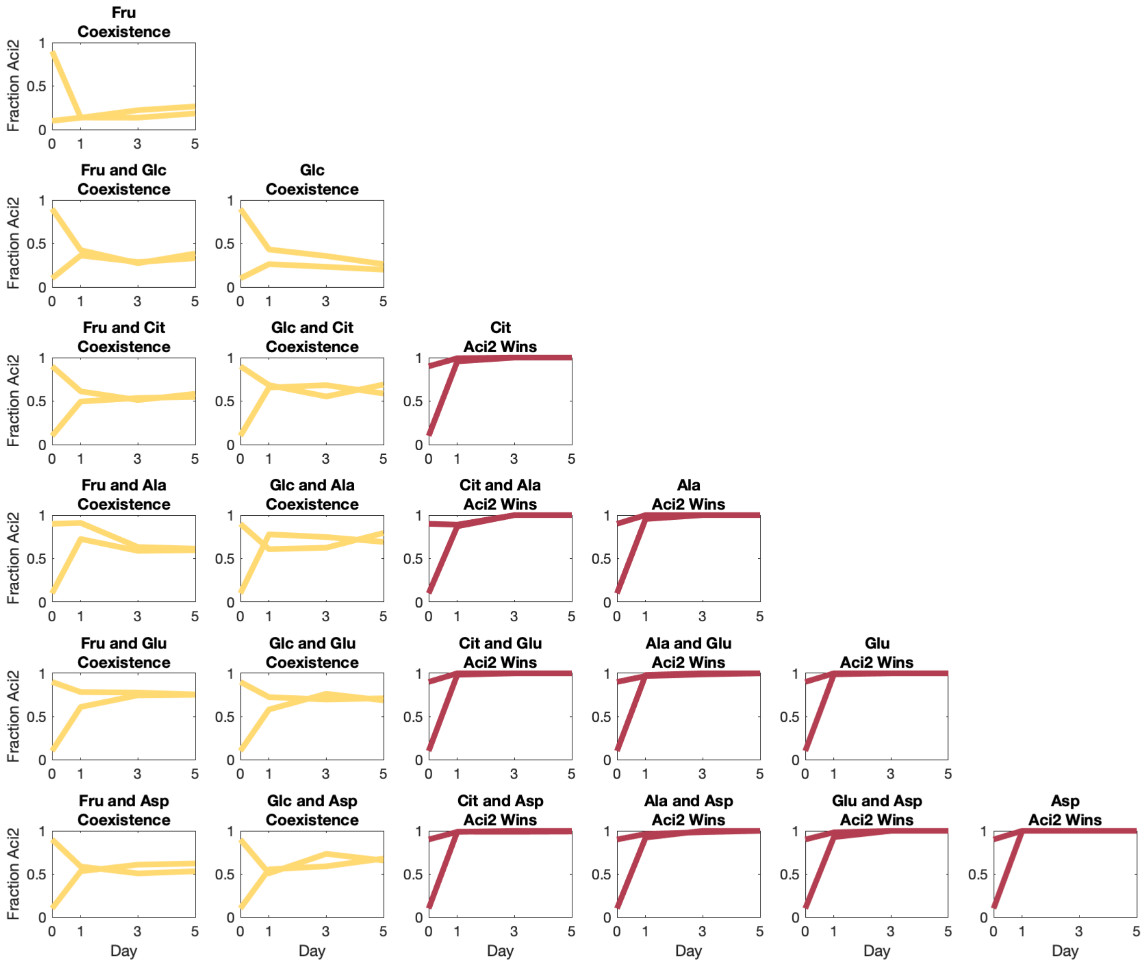


**Supp. Fig. 42** (*E*). Aci2 vs Ka competition data.

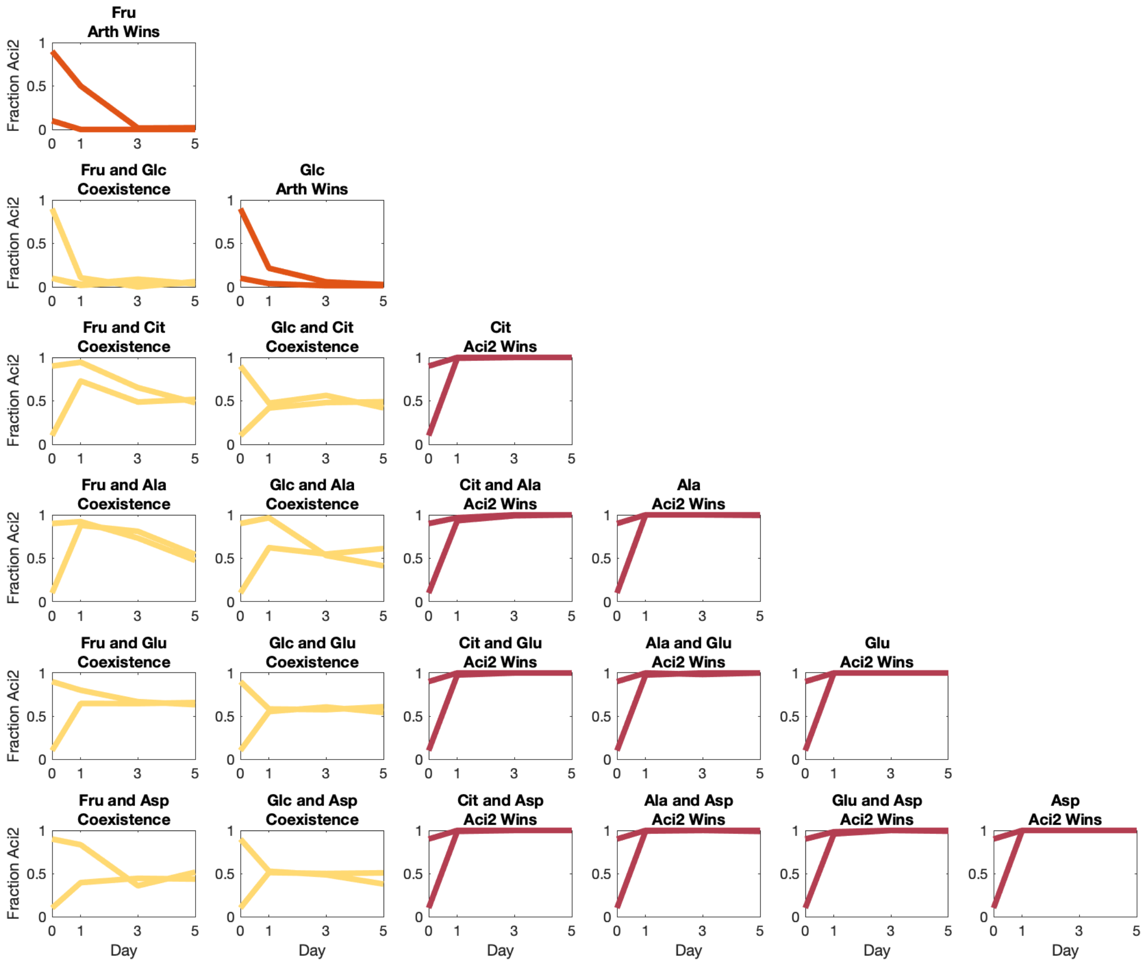


**Supp. Fig. 42** (*F*). **Aci2** vs **Arth** competition data.

On fructose and glucose, Aci2 had a Day 5 mean fraction of 4.7% +/ 0.6%. We determined this to be a case of coexistence.

On fructose, Aci2 had a Day 5 mean fraction of 1.0% +/- 1.4%, having been 0.8% +/- 1.1% on Day 3. We determined this to be a case of exclusion.

On glucose, Aci2 had a Day 5 mean fraction of 2.1% +/0.9%. We determined this to be a case of exclusion. This determination was borderline, but because this is a single-resource competition its outcome does not affect the tallies used in our calculations of the frequency of coexistence as a function of relative growth rates and lag times.

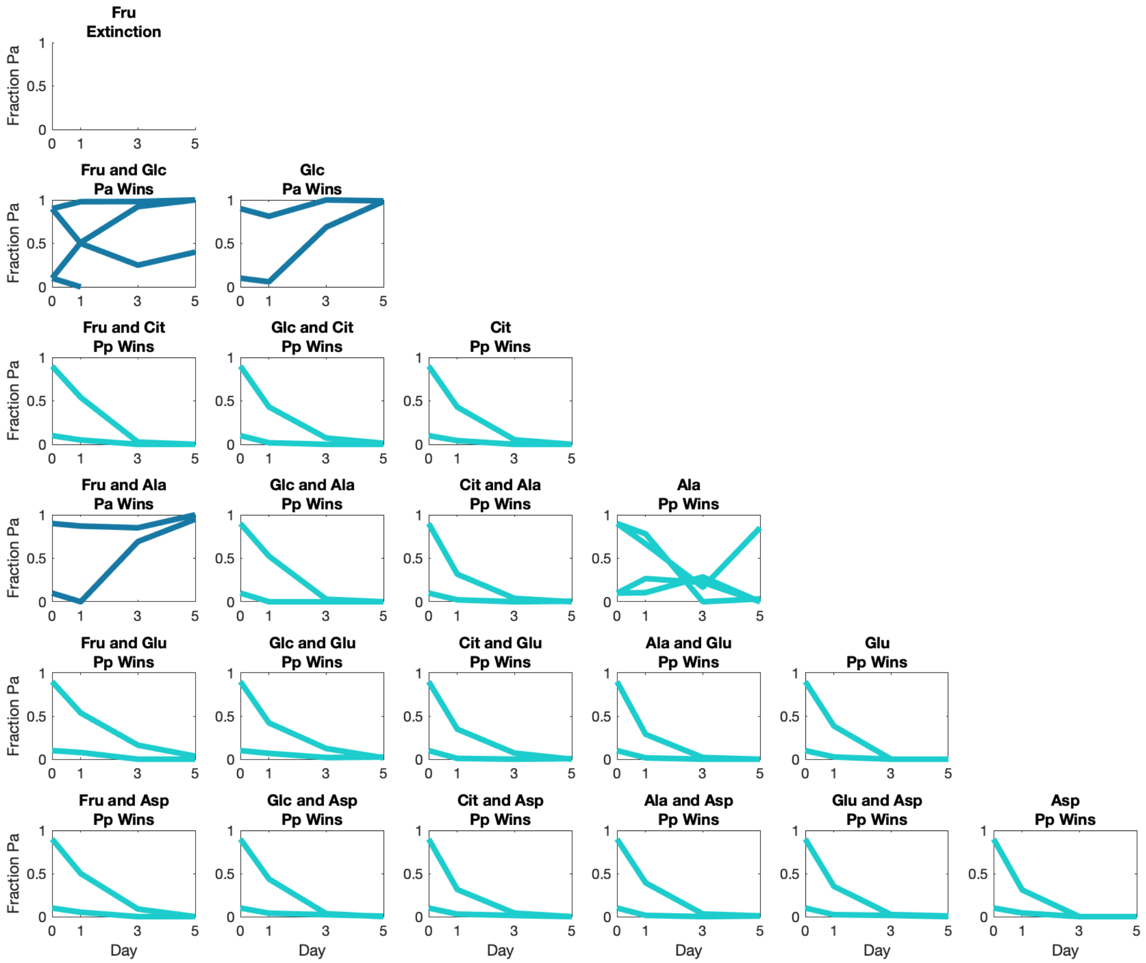


**Supp. Fig. 42** $(G)$. **Pa** vs **Pp** competition data.

On Fructose and Glucose, the species nearly went extinct in the first pair of replicates. In the second pair of replicates Pa excluded Pp while maintaining a large population fraction.

On alanine, one replicate was deemed an outlier and excluded from influencing our conclusions.

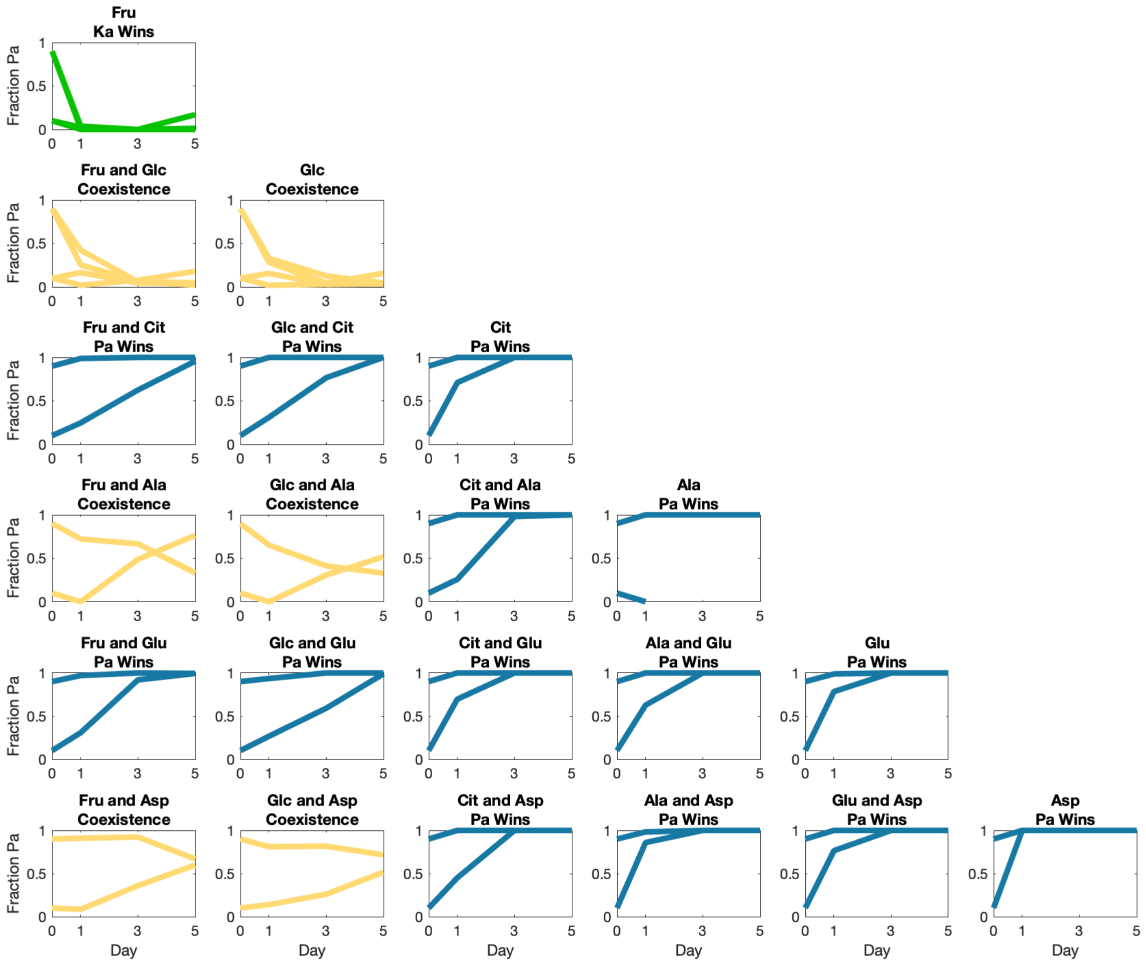


**Supp. Fig. 42** (*H*). **Pa** vs **Ka** competition data.

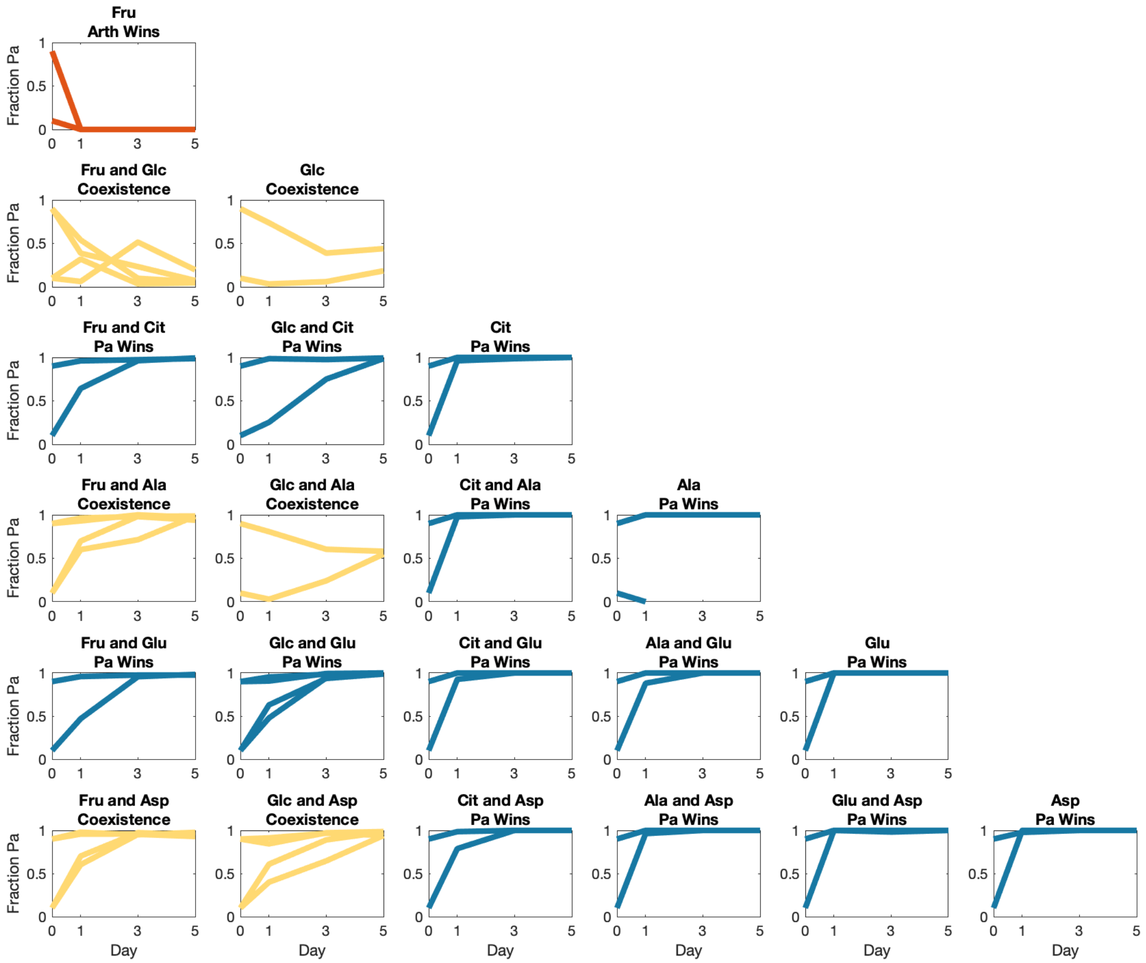


**Supp. Fig. 42** (*I*). **Pa** vs **Arth** competition data.

Excluding the visible outlier, on fructose and glucose, Pa has a mean Day 5 fraction of 3.4% +/- 0.8%. On fructose and alanine, Arth had a mean Day 5 fraction of 3.0% +/- 1.1%. On fructose and aspartate, Arth had a mean Day 5 fraction of 5.4% +/- 1.1%. On glucose and aspartate, Arth had a mean Day 5 fraction of 3.8% +/- 2.5%. We determined these to be a cases of coexistence.

On glucose and glutamate, Ka had a mean Day fraction of 0.6% +/- 0.3%. We determined this to be a case of competitive exclusion.

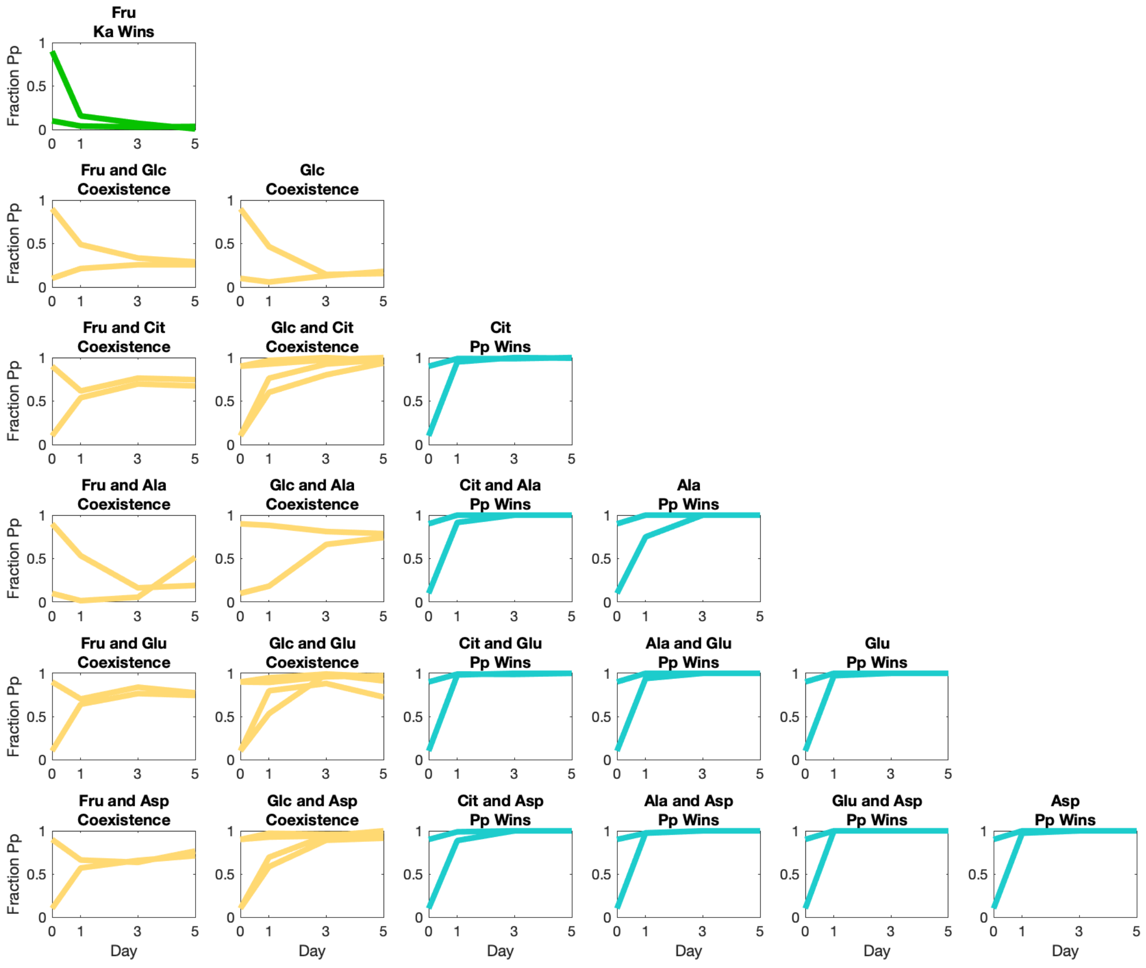


**Supp. Fig. 42** (*J*). **Pp** vs **Ka** competition data.

On glucose and citrate, Ka had a mean Day 5 population fraction of 3.2% +/- 1.3%. On glucose and aspartate, Ka had a mean Day 5 population fraction of 4.6% +/- 1.8%. We determined both of these to be cases of coexistence.

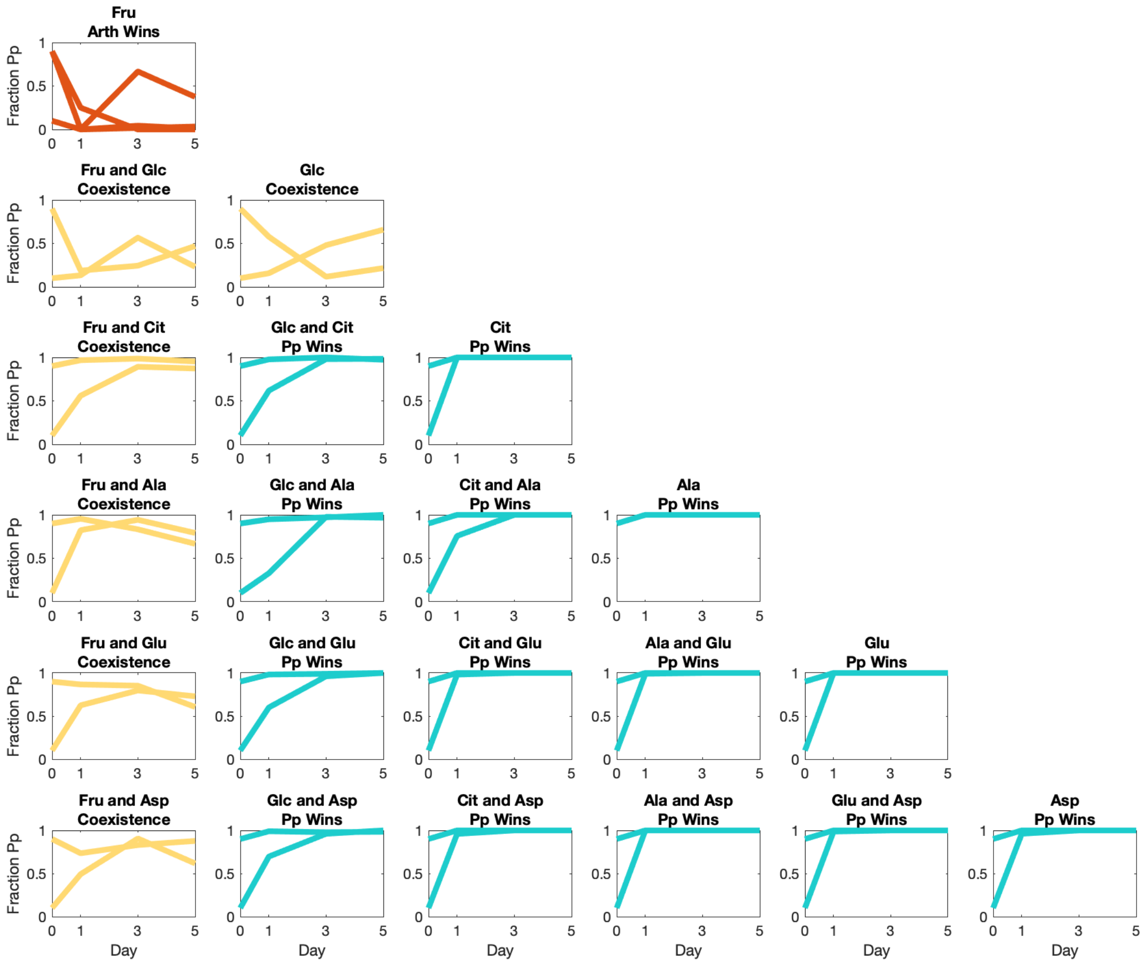


**Supp. Fig. 42** (*K*). **Pp** vs **Arth** competition data.

On fructose, excluding the one outlier, Pp had a mean Day 5 fraction of 1.5% +/- 1.0%. We determined this to be a case of exclusion. This determination was borderline, but because this is a single-resource competition its outcome does not affect the tallies used in our calculations of the frequency of coexistence as a function of relative growth rates and lag times.

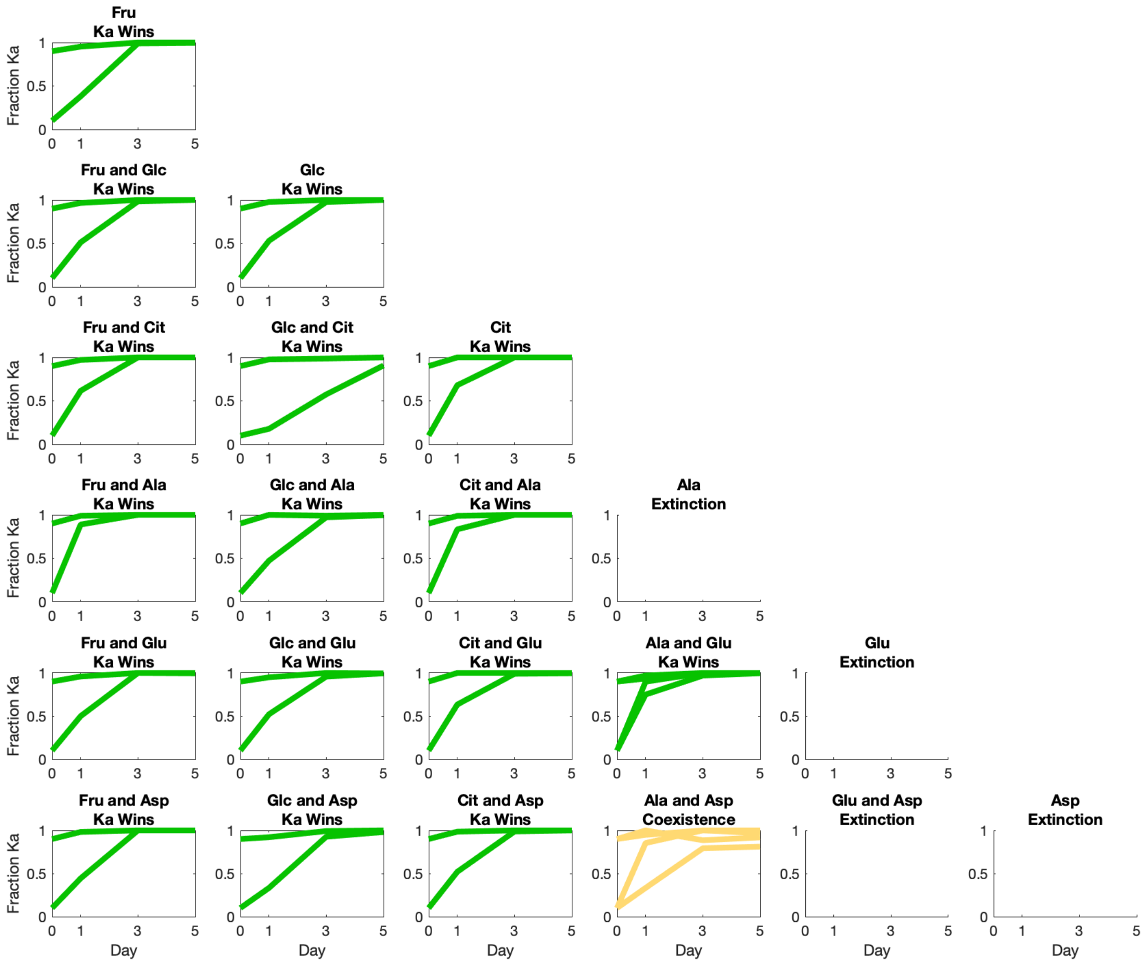


**Supp. Fig. 42** $(L)$. **Ka** vs **Arth** competition data.

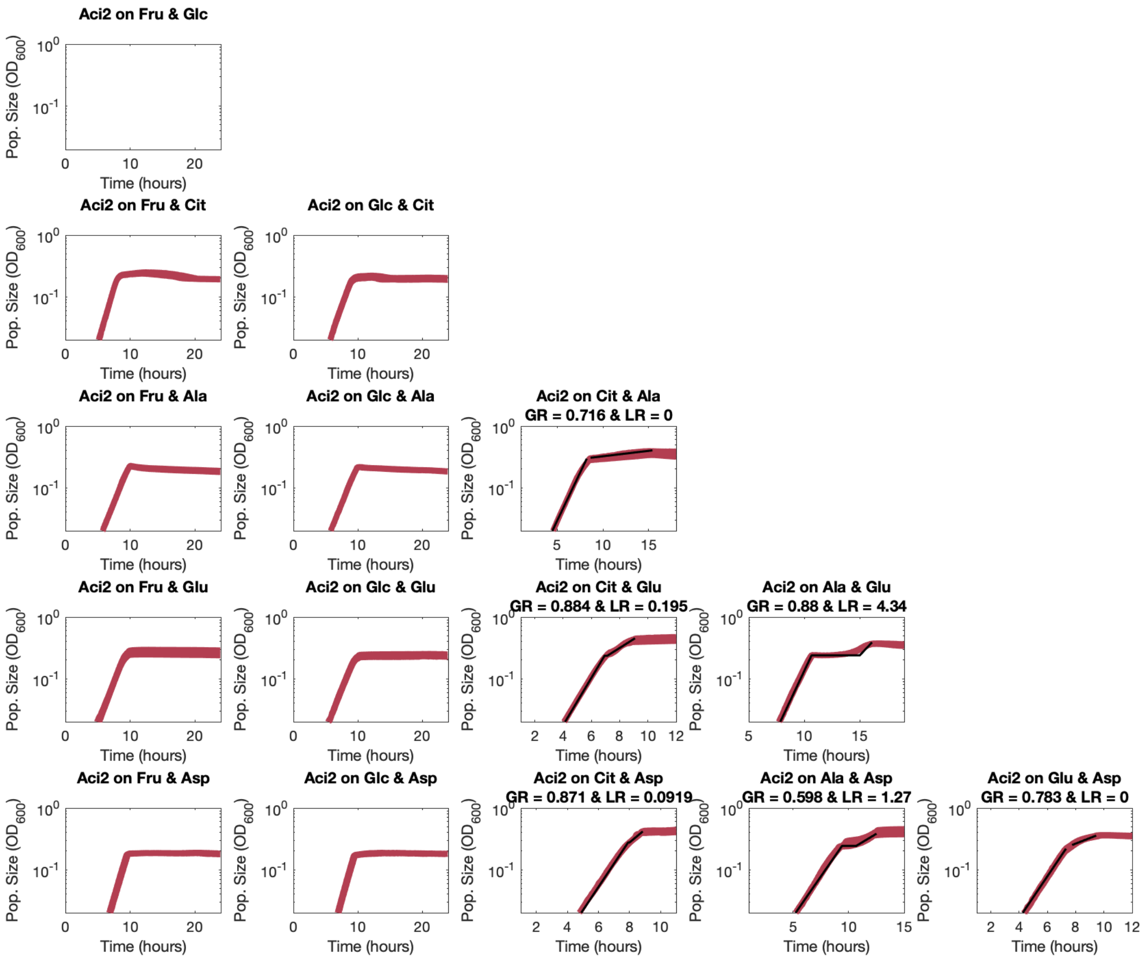


**Supp. Fig. 43** (*A*). Aci2 growth rate and lag time fits.

Aci2 did grow on fructose nor glucose.

Aci2's alanine and glutamate lag time as reported here is its lag time within a sharp recovery model, which is why it does not match the value used elsewhere This value is used here to maintain comparability to the other fits.

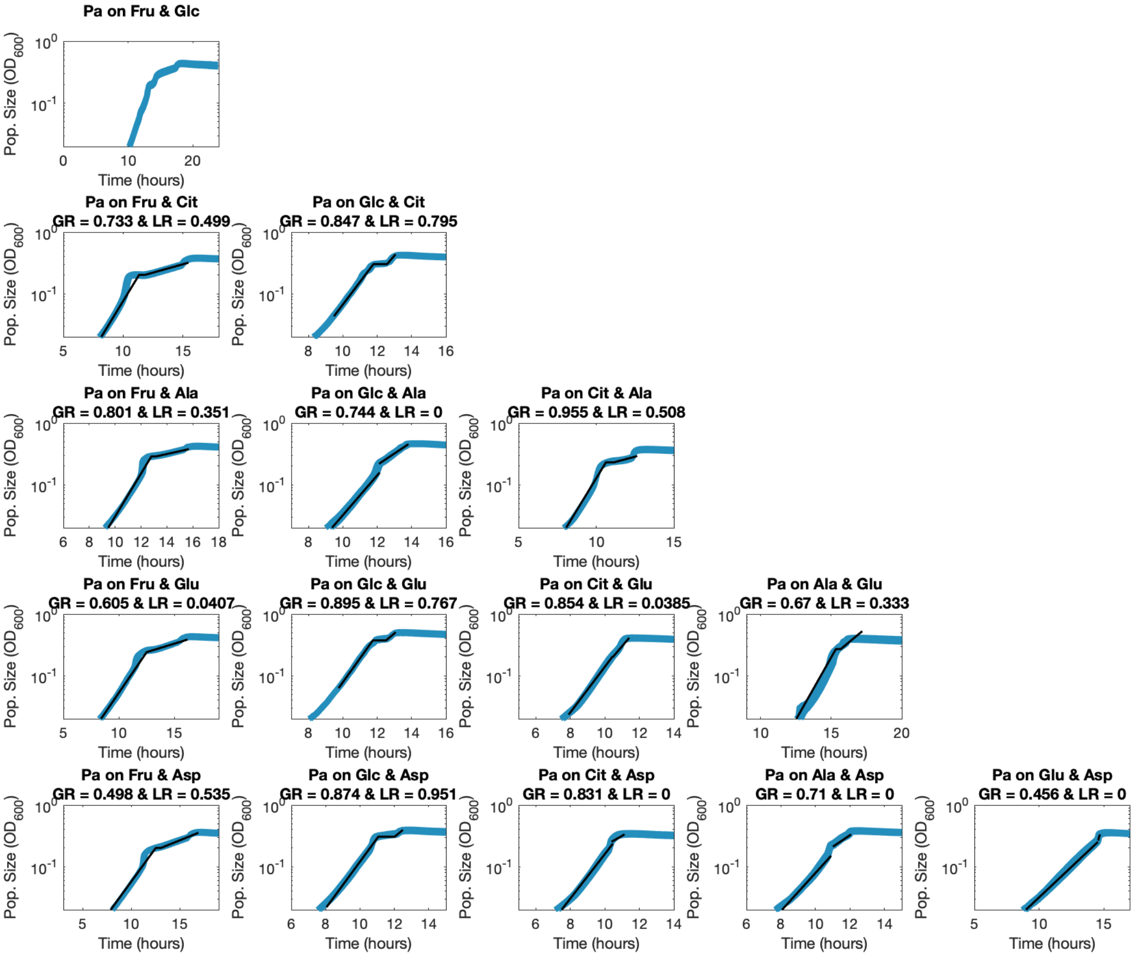


**Supp. Fig. 43** (*B*). Pa growth rate and lag time fits.

Pa had an extra diauxic shift on fructose and glucose, so growth rate and lag time were not fit.

Pa's alanine and glutamate lag time as reported here is its lag time within a sharp recovery model, which is why it does not match the value used elsewhere. This value is used here to maintain comparability to the other fits.

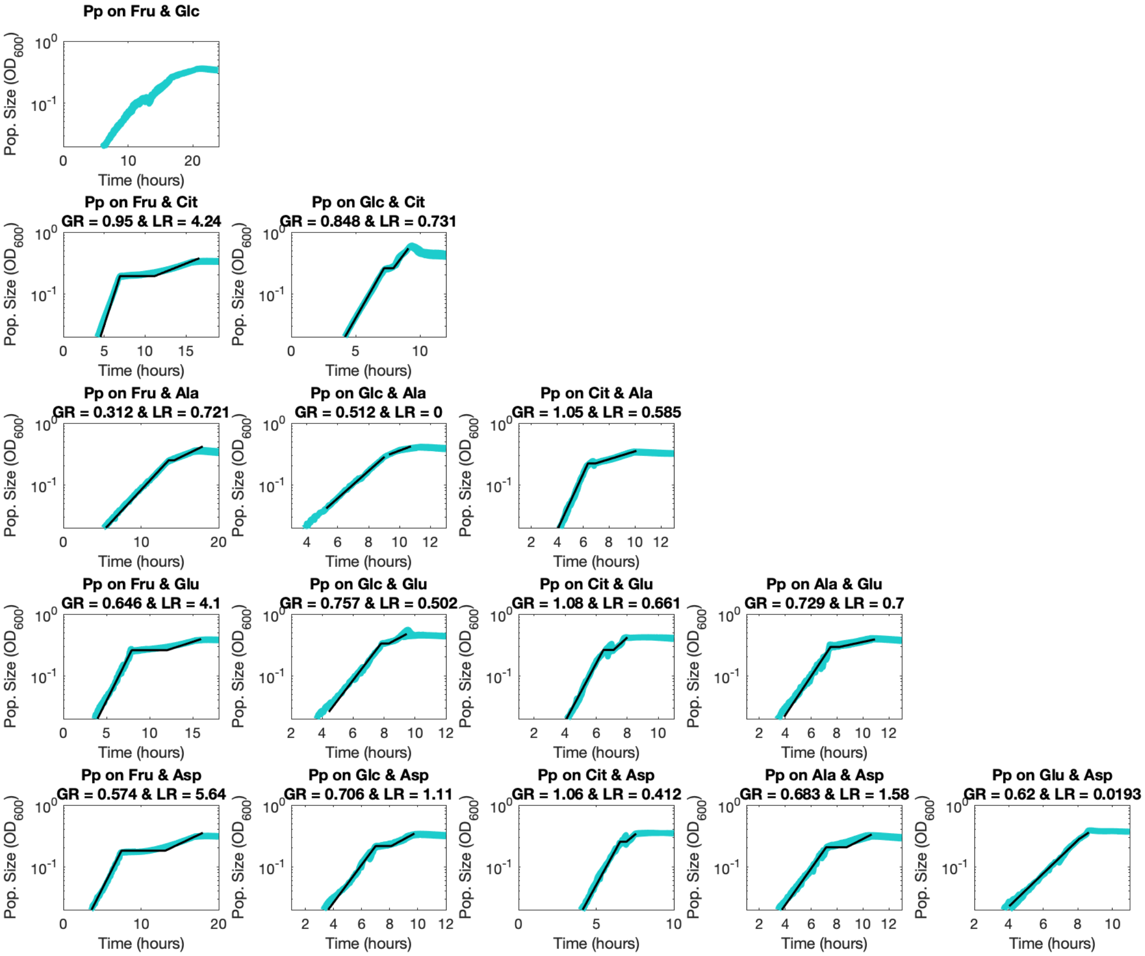


**Supp. Fig. 43** $(C)$. Pp growth rate and lag time fits.

Pp had an extra diauxic shift on fructose and glucose, so growth rate and lag time were not fit.

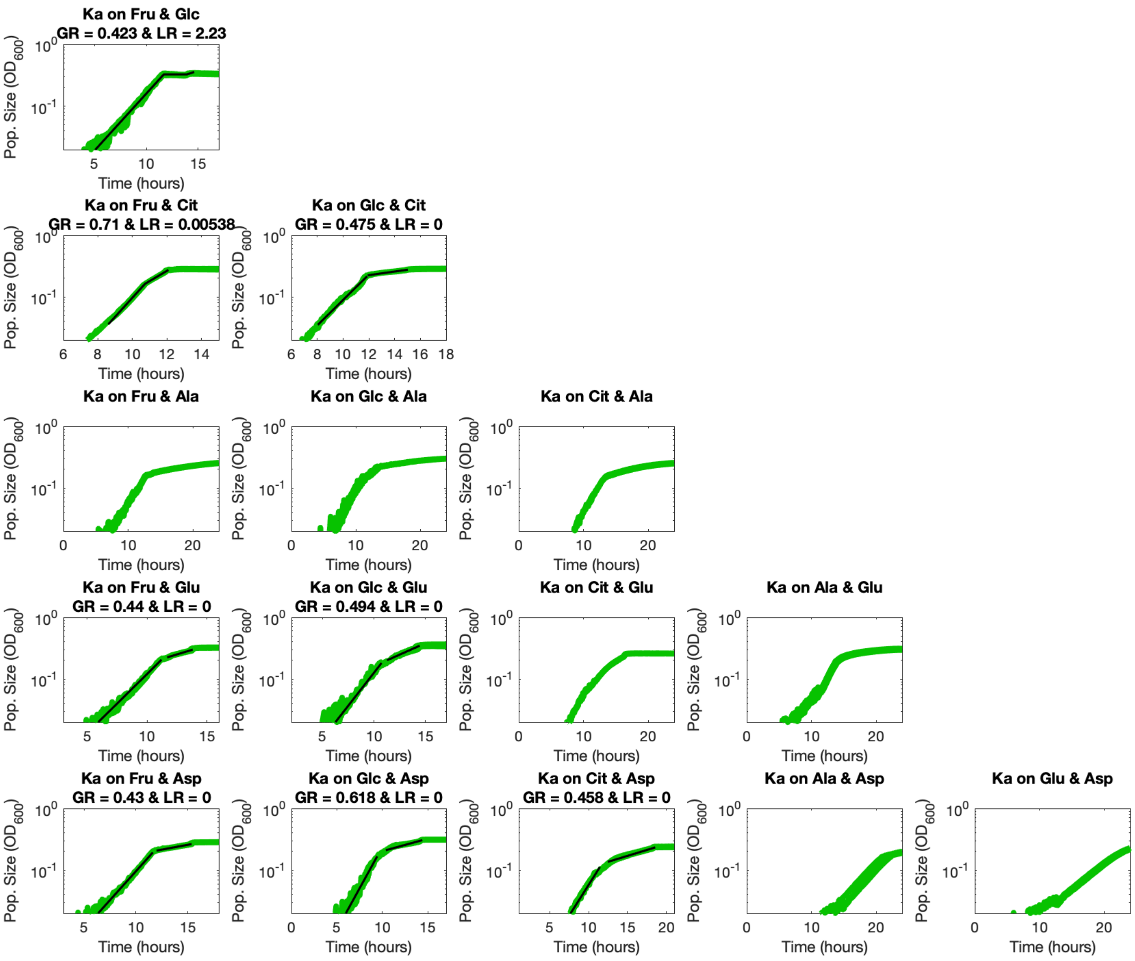


**Supp. Fig. 43** (*D*). Ka growth rate and lag time fits.

Ka did not grow on alanine.

Ka's diauxic shift on alanine and aspartate did not occur early enough for enough post-shift growth to be observed to confidently fit a lag time.

Ka's growth dynamics on citrate and glutamate featured a rounded shape to which it proved difficult to growth dynamics with sufficient confidence.

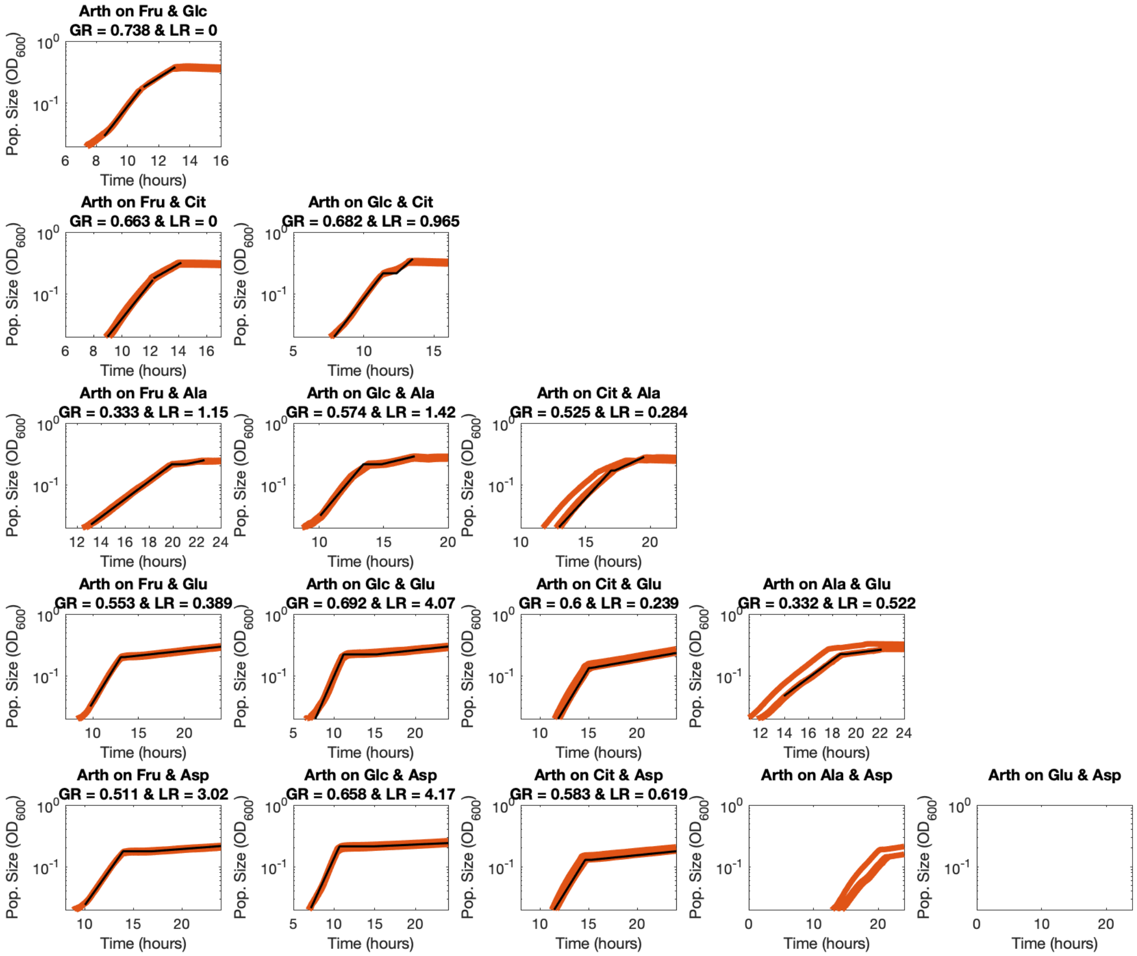


**Supp. Fig. 43** (*E*). Arth growth rate and lag time fits.

Arth diauxic shift on alanine and aspartate did not occur early enough for enough post-shift growth to be observed to confidently fit a lag time.

Arth grew slightly on the combination of glutamate and aspartate but not enough for its optical density to sufficiently exceed our background noise.
